# A qualitative review of Oxford Nanopore Sequencing datasets for RNA modifications

**DOI:** 10.1101/2024.09.26.615132

**Authors:** Madhurananda Pahar, Qian Liu

## Abstract

There are many oxford nanopore datasets available to study methylations. Methylations and modifications occur at nucleotides such as adenine (A), cytosine (C), guanine (G) and theanine (T) or uracil (U). Among all these provided datasets, some have the most common m6A methylation and others have m5C etc. using various real organism reference sequences such as human, mouse and artificial reference sequences which are prepared in the laboratory such as curlcake and IVT. These datasets are required to be organized by the methylation types to research ONT datasets. Here we provide a summary of the read qualities, base mapping success rates etc. for these methylation types and reference genomes. We have used minimap2 base mapping and longreadsum results. We find that methylated data have lower success rates than non-methylated data and mapping quality is lower for the real reference genomes such as human and mice. This could be because they contain more than 100,000 transcriptomes whereas artificial reference sequences contain only a few transcriptomes. Datasets which contain artificially created reference sequences have a higher quality than the others, thus they are recommended to be used for methylation or modification classification tasks in the future.

**Dataset:** All datasets used in this study are the publicly available.

**Dataset License:** All datasets used in this study are the publicly available.

## 1. Introduction

A well-functioning body requires proper instructions at the right time. Such instructions including to build and maintain cells are coded inside deoxyribonucleic acid (DNA) in the form of four nucleotides which are Adenine (A), Thymine (T), Cytosine (C) and Guanine (G) and they form base pairs to deliver the instructions. To understand these instructions, DNA must be copied into ribonucleic acid (RNA) and this process is called transcription. The readouts are called transcripts and the collection of all such readouts is called a transcriptome [1]. RNA sequencing (RNA-seq) is vital for the study of the transcriptome, which helps researchers to better understand cell types, cell functions and activities which may contribute to diseases [2]. Thus RNA sequencing replicates the sequence of the DNA and a gene can be turned on or off at a specific time and place in the cell and tissues of an organism by analyzing the entire collection of RNA sequences in that cell i.e. the transcriptome [3]. RNA-seq is a highly sensitive and accurate tool for measuring expression across the transcriptome, thus identifying features such as transcript isoforms, single nucleotide variants, and gene fusions [4].

RNA modifications have become popular due to their ability to influence several RNA processes like transportation, generation, metabolization and function thus becoming critical regulators of cell biology. Recent studies have revealed that RNA modifications involve multiple biological processes including migration, activation, differentiation, polarization and some immune-related diseases [5]. RNA modifications are the mechanisms to alter genetic information and recently more than 130 different RNA modifications have been discovered. Mapping selected RNA modifications at single-nucleotide resolutions is complex but has been successfully implemented to increase the knowledge of the biological function of RNA modifications, which is the study of epitranscriptome [6]. The study of epitranscriptome has been advanced to clinical usage over the past few years. Recent studies also provide evidence for the fact that the way RNA modifications regulate basic cell functions in higher organisms can also help to understand how RNA modifications can cause disease in humans [7].

To date, over 150 different types of RNA modifications have been identified across all species and studies have revealed that they play a pivotal role in many cellular processes. Among them, N6-methyladenosine (m^6^A) which is the most frequent chemical modification in messenger RNA also participates in various physiological processes, such as viral infection and cancer progression. m^6^A methyltransferase and m^6^A demethylase enzymes act as the writer and eraser constituting the dynamic and reversible m^6^A modification. These writers and erasers play a vital role in determining the abundance of m^6^A modifications and their function and distribution [8]. Thus, m^6^A is an important regulator of gene expression. Transcriptome-wide m^6^A detection has primarily been achieved using the next-generation sequencing (NGS) platform. There exists a fundamental challenge for mapping diverse RNA modification types simultaneously. Moreover, NGS-based methods are often not quantitative and contain high false-positive rates. They are also inconsistent when using distinct antibodies, and are unable to produce maps for highly repetitive regions. They are also unable to provide information regarding the co-occurrence of distant modifications in the same transcripts and require multiple ligation steps and extensive polymerase chain reaction amplification during the library preparation, introducing undesired biases in the sequencing data [9].

Recently, direct RNA sequencing (DRS) using Oxford Nanopore Technologies (ONT) has also demonstrated a promising alternative option as it has the potential to detect virtually any given RNA modification present in native RNA molecules [10]. Nanopore sequencing is a process of investigating nucleic acids and other biomacromolecules where a nano-scale hole is embedded, and the electrochemical signal is measured. Oxford Nanopore Technology (ONT) has been the most successful application of nanopore technology and marked the beginning of the fourth generation of gene sequencing technology [11]. These rapid advances in nanopore technologies for sequencing single-long DNA and RNA molecules have led to substantial improvements in accuracy, read length and throughput. This also required extensive development of experimental and bioinformatics methods to fully explore nanopore long reads to investigate genomes, transcriptomes, epigenomes and epitranscriptomes [12]. Unlike short reads where retrieving RNA modifications per nucleotide can be challenging, ONT is a long-read sequencing technique [13]. Unlike the Pacific Bioscience (PacBio) platform, where the sequencing requires a high coverage but suffers from low discriminatory power; ONT performs better in mapping nucleotide modifications as it uses direct sequencing for DNA or RNA. The flow of current is different for the modified nucleotides than those that are unmodified and those differences help detect the modifications such as m^6^A. Raw ONT signal is required to pass through the alignment process which is called base-calling, which identifies a nucleotide using a base caller called guppy which uses a bi-directional recurrent neural network.

Statistical and machine learning-based tools have already shown promise in detecting m^6^A modifications while using ONT signals and both current and alignment features [14]. Currents from the nanopore are assigned to each nucleotide or event using their features such as mean, median, standard deviation etc. For example, Tombo [15] uses statistical tests, whereas MINES [16], Nanom6A [17] and m6Anet [18] use machine learning on current features. Nanocompore [19] and xPore [20] also use clustering on current features. DiffErr [21], DRUMMER [22] and ELIGOS [23] use statistical tests, whereas Epinano [24] uses machine learning on the alignment feature resulting from the base-calling errors [10]. Recently, innovative machine-learning techniques such as recurrent neural networks have been applied to long-read Oxford Nanopore sequencing data to detect modifications in DNA [24], [25]. Similar techniques have also been found useful in reducing base-calling errors in nanopore sequencing signals [26]. Currently, we also adopted a convolutional neural network trained and evaluated on traditional signal processing features extracted from long-read ONT signals to detect the modified nucleotides in RNA sequences. This required us carrying out a thorough investigation of the publicly available Nanopore sequencing data which can be used in modification or methylation classification. We found seven of such sources and they are:

- Pratanwanich et al. [20]
- Mateos et al. [27]
- Liu et al. [24]
- Begik et al. [9]
- Jenjaroenpun et al. [28]
- Zhong et al. [10]
- Stephenson et al. [29]

These sources represent eight different reference transcriptomes, and they are:

- Human (Homo sapiens)
- Mouse (Mus musculus)
- Curlcake for m6A methylation ([24])
- Curlcake for pseudouridine methylation ([9])
- In vitro test (IVT)
- Yeast (Saccharomyces cerevisiae)
- Escherichia Coli (E Coli)
- Arabidopsis Thaliana

In this study, we first describe the methods used to calculate the analytics in the next section. Then, we start explaining the dataset according to various methylation types, reference sequences and sources discussed in the previous paragraph. Next, the results are discussed, and the following research questions were answered:

- What are some of the existing methylated/modified datasets available today?
- What is the size of those data and how many reads and bases are there?
- How well do those data align with their corresponding reference transcriptomes?
- How do they compare to each other, and which dataset is more suitable to be used for modification/methylation detection?

## 2. Methods

We have downloaded Oxford nanopore data from seven different papers. They are as follows: The data download pipeline has been shown in the following figure, After downloading the data, we pre-process the data (download using wget, unzip the corpus, multiple read to single read, delete the existing analyses present), basecalling, carry out BAM analysis [30], longreadsum and finally do Tombo resquiggle and check mapping quality. The codes for this data analysis is presented in this <u>GitHub link</u>.

Results are described using a table showing the dataset name, type, size in gigabytes (GB), number of reads and bases, success rate of base mapping, Tombo and pass rates.

Here in the tables, size is the size of FAST5 files after deleting the previous analyses. Base mapping success rate is after using minimap2 for base mapping. The bam.stat file for HEK293T-KD-Ctrl-rep1 produces the following information:

### 2.1. Base-mapping success rate

Thus the base mapping success rate is 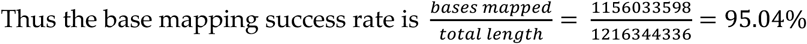

### 2.2. Tombo percents

Similarly, 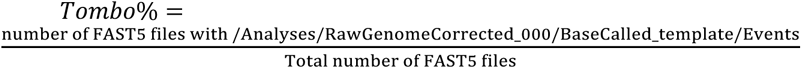

### 2.3. Pass rates

Also, 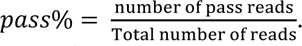. The number of pass reads are calculated by dividing the number of lines in the fastq files in the ‘pass’ folder after base-calling by 4, as each fastq file contains 4 lines of information (sequence identifier, nucleotide sequence, strand, basecall quality scores) for each read.

We also show the length of the aligned query sequence, mapping quality and transcripts. The threshold of the query align length was set at 200 [31] and the threshold of the mapping quality was set at 20 [32].

## 3. Datasets from various methylation types

All figures and tables should be cited in the main text as Figure 1, Table 1, etc.

**Figure 1:**
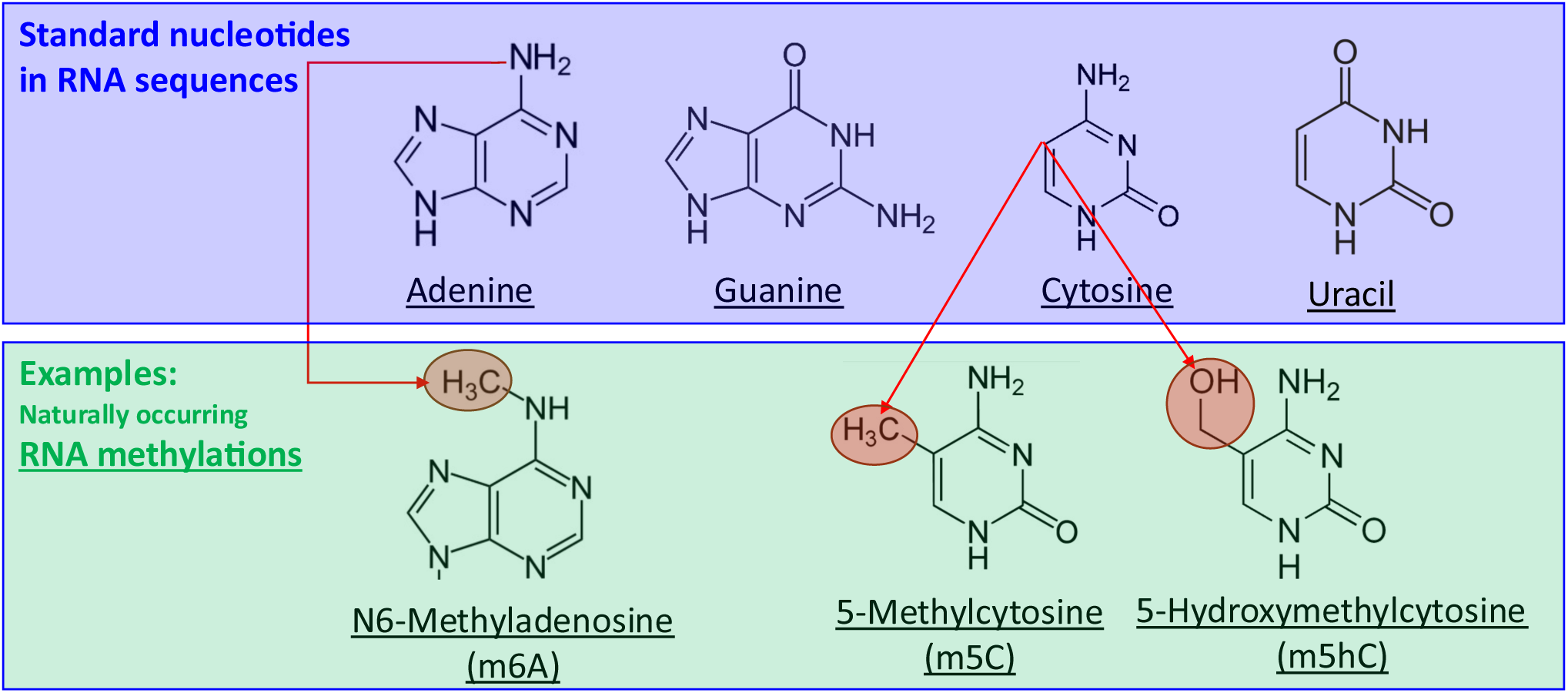
Examples of RNA methylations such as m6A, m5C and m5hC. Here, the standard nucleotides are shown at the top and then naturally occurring RNA methylations are shown at the bottom.

**Table 1.** This table explains the datasets for human and m6A methylation and shows size (in GB), the number of reads and bases, corresponding references and base mapping, Tombo success rates and pass rates (see Section 2)

| Datasets | Type | Size (GB) | #READ | #BASE | Reference | Base map% | Tombo % | Pass % |
| --- | --- | --- | --- | --- | --- | --- | --- | --- |
| HEK293T-KD-Ctrl-rep1 | KD-Ctrl | 110 | 1,233,669 | 1,216,344,336 | [20] | 95.04 | 93.34 | 99.96 |
| HEK293T-KD-Ctrl-rep2 | KD-Ctrl | 85 | 1001619 | 963,174,641 | [20] | 97.44 | 96.27 | 99.97 |
| HEK293T-KD-Ctrl-rep3 | KD-Ctrl | 113 | 1247684 | 1,253,510,129 | [20] | 97.58 | 96.26 | 99.98 |
| HEK293T-KD-rep1 | KD-Ctrl | 160 | 1890698 | 1,778,988,763 | [20] | 95.91 | 94.26 | 99.96 |
| HEK293T-KD-rep2 | KD-Ctrl | 105 | 1,209,898 | 1,156,680,216 | [20] | 97.86 | 96.41 | 99.98 |
| HEK293T-KD-rep3 | KD-Ctrl | 60 | 721,199 | 669,875,292 | [20] | 97.49 | 96.45 | 99.98 |
| HEK293T-Mettl3-KO-rep1 | Mettl3-KO | 126 | 1,490,210 | 1,421,483,252 | [20] | 96.80 | 93.05 | 94.18 |
| HEK293T-Mettl3-KO-rep2 | Mettl3-KO | 139 | 1,815,589 | 1,806,479,910 | [20] | 93.85 | 90.53 | 93.66 |
| HEK293T-Mettl3-KO-rep3 | Mettl3-KO | 122 | 1,677,075 | 1,629,147,850 | [20] | 94.14 | 91.16 | 94.23 |
| HEK293T-WT-0-rep1 | WT | 125 | 1,353,473 | 1,456,531,193 | [20] | 98.30 | 95.63 | 99.97 |
| HEK293T-WT-0-rep2 | WT | 113 | 1,286,100 | 1,259,966,253 | [20] | 97.42 | 94.95 | 99.97 |
| HEK293T-WT-0-rep3 | WT | 161 | 1,730,587 | 1,862,604,111 | [20] | 98.29 | 95.87 | 99.98 |
| HEK293T-WT-100-rep1 | WT | 84 | 931,436 | 948,388,596 | [20] | 97.91 | 94.96 | 99.95 |
| HEK293T-WT-100-rep2 | WT | 93 | 1,001,759 | 1,079,808,526 | [20] | 97.02 | 94.22 | 99.98 |
| HEK293T-WT-100-rep3 | WT | 174 | 1,850,033 | 1,938,165,278 | [20] | 98.02 | 95.42 | 99.97 |
| HEK293T-WT-25-rep1 | WT | 91 | 983,685 | 1,008,823,857 | [20] | 97.91 | 94.87 | 99.96 |
| HEK293T-WT-25-rep2 | WT | 121 | 1,501,189 | 1,582,964,558 | [20] | 91.13 | 86.43 | 89.09 |
| HEK293T-WT-25-rep3 | WT | 132 | 1,414,005 | 1,487,033,409 | [20] | 97.83 | 95.38 | 99.98 |
| HEK293T-WT-25-rep4 | WT | 121 | 1,501,189 | 1,582,964,558 | [20] | 91.13 | 86.43 | 89.09 |
| HEK293T-WT-50-rep1 | WT | 97 | 1,028,780 | 1,105,111,672 | [20] | 97.69 | 94.98 | 99.97 |
| HEK293T-WT-50-rep2 | WT | 117 | 1,254,029 | 1,304,216,171 | [20] | 97.01 | 94.41 | 99.98 |
| HEK293T-WT-50-rep3 | WT | 61 | 779,711 | 797,807,374 | [20] | 92.70 | 87.21 | 88.34 |
| HEK293T-WT-50-rep4 | WT | 61 | 779,711 | 797,807,374 | [20] | 92.70 | 87.21 | 88.34 |
| HEK293T-WT-75-rep1 | WT | 139 | 1,469,497 | 1,534,912,329 | [20] | 97.48 | 94.62 | 99.97 |
| HEK293T-WT-75-rep2 | WT | 247 | 2,658,577 | 2,699,149,633 | [20] | 97.67 | 94.95 | 99.97 |
| HEK293T-WT-75-rep3 | WT | 247 | 2,658,577 | 2,699,149,633 | [20] | 97.67 | 94.95 | 99.97 |
| HEK293T-WT-75-rep4 | WT | 89 | 1,124,706 | 1,135,182,896 | [20] | 92.15 | 86.02 | 86.70 |
| HEK293T-WT-rep1 | WT | 89 | 1,040,661 | 979,367,769 | [20] | 96.53 | 93.16 | 94.61 |
| HEK293T-WT-rep2 | WT | 111 | 1,396,000 | 1,505,941,085 | [20] | 93.11 | 89.29 | 92.71 |
| HEK293T-WT-rep3 | WT | 42 | 513,670 | 516,871,680 | [20] | 91.75 | 86.81 | 89.73 |
| Myeloma-rep1 | Myeloma | 30 | 368,440 | 375,131,307 | [20] | 72.80 | 71.16 | 92.96 |
| Myeloma-rep2 | Myeloma | 84 | 914,696 | 1,015,493,343 | [20] | 77.82 | 74.86 | 95.30 |
| Myeloma-rep3 | Myeloma | 47 | 601,894 | 587,630,567 | [20] | 68.65 | 69.29 | 93.44 |
| HS_H460_fast5 | NativeRNA | 150 | 1,788,451 | 2,021,194,598 | [28] | 96.65 | 92.77 | 92.48 |
| HS_rRNA_dRNA_fast5 | NativeRNA | 5.2 | 49,310 | 63,231,961 | [28] | 93.19 | 88.05 | 89.21 |
| HS_SAEC_fast5 | NativeRNA | 151 | 1,894,424 | 2,008,298,396 | [28] | 97.16 | 93.83 | 93.82 |
| IVT_Hs_mRNA | NativeRNA | 124 | 2,600,523 | 1,585,075,558 | [28] | 94.06 | 91.2 | 93.96 |

### 3.1. m6A methylation types

The following datasets describe the m6A methylation types

#### 3.1.1. Human

The above figure shows that more than 85% of reads and bases are successfully mapped for all the other datasets except Myeloma, as it drops down to 65% to 80%.

**Figure 2.**
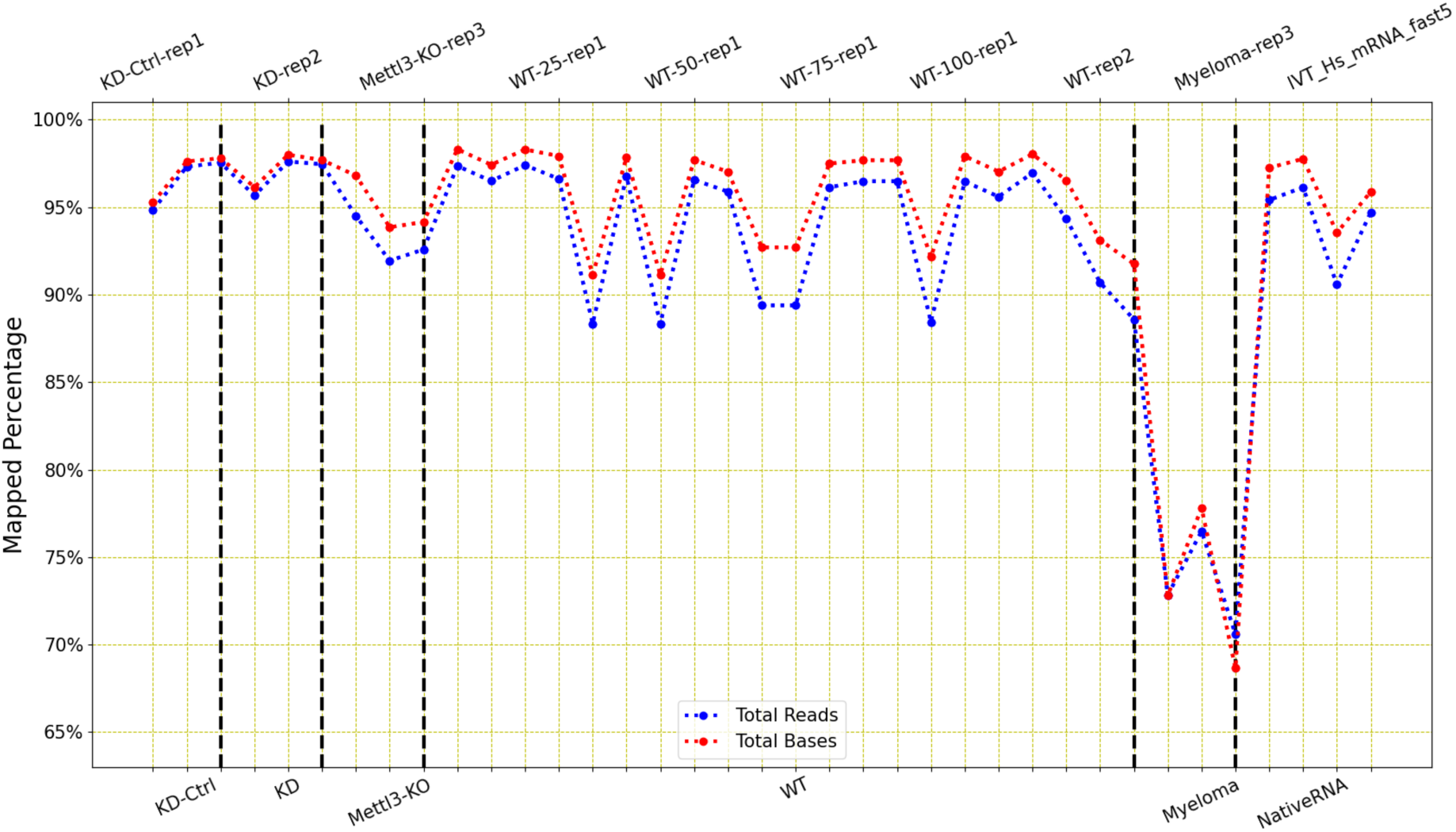
LongReadSum mapping percentage of reads and bases of Human m6A methylation data: This figure shows that even though there are variations present in these dataset types, Myeloma has the least mapping success rate for both reads and bases among all others.

The ONT data for the m6A methylation for human genome originates in Pratanwanich et al. [20] and Jenjaroenpun et al. [28]. Pratanwanich et al. [20] contains 33 different datasets, 3 of which are ‘KD-Ctrl’, 3 are ‘KD’, 3 are ‘Mettl3-KO’, 21 are ‘WT’, 3 are Myeloma. Jenjaroenpun et al. [28] contains only 4 ‘Native RNA’ datasets. These six types of datasets are explained in the figure below.

**Figure 3.**
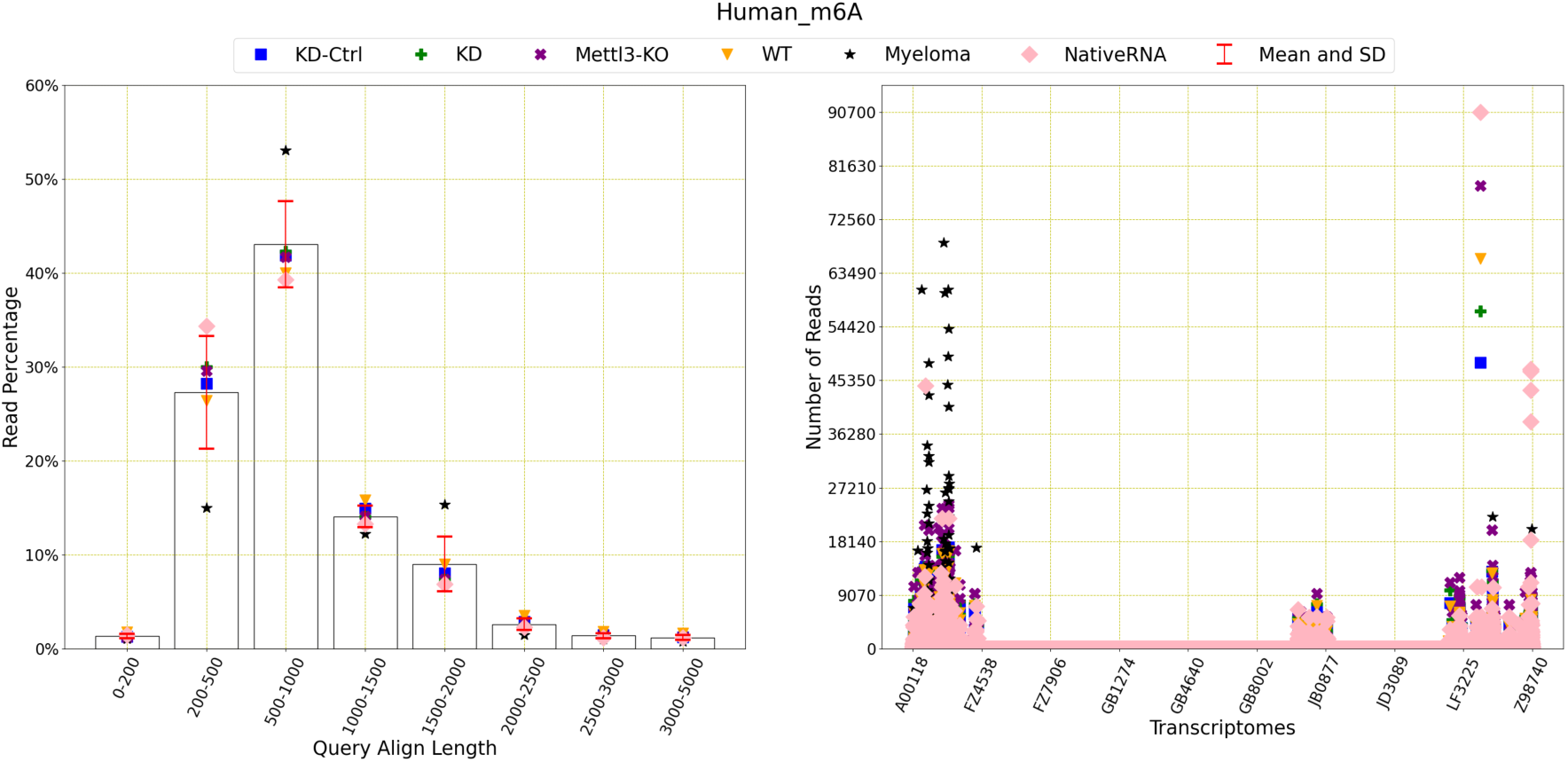
Read distribution of Human m6A methylation data: This figure shows the distribution of the reads for query align length, mapping quality and transcriptomes. As human transcriptome contains 3001485 transcripts, only 10 of them are shown on the x-axis. Mettl3-KO outperforms the others in terms of query align lengths and mapping quality and Myeloma performs the worst. The average number of reads with query length under 200 is 89,813 and over 200 is 6,263,805. This means 98.59% of the reads have a query length of greater than 200. The average number of reads of mapping quality less than 20 is 6,318,666 and over 20 is 122,849. This shows that only 1.91% of reads have a mapping quality higher than 20.

The above figure and table show that even though all these datasets perform reasonably well, there are some variations among the quality of those datasets. Mettle-KO has the highest base-mapping success rate, Tombo align success rate and pass rate. The query align lengths and mapping qualities of the reads present in Mettl3-KO are longer and better than the others as well.

Based on these two figures, we recommend using all datasets except Myeloma.

#### 3.1.2. Mouse

The ONT datasets for m6A methylation using mouse reference sequence is collected from Jenjaroenpun et al. [28] and Zhong et al. [10].

**Table 2.** This table explains the datasets for mouse and m6A methylation and shows size (in GB), the number of reads and bases, corresponding references and base mapping, Tombo success rates and pass rates (see Section 2)

| Datasets | Size | #READ | #BASE | Ref | Base map% | Tombo % | Pass % |
| --- | --- | --- | --- | --- | --- | --- | --- |
| mESCs_Mettl14_KO-ref_fast5 | 90 | 1,810,851 | 948,483,606 | [28] | 98.66 | 96.78 | 97.15 |
| mESCs_Mettl14_WT_fast5 | 76 | 1,589,429 | 739,869,336 | [28] | 98.57 | 96.67 | 97.05 |
| mESCs_Mettl3_KO-ref_fast5 | 70 | 1,527,561 | 669,899,409 | [28] | 98.56 | 96.7 | 97.28 |
| mESCs_Mettl3_WT_fast5 | 162 | 3,163,286 | 1,667,549,503 | [28] | 98.79 | 97.02 | 97.31 |
| mES_KO | 115 | 1,317,434 | 1,413,447,113 | [10] | 99.46 | 91.31 | 99.97 |
| mES_WT | 109 | 1,310,537 | 1,387,765,383 | [10] | 99.50 | 90.57 | 99.98 |

The base mapping percentage, Tombo and pass rates are high for all six datasets.

**Figure 4.**
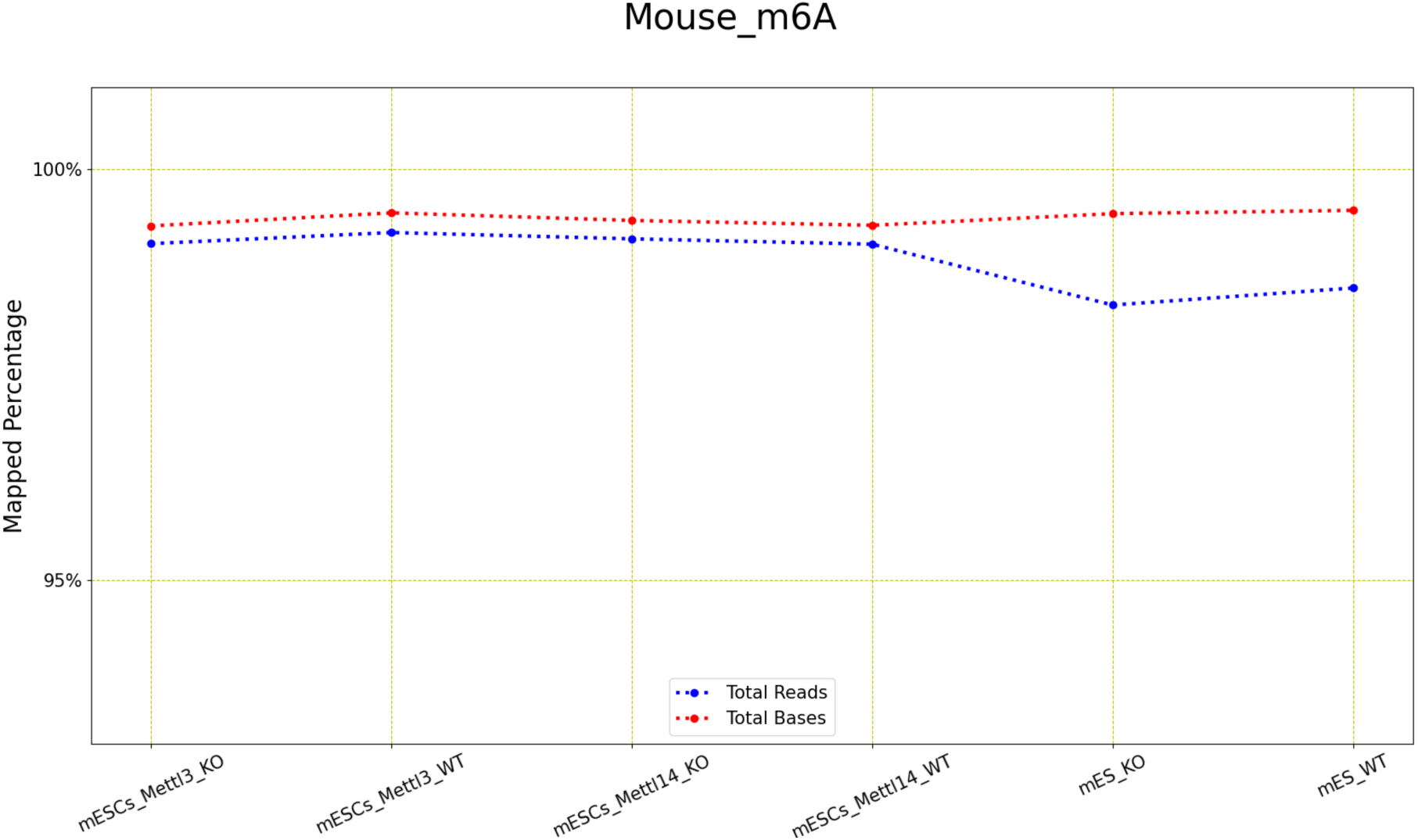
LongReadSum mapping percentage of reads and bases of Mouse m6A methylation data: This figure shows that all six datasets have a high (over 95%) mapping success rate for both reads and bases.

The above figure shows very little variance between the read and base mapping percentages.

There are altogether six ONT datasets for m6A methylation and mouse genome and four of them are from Jenjaroenpun et al. [28] and two are from Zhong et al. [10]. Their read distribution has shown in the following figure.

**Figure 5.**
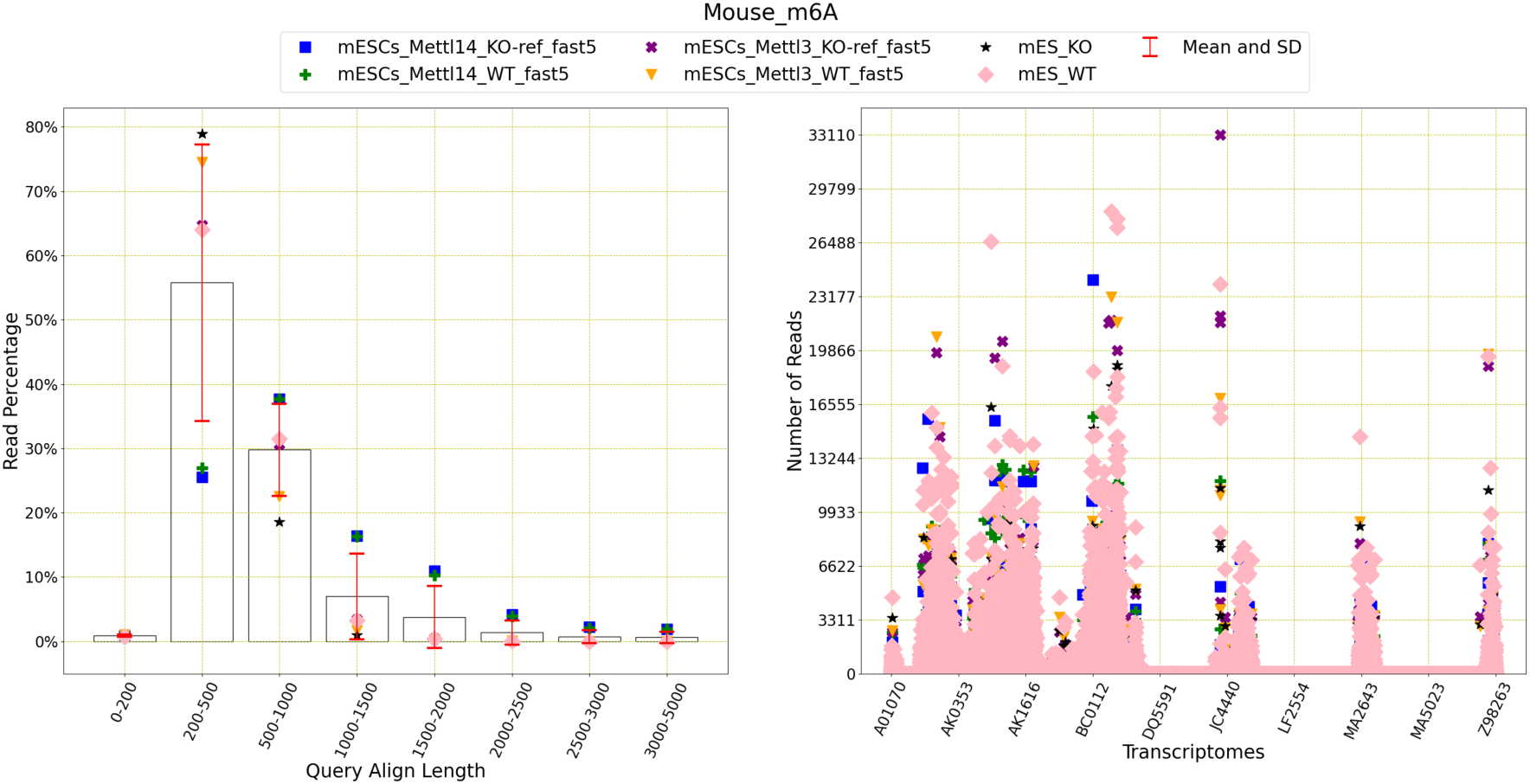
Read distribution of Mouse m6A methylation data: This figure shows the distribution of the reads for query align length, mapping quality and transcriptomes. As mouse transcriptome contains 586,852 transcripts, only 10 of them are shown on the x-axis. mESCs_Mettl3_WT_fast5 outperforms the others in terms of query align lengths and mapping quality. The average number of reads with query length under 200 is 84,963 and over 200 is 9,391,757. This means 99.10% of the reads have a query length of greater than 200. The average number of reads of mapping quality less than 20 is 9,407,566 and over 20 is 115,700. This shows that only 1.21% of reads have a mapping quality higher than 20.

The above figure shows that there exist some variations among the read distributions of those six datasets in terms of query length and mapping quality. Most of the query lengths lie in between 0 to 1000 and the mapping quality over 40 is rare. Among all 586,852 transcripts, only some are present among the reads found in those six datasets.

Based on the results, we recommend using all datasets for m6A methylation using mouse reference sequence noting that the mapping quality is low.

#### 3.1.3. Curlcake ([24])

An artificially made RNA sequence is used to detect m6A methylation in Liu et al. [24]. There are four datasets which are explained below.

**Table 3.** This table explains the datasets for Curlcake ([24]) and m6A methylation and shows size (in GB), the number of reads and bases, corresponding references and base mapping, Tombo success rates and pass rates (see Section 2)

| Datasets | Size (GB) | #READ | #BASE | Reference | Base map% | Tombo % | Pass % |
| --- | --- | --- | --- | --- | --- | --- | --- |
| GSM3528749 | 17 | 66,736 | 57,953,462 | [24] | 94.66 | 77.96 | 76.65 |
| GSM3528750 | 41 | 134,374 | 114,498,490 | [24] | 64.13 | 40.93 | 45.93 |
| GSM3528751 | 229 | 846,595 | 903,318,154 | [24] | 95.17 | 77.57 | 79.73 |
| GSM3528752 | 164 | 638,860 | 535,900,320 | [24] | 74.78 | 44.28 | 53.58 |

Among these four datasets, GSM3528750 and GSM3528752 are methylated whereas GSM3528749 and GSM3528751 are not methylated. The base mappings, Tombo and pass success rates vary between methylated and non-methylated groups and the table shows that methylated groups perform worse than non-methylated groups.

**Figure 6.**
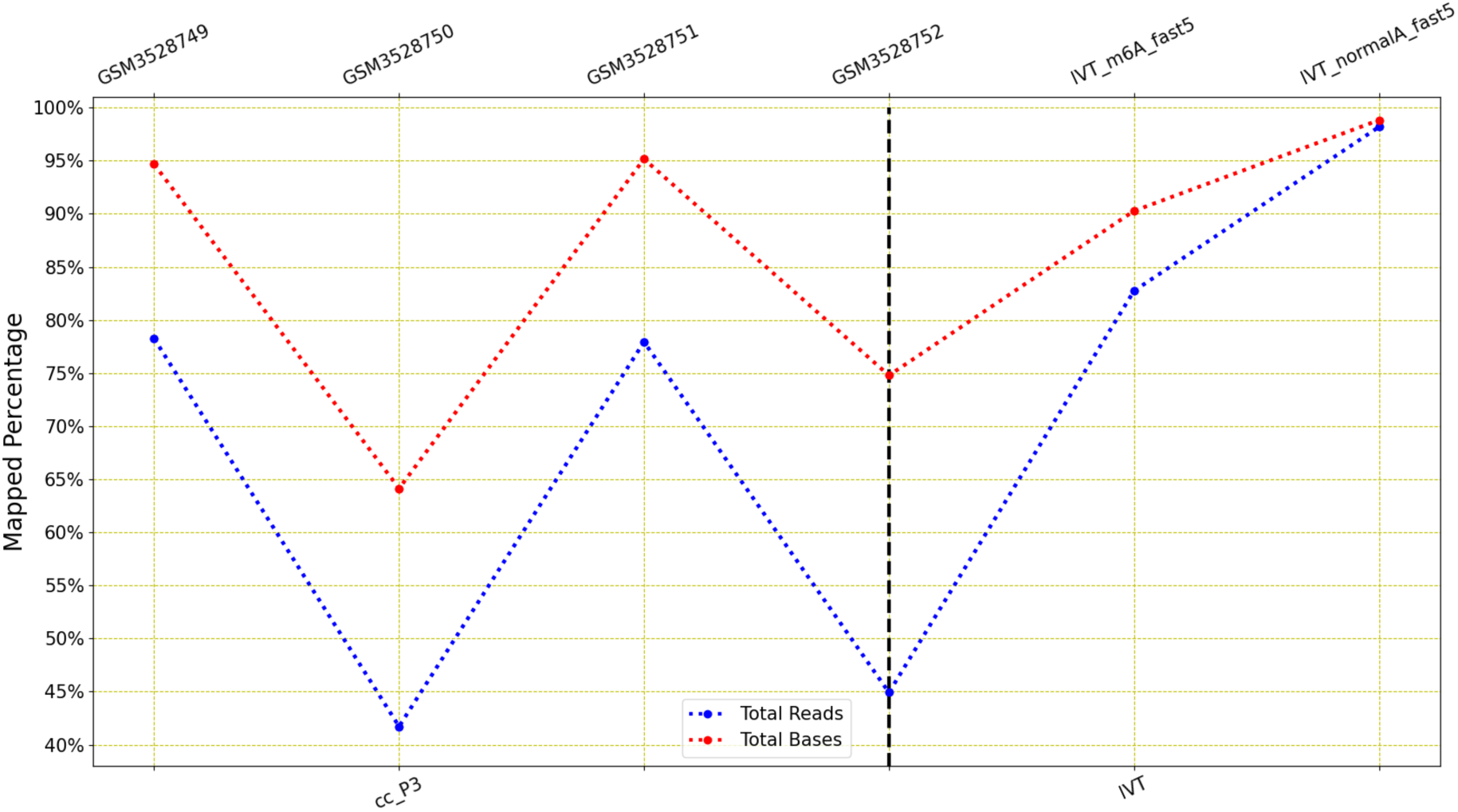
**LongReadSum mapping percentage of reads and bases of Curlcake (**[24]**) m6A and IVT m6A methylation data:** This figure shows that the mapping success rate varies significantly between methylated and non-methylated data.

The similar trend is also found among the mapping success rates for both bases and reads. The success rate is high (about 95%) for the non-methylated data and low (about 64% to 75%) for the methylated data.

**Figure 7.**
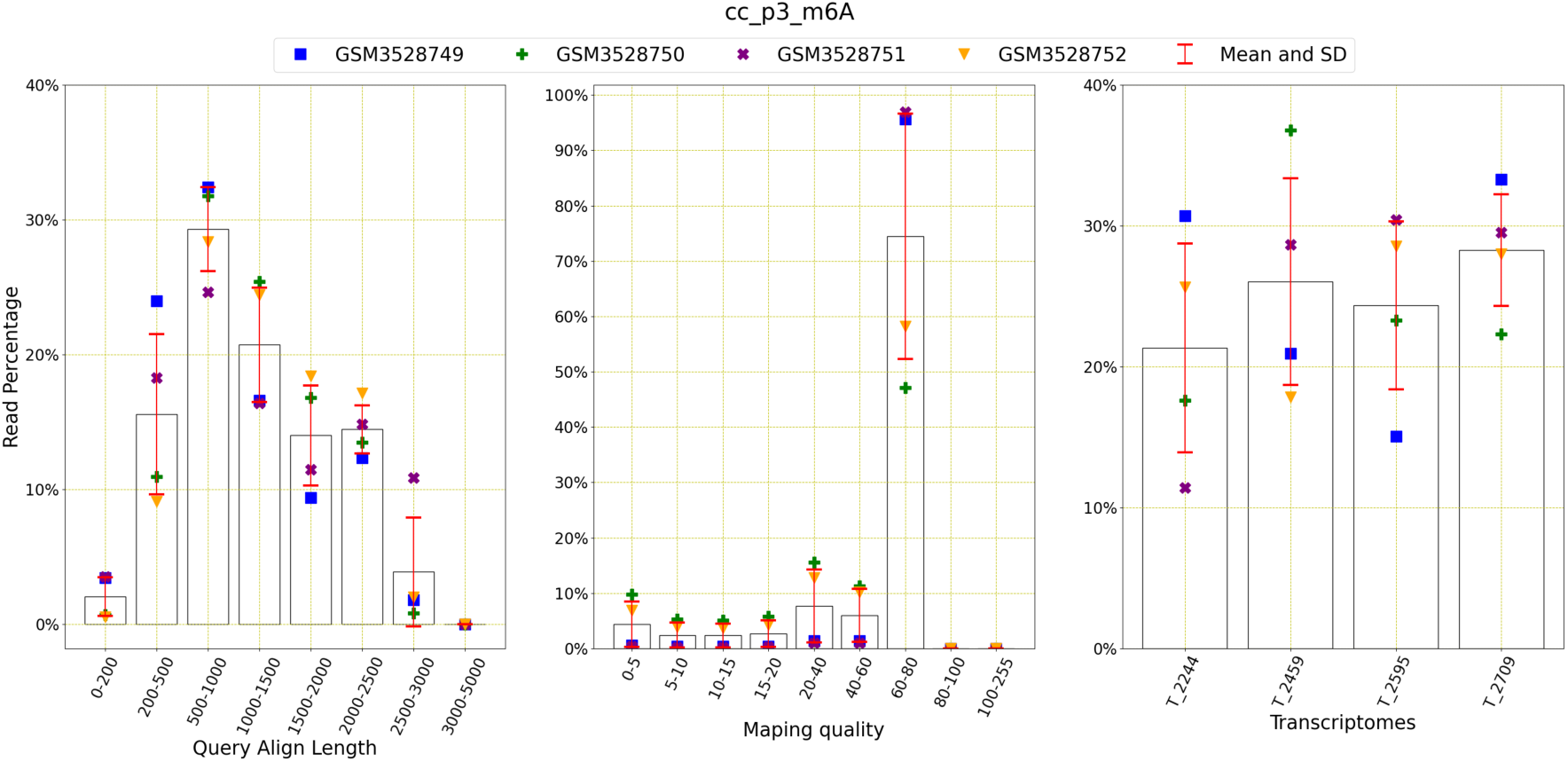
**Read distribution of Curlcake (**[24]**) m6A methylation data:** This figure shows the distribution of the reads for query align length, mapping quality and transcriptomes. The average number of reads with query length under 200 is 6,759 and over 200 is 256,304. This means 97.43% of the reads have a query length of greater than 200. The average number of reads of mapping quality less than 20 is 19,027 and over 20 is 244,036. This shows that 92.77% of reads have a mapping quality higher than 20.

The table indicates that the number of reads is not consistent among four datasets. The above figure indicates that most of the reads have a higher query align length of 500 to 2000. The mapping quality is also higher than 40 for most of the reads and all four transcriptomes appear almost equally among the reads.

#### 3.1.4. IVT

IVT is also a laboratory-made transcriptome which has been used for several types of methylation detection including m6A [28].

**Table 4.** This table explains the datasets for IVT and m6A methylation and shows size (in GB), the number of reads and bases, corresponding references and base mapping, Tombo success rates and pass rates (see Section 2)

| Datasets | Size | #READ | #BASE | Ref | Base map% | Tombo % | Pass % |
| --- | --- | --- | --- | --- | --- | --- | --- |
| IVT_m6A_fast5 | 90 | 1,482,437 | 1,053,858,300 | [28] | 90.27 | 59.44 | 88.65 |
| IVT_normalA_fast5 | 19 | 383,209 | 238,341,386 | [28] | 98.82 | 96.63 | 96.95 |

Here, IVT_m6A_fast5 is the methylated and IVT_normalA_fast5 is not methylated. The datasets vary in sizes and numer of reads and Tombo and pass rate. Tombo and and pass rates are lower for the methylated data. However, the base mapping success rate is high, over 90% for both of these datasets.

**Figure 8.**
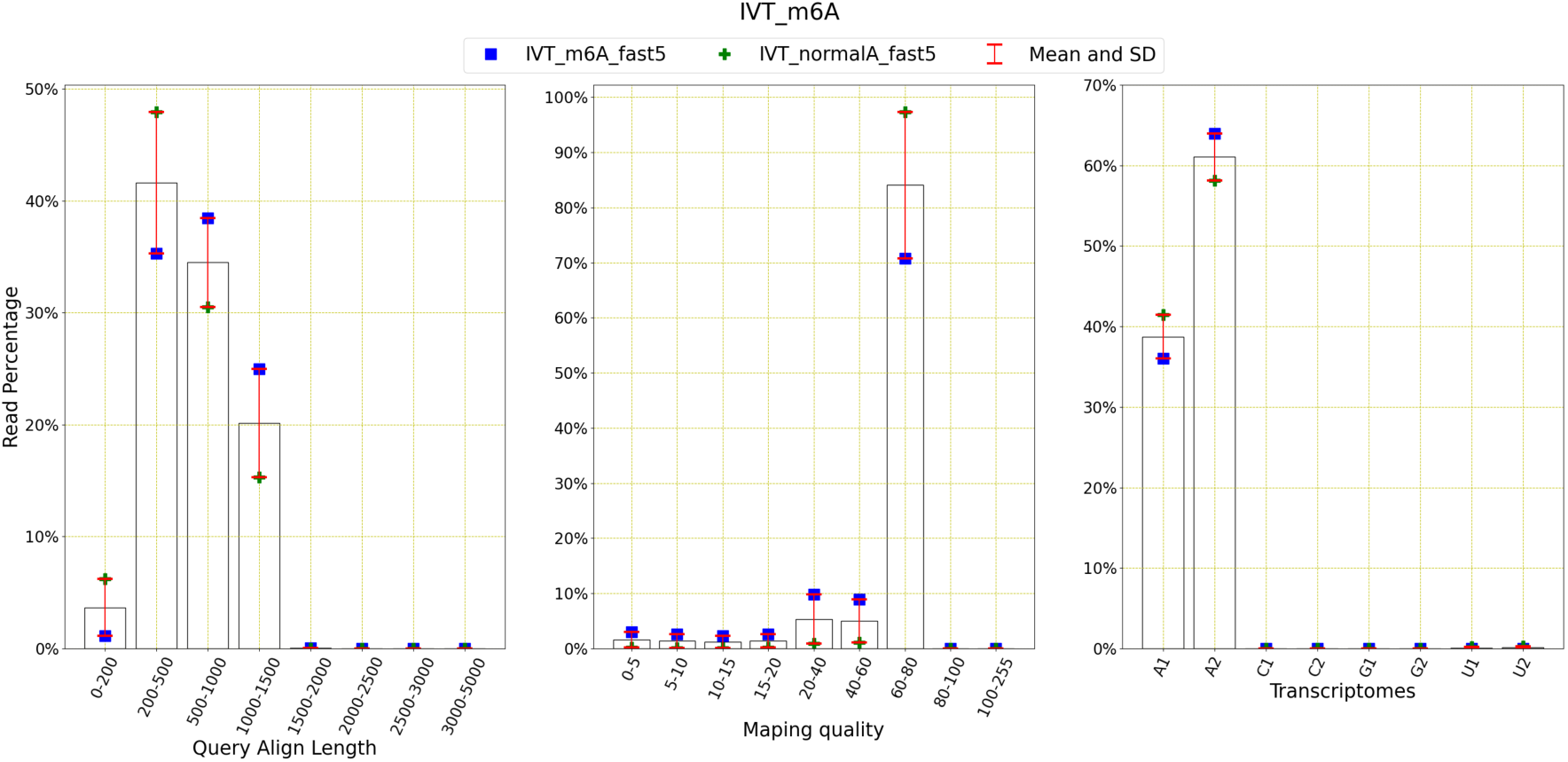
Read distribution of IVT m6A methylation data: This figure shows the distribution of the reads for query align length, mapping quality and transcriptomes. The average number of reads with query length under 200 is 18,961 and over 200 is 793,816. This means 97.67% of the reads have a query length of greater than 200. The average number of reads of mapping quality less than 20 is 66,927 and over 20 is 745,850. This shows that 91.77% of reads have a mapping quality higher than 20.

The average number of reads with query length under 200 is 89,813 and over 200 is 6,263,805. This means 98.59% of the reads have a query length of greater than 200. The average number of reads of mapping quality less than 20 is 6,318,666 and over 20 is 122,849. This shows that only 1.91% of reads have a mapping quality higher than 20.

IVT also contains six transcriptomes, and each pair is responsible for one methylation type. The transcriptomes A1 and A2 are developed to detect A nucleotide type of methylations such as m6A and the above figure shows that A2 appears more in reads than A1.

Based on these results, all the datasets provided for both IVT and curlcake [24] are recommended to be used due to their high mapping quality and good success rates for Tombo and base mapping. This is also to be noted that methylated datasets have lower rates than non-methylated datasets.

#### 3.1.5. Yeast

The ONT datasets for m6A methylation using yeast come from Liu et al. [24]. There are six datasets present all together.

**Table 5.**
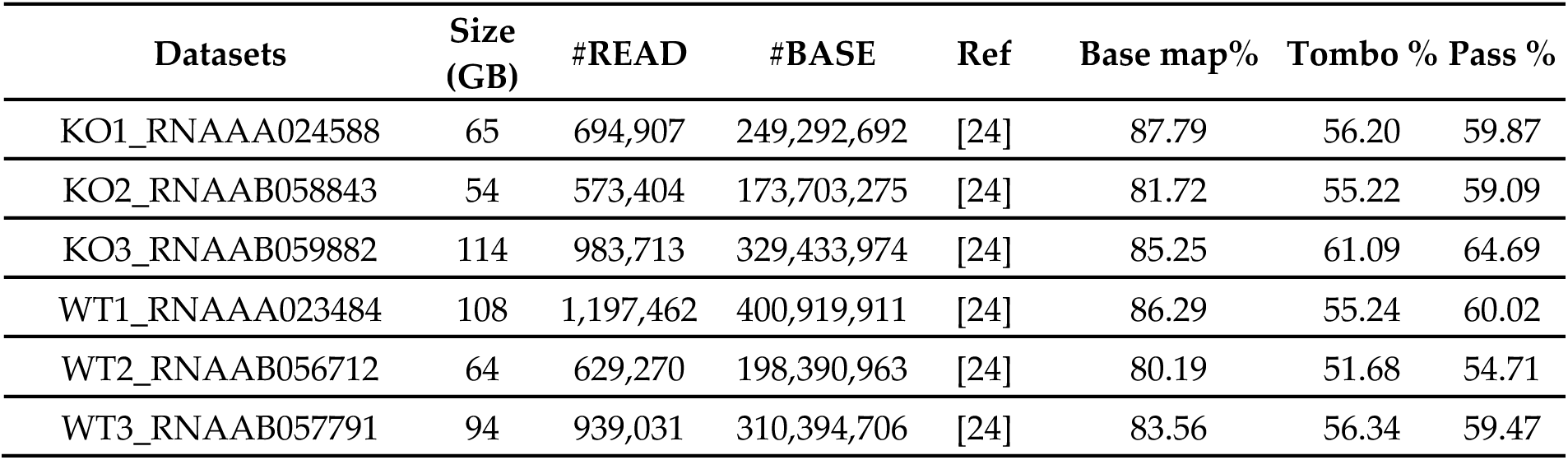
This table explains the datasets for yeast and m6A methylation and shows size (in GB), the number of reads and bases, corresponding references and base mapping, Tombo success rates and pass rates (see Section 2)

| Datasets | Size (GB) | #READ | #BASE | Ref | Base map% | Tombo % | Pass % |
| --- | --- | --- | --- | --- | --- | --- | --- |
| KO1_RNAAA024588 | 65 | 694,907 | 249,292,692 | [24] | 87.79 | 56.20 | 59.87 |
| KO2_RNAAB058843 | 54 | 573,404 | 173,703,275 | [24] | 81.72 | 55.22 | 59.09 |
| KO3_RNAAB059882 | 114 | 983,713 | 329,433,974 | [24] | 85.25 | 61.09 | 64.69 |
| WT1_RNAAA023484 | 108 | 1,197,462 | 400,919,911 | [24] | 86.29 | 55.24 | 60.02 |
| WT2_RNAAB056712 | 64 | 629,270 | 198,390,963 | [24] | 80.19 | 51.68 | 54.71 |
| WT3_RNAAB057791 | 94 | 939,031 | 310,394,706 | [24] | 83.56 | 56.34 | 59.47 |

Among six datasets, there are three wild types and three knocked out types. The size of those datasets along with the number of reads and bases do not vary between these two groups. The base mapping success rates are higher than Tombo and pass rates. The base mapping success rates are over 80%, where the others are always lower than 65%.

There is not much variation present among all six datasets types. The mapped read percentage is lower than the base percentages.

**Figure 9.**
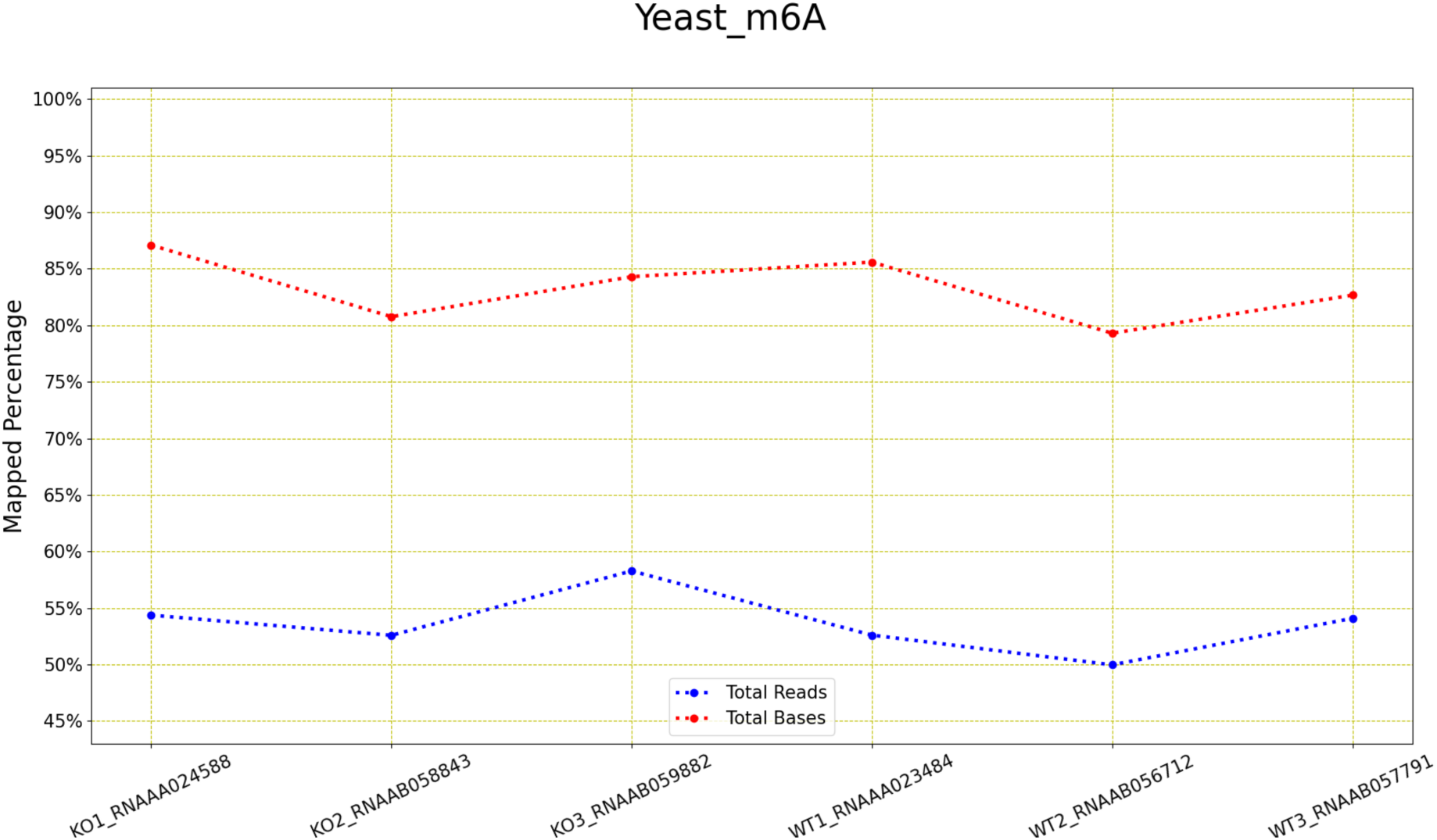
LongReadSum mapping percentage of reads and bases of yeast m6A methylation data: This figure shows that the mapping success rates are the same between all six datasets.

**Figure 10.**
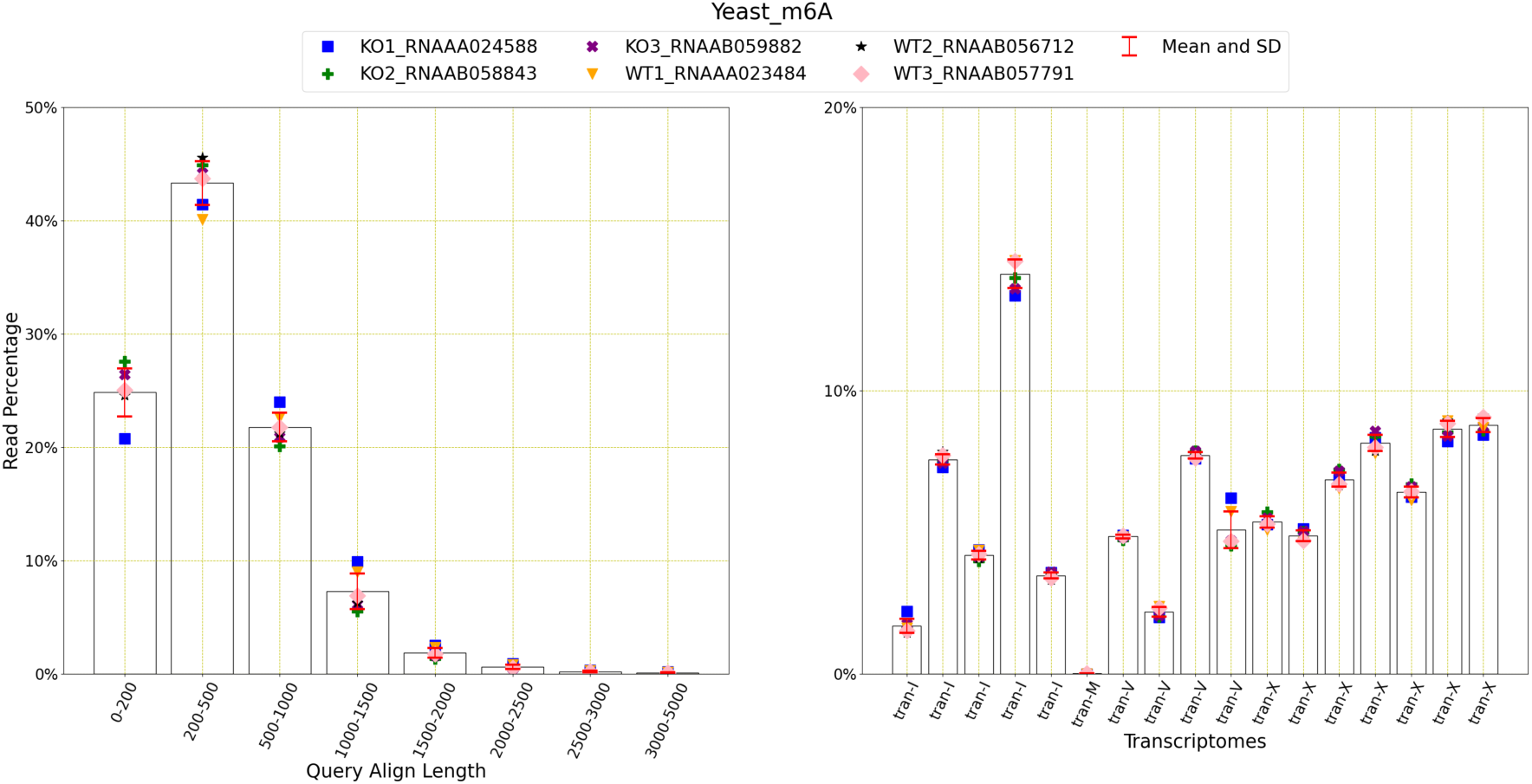
Read distribution of Yeast m6A methylation data: This figure shows the distribution of the reads for query align length, mapping quality and transcriptomes. The average number of reads with query length under 200 is 115,231 and over 200 is 347,176. This means 75.08% of the reads have a query length of greater than 200. The average number of reads of mapping quality less than 20 is 25,166 and over 20 is 437,771. This shows that 94.56% of reads have a mapping quality higher than 20.

The above figure confirms that similar trend is present between all six datasets and WT and KO types. Most of the reads contain query length of 200-500 and mapping qualities are mostly higher than 60. However, the read distribution among the transcriptome varies significantly.

Based on the results, we show that all datasets can be used here, noting that the difference between read and base success rates.

#### 3.1.6. E Coli

There is only one dataset present for m6A methylation using E Coli.

For this dataset, the base mapping success, Tombo and pass rates are good, ranging between 83 to 90%.

**Table 6.** This table explains the datasets for E Coli and m6A methylation and shows size (in GB), the number of reads and bases, corresponding references and base mapping, Tombo success rates and pass rates (see Section 2)

| Datasets | Size (GB) | #READ | #BASE | Ref | Base map% | Tombo % | Pass % |
| --- | --- | --- | --- | --- | --- | --- | --- |
| EC_rRNA_dRNA_fast5 | 35 | 324,837 | 301,183,340 | [28] | 90.02 | 83.79 | 84.28 |

**Figure 11.**
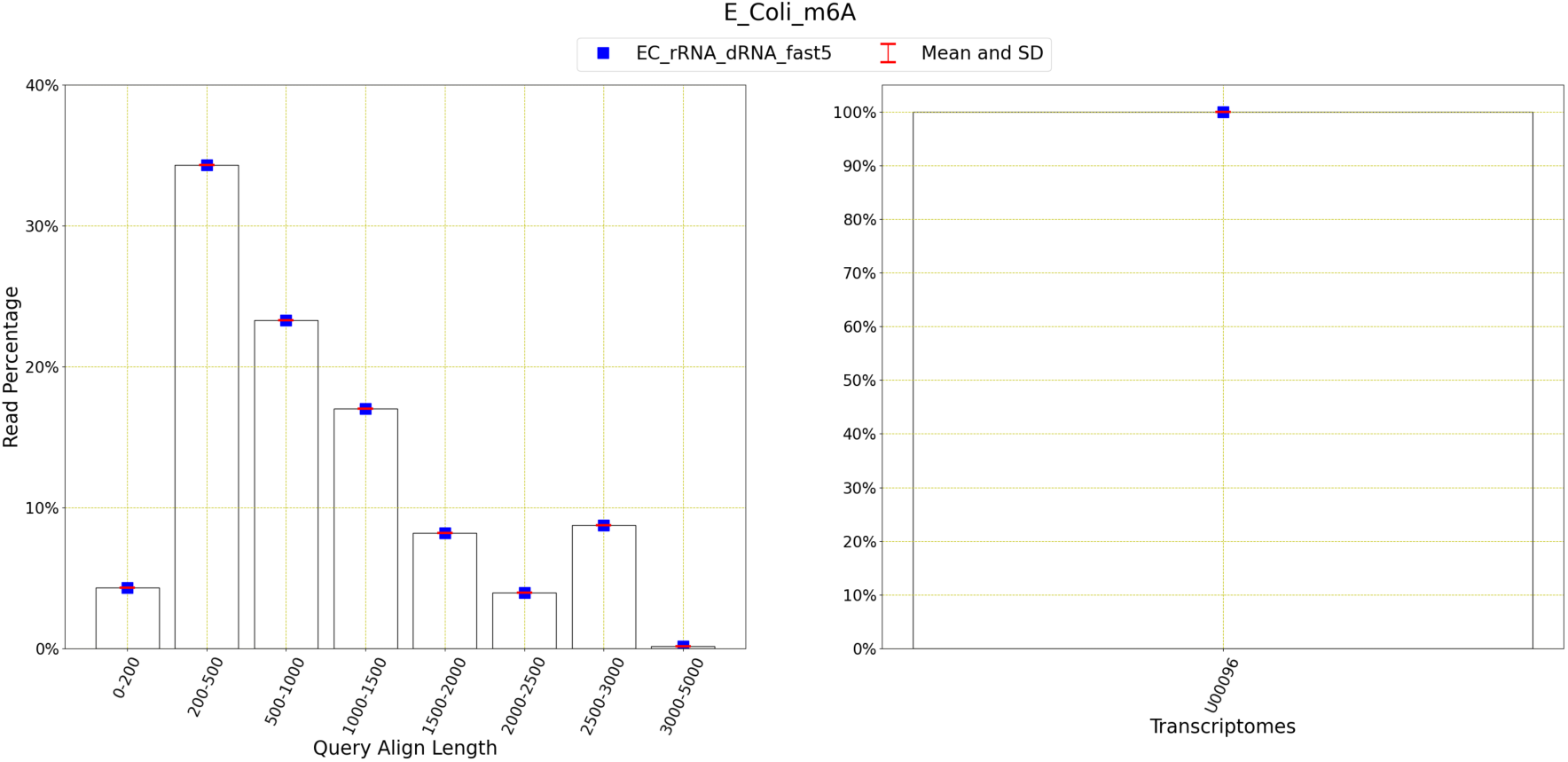
Read distribution of E Coli m6A methylation data: This figure shows the distribution of the reads for query align length, mapping quality and transcriptomes. The average number of reads with query length under 200 is 61,667 and over 200 is 1,362,206. This means 95.67% of the reads have a query length of greater than 200. The average number of reads of mapping quality less than 20 is 1,374,965 and over 20 is 51,086. This shows that 3.58% of reads have a mapping quality higher than 20.

The figure shows that most reads have the query length of 200 to 500 and the mapping quality of 0 to 20. The E coli reference sequence (shall I add a reference here?) contains only one transcript, thus all the reads are mapped using that transcript only.

#### 3.1.7. Arabidopsis

The ONT datasets for m6A methylation using Arabidopsis originate from Zhong et al. [10]. Among those 15 datasets, there are five ‘col0_drs’, two ‘col0_drs_adapter’, four ‘vir1’ and four ‘VIRc’ types.

**Table 7.**
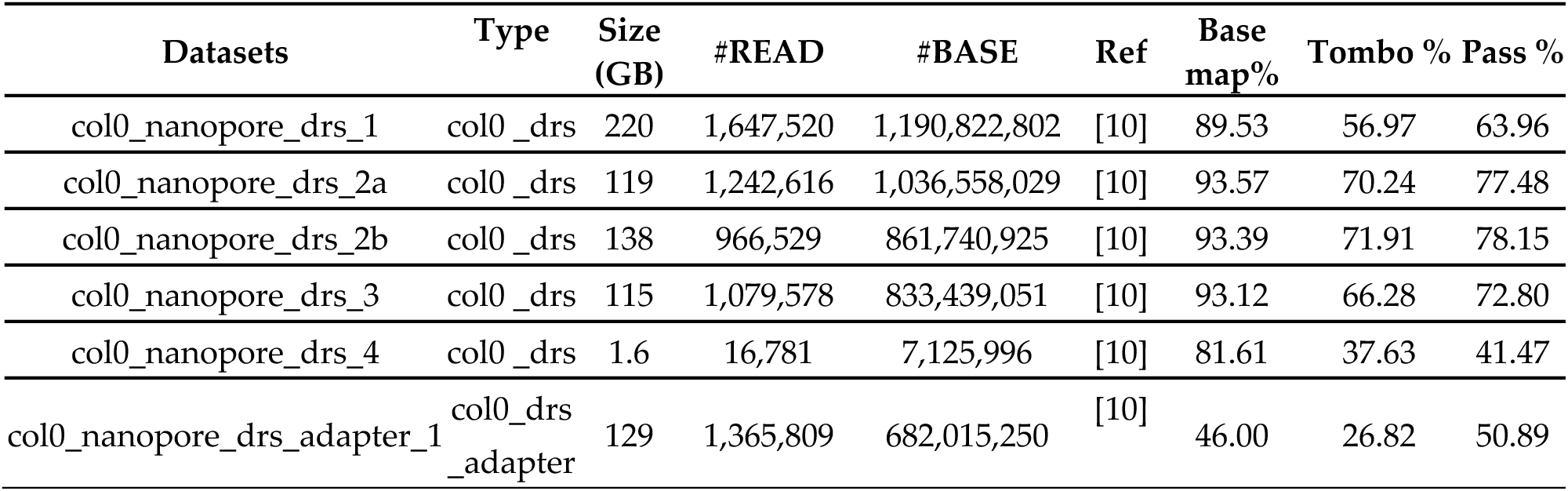
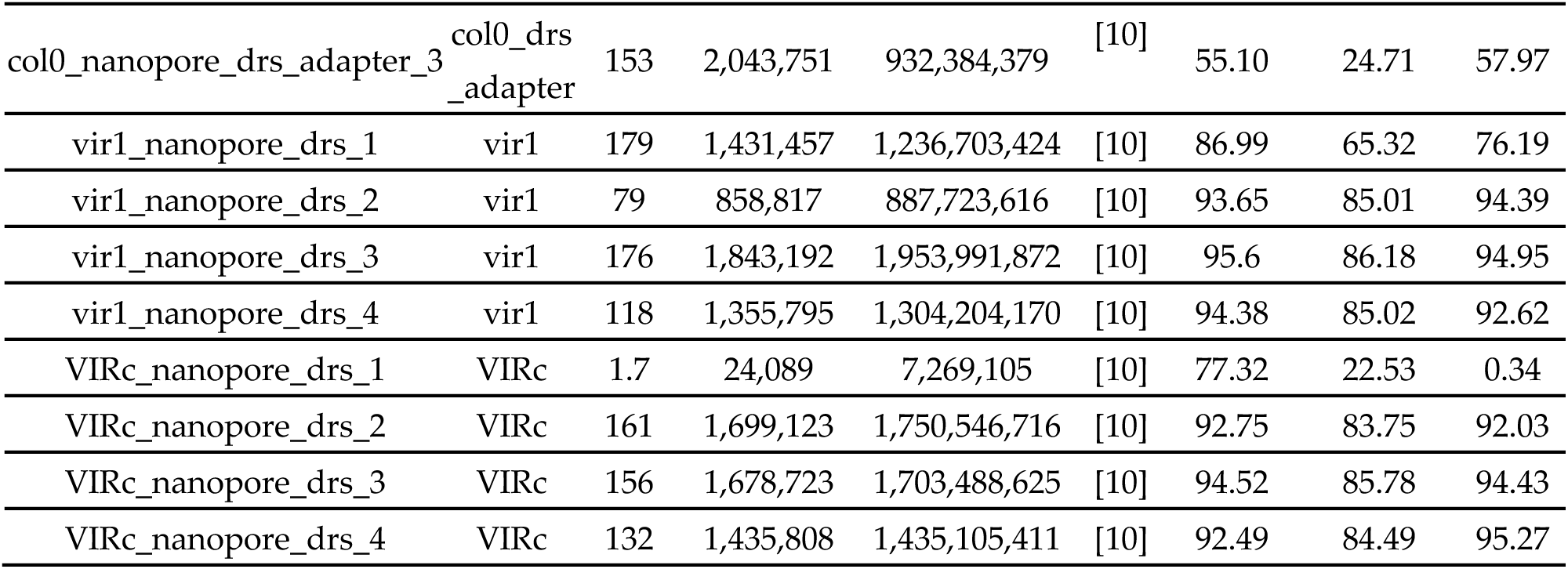
This table explains the datasets for arabidopsis and m6A methylation and shows size (in GB), the number of reads and bases, corresponding references and base mapping, Tombo success rates and pass rates (see Section 2)

The size, number of reads and bases for all 15 datasets are not consistent. Data types such as ‘col0_drs_adapter’ didn’t perform as good as others. Also, ‘VIRc_nanopore_drs_1’ is smaller than others in the ‘VIRc’ category and performed worse.

**Figure 12.**
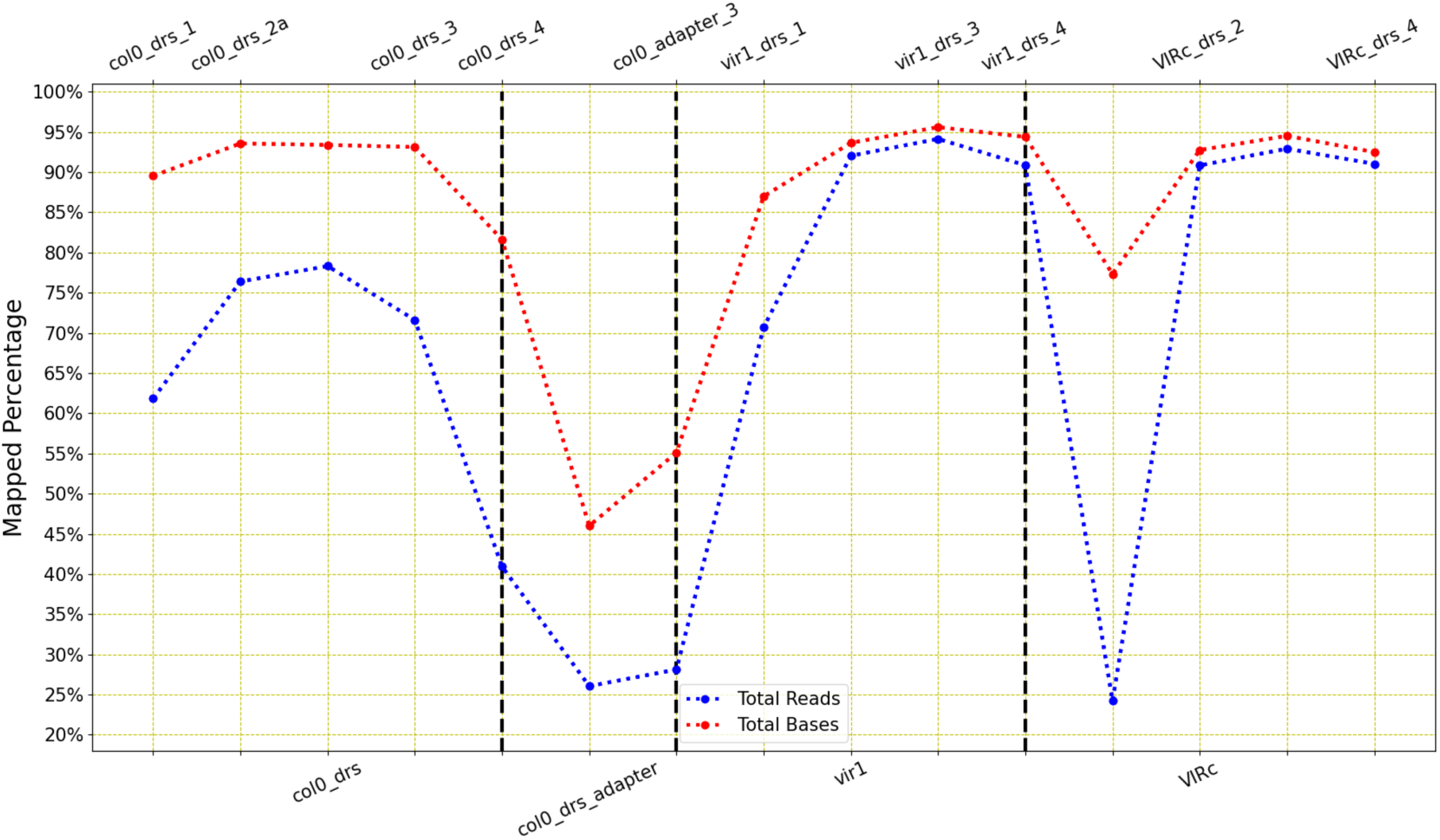
LongReadSum mapping percentage of reads and bases of arabidopsis m6A methylation data: This figure shows that the mapping success rates vary a lot between all 15 datasets.

**Figure 13.**
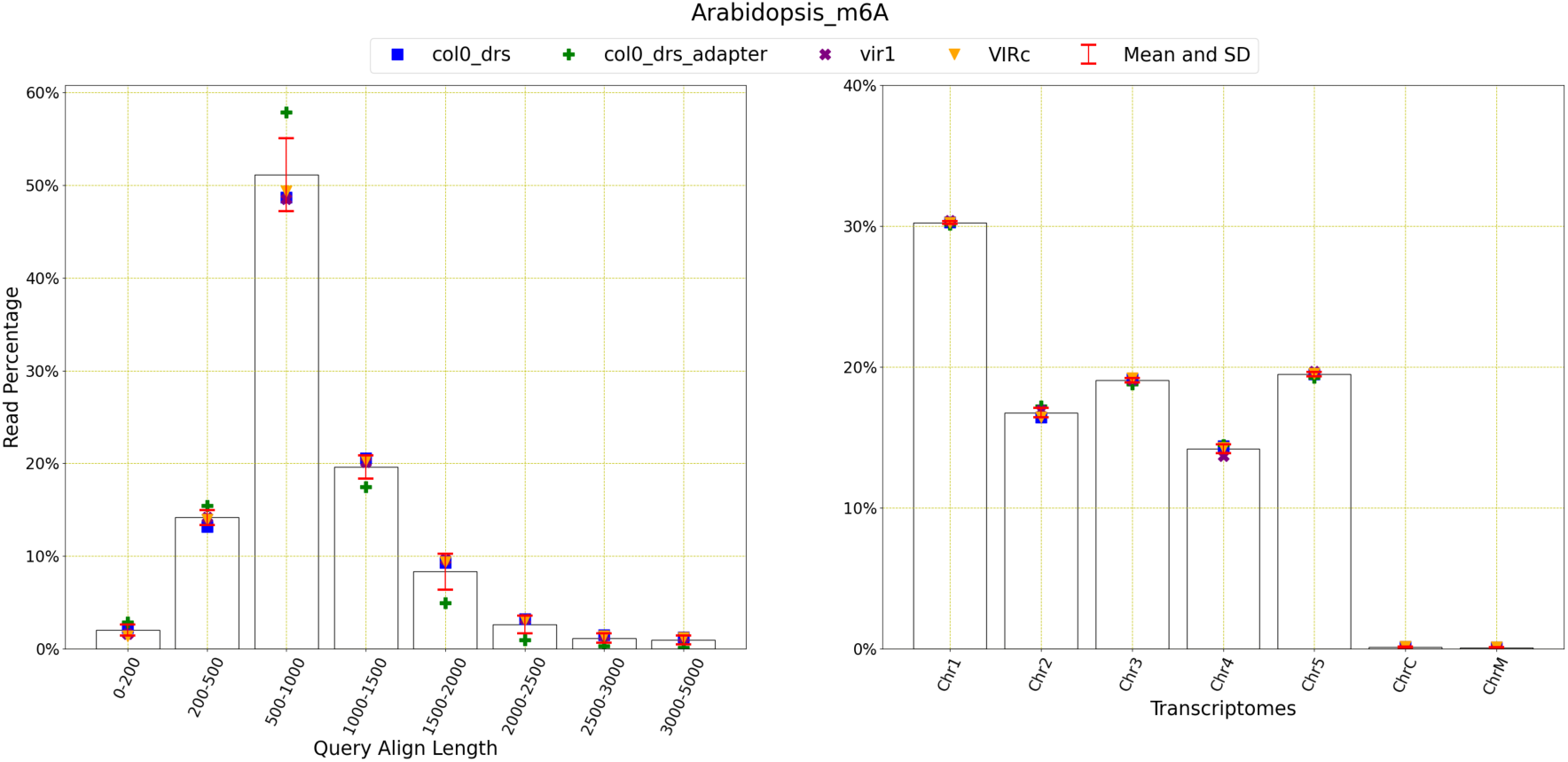
Read distribution of Arabidopsis m6A methylation data: This figure shows the distribution of the reads for query align length, mapping quality and transcriptomes. The average number of reads with query length under 200 is 16,238 and over 200 is 872,638. This means 98.17% of the reads have a query length of greater than 200. The average number of reads of mapping quality less than 20 is 22,276 and over 20 is 867,269. This shows that 97.5% of reads have a mapping quality higher than 20.

The above figure provides further evidence to the fact that ‘col0_drs_adapter’ group performed the worst, even though the number of reads and bases are the similar to the other datasets. It can also be noticed that the difference between the mapping percentages of reads and bases is bigger for ‘col0_drs’ than ‘vir1’.

**Figure 14.**
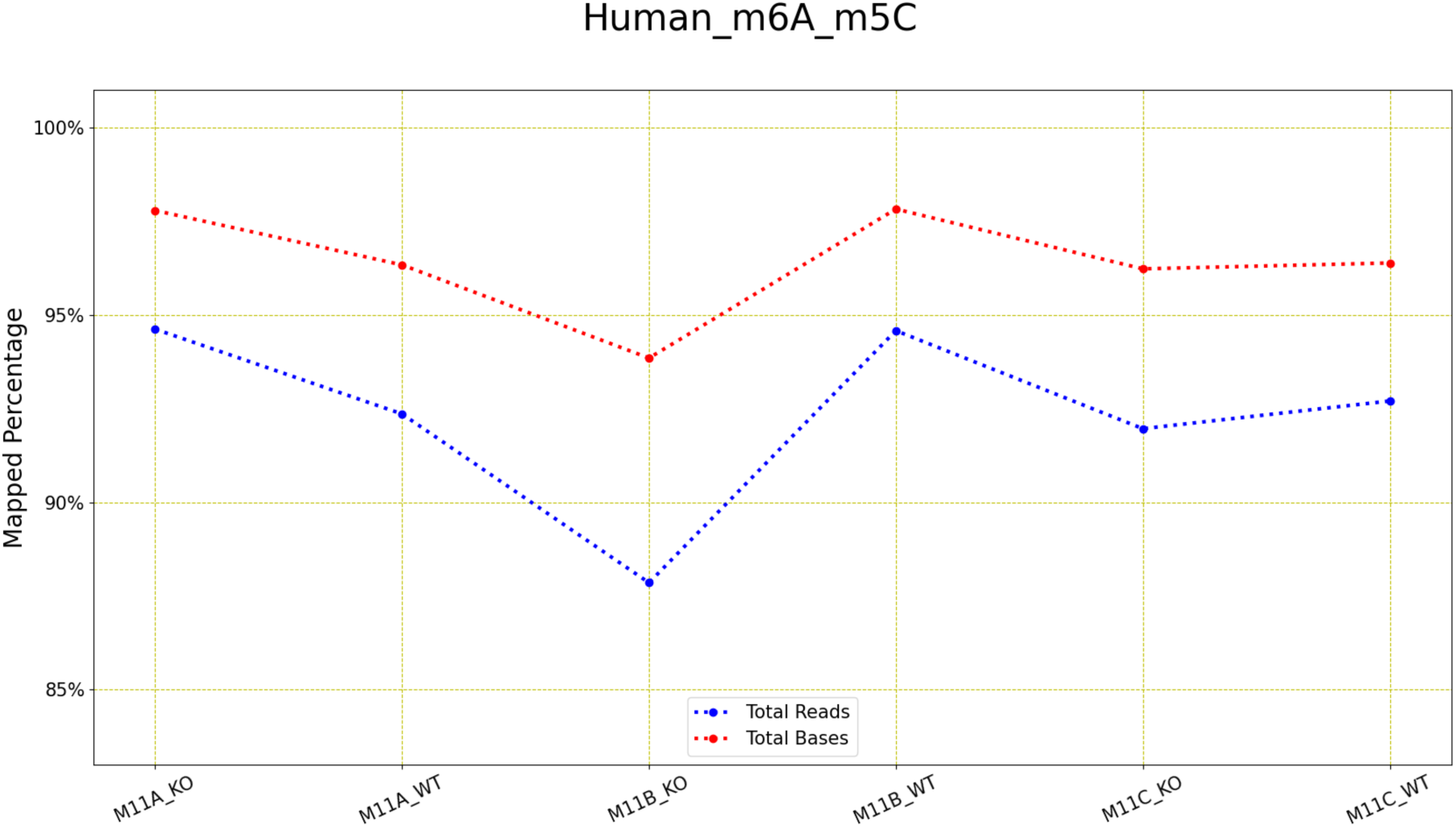
LongReadSum mapping percentage of reads and bases of human m6A and m5C methylation data: This figure shows that the mapping success rates are good with slight variation in performance.

**Figure 15.**
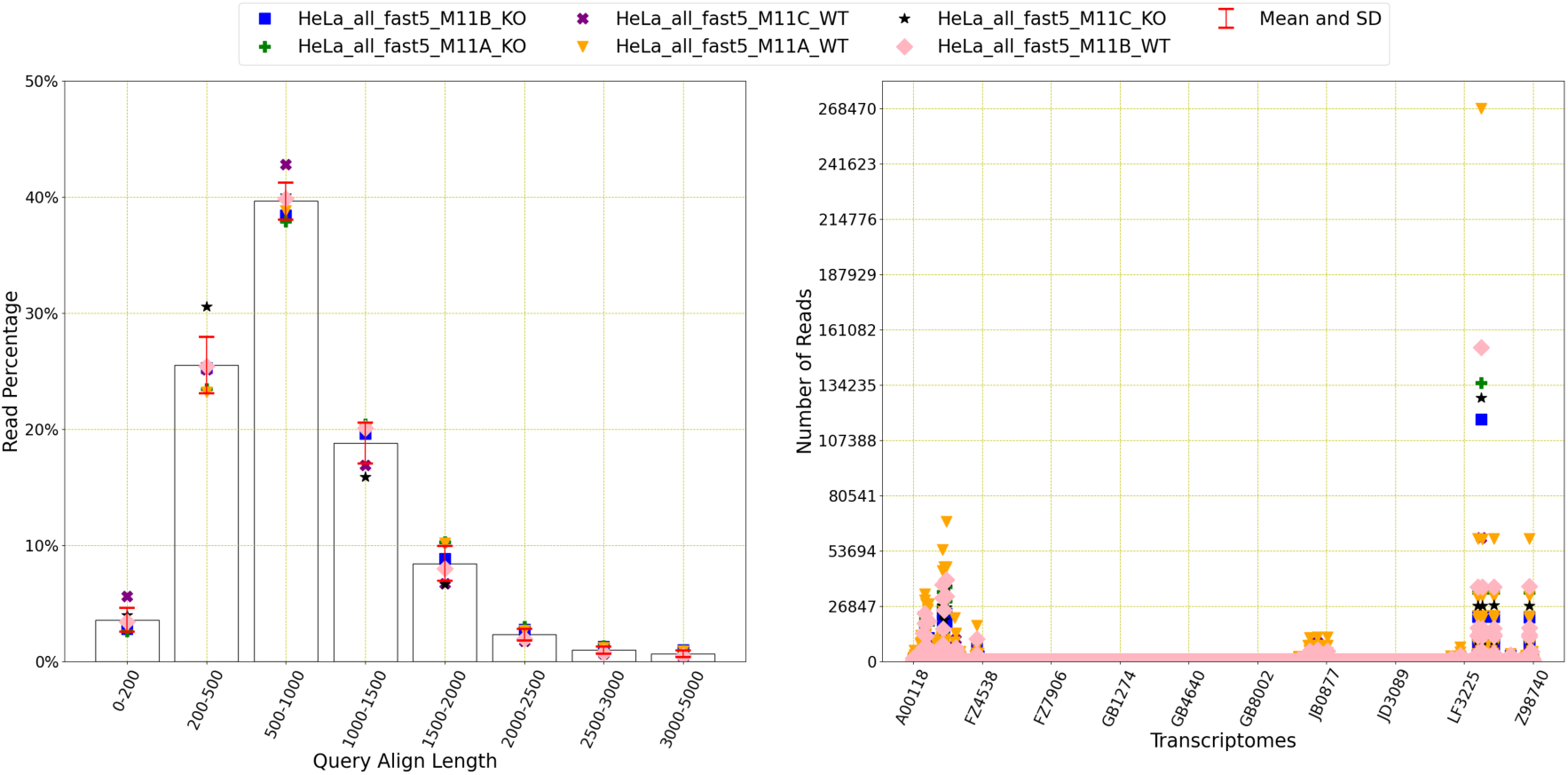
Read distribution of Human m6A and m5C methylation data: This figure shows the distribution of the reads for query align length, mapping quality and transcriptomes. The average number of reads with query length under 200 is 118,988 and over 200 is 3,397,568. This means 96.62% of the reads have a query length of greater than 200. The average number of reads of mapping quality less than 20 is 3,428,889 and over 20 is 114,789. This shows that 3.24% of reads have a mapping quality higher than 20.

The above figure shows that there is a similar trend present for all four types of datasets. The query length lies between 500 to 1000 and the mapping quality is higher than 60 for most of the reads. Most of the reads contain Chr1 transcriptome and ChrM is almost missing for all four datasets types.

Based on the results, we suggest that ‘vir1’ dataset type is the best among the others as it has higher read and base success rates than the other datasets. It is also to be noted that the mapping qualities are high (>60) for all datasets.

### 3.2. m6A and m5C

There are also datasets which contain ONT data specifically designed for both m6A and m5C methylations. They are summarized here according to the reference sequence used to prepare the data.

#### 3.2.1. Human

The ONT datasets prepared for both m6A and m5C methylations using human reference sequence are presented in Mateos et al.. There are six of them and they can be categorized as wild type (WT) and knocked out (KO).

**Table 8.** This table explains the datasets for human and m6A and m5C methylation and shows size (in GB), the number of reads and bases, corresponding references and base mapping, Tombo success rates and pass rates (see Section 2)

| Datasets | Size (GB) | #READ | #BASE | Ref | Base map% | Tombo % | Pass % |
| --- | --- | --- | --- | --- | --- | --- | --- |
| HeLa_all_fast5_M11A_KO | 44 | 630,084 | 595,179,197 | [27] | 97.02 | 92.44 | 94.59 |
| HeLa_all_fast5_M11A_WT | 50 | 706,673 | 682,067,109 | [27] | 95.48 | 89.94 | 91.61 |
| HeLa_all_fast5_M11B_KO | 22 | 332,790 | 265,554,918 | [27] | 92.61 | 85.74 | 89.97 |
| HeLa_all_fast5_M11B_WT | 101 | 1,379,595 | 1,334,415,581 | [27] | 96.73 | 92.32 | 94.59 |
| HeLa_all_fast5_M11C_KO | 53 | 806,061 | 651,947,214 | [27] | 95.11 | 89.36 | 91.70 |
| HeLa_all_fast5_M11C_WT | 48 | 720,023 | 634,760,138 | [27] | 95.61 | 90.60 | 93.08 |

\* Tables may have a footer.

The above six datasets show high values (over 85%) for base mapping, Tombo and pass success rates, even though their size and number of reads/bases vary.

The above table and figure show that all six ONT datasets containing both WT and KO have good performance for mapping percentage for both the reads and bases and also performed well for Tombo.

The above figure shows that even thought the number of reads vary, most of the reads have the query length of 500 to 1000 and the mapping quality is less than 20. The reads do not contain all the human transcriptomes and only a few are present.

**Table 9.** This table explains the datasets for mouse and m6A and m5C methylation and shows size (in GB), the number of reads and bases, corresponding references and base mapping, Tombo success rates and pass rates (see Section 2)

| Datasets | Type | Size (GB) | #READ | #BASE | Ref | Base map% | Tombo % | Pass % |
| --- | --- | --- | --- | --- | --- | --- | --- | --- |
| AR1_Mousebrain | 386ng | 1.1 | 10,459 | 7,589,420 | [27] | 71.76 | 64.43 | 66.34 |
| AR2_Mousebrain | 812ng | 5.6 | 66,580 | 51,241,709 | [27] | 91.02 | 82.42 | 84.32 |
| AR8_Mousebrain | 4720ng | 5.1 | 155,197 | 26,040,692 | [27] | 60.85 | 29.41 | 35.82 |
| AR9_Mousebrain | 4720ng | 6.1 | 151,785 | 36,914,996 | [27] | 70.21 | 43.91 | 49.07 |
| AR12_Mousebrain | 2573ng | 12 | 171,577 | 160,433,892 | [27] | 91.44 | 85.29 | 87.40 |
| AR13_Mousebrain | 2573ng | 29 | 404,000 | 404,479,731 | [27] | 96.63 | 92.09 | 93.83 |
| AR14_Mousebrain | 2573ng | 33 | 449,778 | 433,114,621 | [27] | 95.50 | 89.42 | 91.38 |
| AR15_Mousebrain | 2573ng | 40 | 544,206 | 267,372,968 | [27] | 95.06 | 56.15 | 91.11 |
| AR16_Mousebrain | 1546ng | 14 | 196,000 | 190,029,100 | [27] | 97.22 | 92.74 | 94.73 |
| AR17_MouseBrain | 1546ng | 23 | 312,590 | 308,720,934 | [27] | 96.45 | 90.68 | 92.17 |
| AR18_Mousebrain | 1546ng | 35 | 465,467 | 472,517,705 | [27] | 96.44 | 91.27 | 92.66 |
| N7_Mousebrain | 500ng | 91 | 933,011 | 964,127,856 | [27] | 96.48 | 90.82 | 92.16 |
| N8_Mousebrain | 500ng | 30 | 332,469 | 313,129,171 | [27] | 96.77 | 87.45 | 89.07 |
| N17_Mousebrain | 500ng | 4.3 | 42,630 | 39,075,326 | [27] | 93.67 | 84.94 | 87.17 |

#### 3.2.2. Mouse

The ONT datasets for m6A and m5C methylations using the mouse sequence is also presented at Mateos et al. [27]. There are altogether 14 datasets and they are separated into six groups.

The base mapping, Tombo and pass rates are good for some datasets and poor for a few such as 386ng and 4720ng. The size of the datasets also vary a lot.

**Figure 16.**
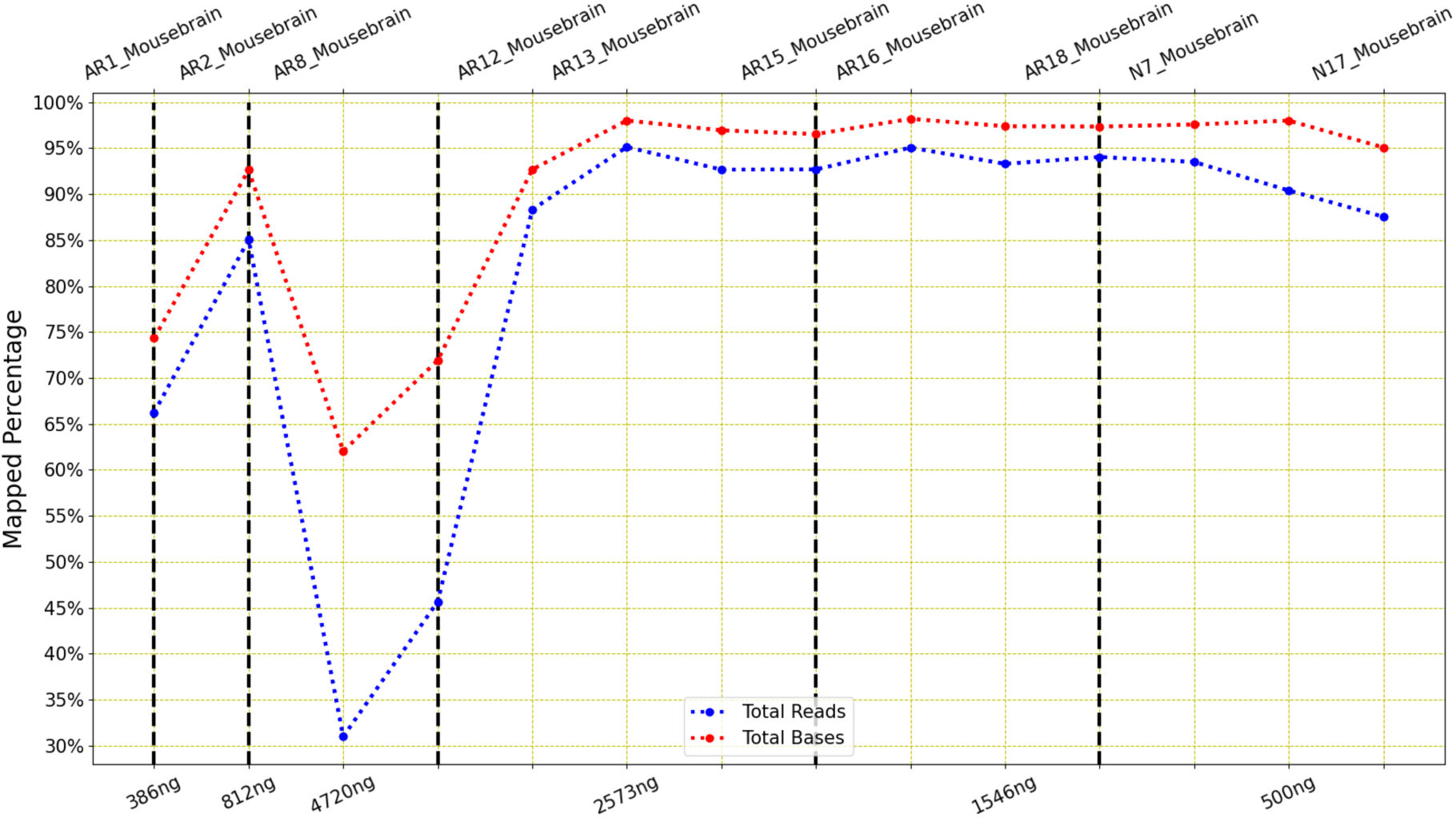
LongReadSum mapping percentage of reads and bases of mouse m6A and m5C methylation data: This figure shows a huge deviation of mapping success rates for one dataset to the other. The mapping ratio gap between the reads and bases are also not consistent at all.

There is a huge performance variation present in the figure above. The mapping success ratio varies from as high as 95% to as low as 30%. However, most of the datasets performed well (over 85%) in general, except 386ng and 4720ng. It can also be noticed that the difference between ratios of reads and bases is also large for 4720ng datasets.

**Figure 17.**
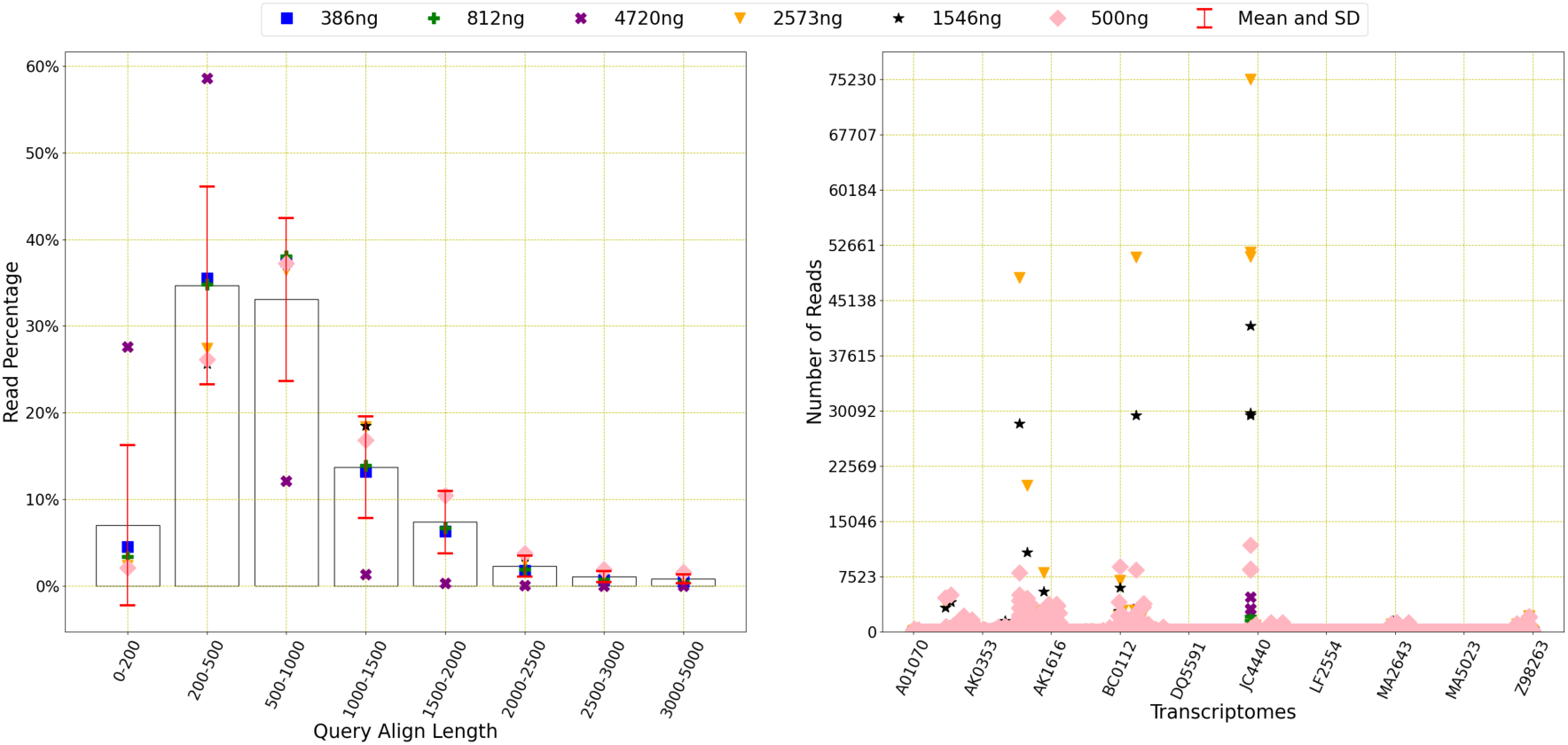
Read distribution of Mouse m6A and m5C methylation data: This figure shows the distribution of the reads for query align length, mapping quality and transcriptomes. The average number of reads with query length under 200 is 32379 and over 200 is 872710. This means 96.42% of the reads have a query length of greater than 200. The average number of reads of mapping quality less than 20 is 894066 and over 20 is 22662. This shows that 2.47% of reads have a mapping quality higher than 20.

Reads are not distributed in a similar pattern for the query align lengths. For 500ng, most reads contain the query length of 500 to 1000. However, 4720ng contains reads who mostly contain the query length of 200-500. However, the mapping quality is lower than 20 for most of reads for all six types of datasets. It can also be noticed that not all transcripts appear in reads for all six datasets.

### 3.3. Pseudouridine

The methylation type pseudouridine (Ψ) is also been investigated by some authors even though they are not that common.

#### 3.3.1. Curlcake ([9])

Another curlcake reference sequence was developed by Begik et al. [9]. This was artificially made. For pseudouridine methylation type, there are only three datasets as explained below.

**Table 10.** This table explains the datasets for curlcake ([9]) and pseudouridine (Ψ) methylation and shows size (in GB), the number of reads and bases, corresponding references and base mapping, Tombo success rates and pass rates (see Section 2)

| Datasets | Size (GB) | #READ | #BASE | Methyl types | Ref | Base map% | Tombo % | Pass % |
| --- | --- | --- | --- | --- | --- | --- | --- | --- |
| pU | 3.5 | 42,386 | 20,208,901 | Ψ | [9] | 0.31 | 0.39 | 15.35 |
| RNAAB063141_00DMS | 3.6 | 77,629 | 16,071,894 | UNM-S | [9] | 0.01 | 0.01 | 55.03 |
| RNA198724_stoichiometry | 62 | 457,955 | 679,378,093 | Ψ with varying stoichiometries | [9] | 97.82 | 90.01 | 88.36 |

\* Tables may have a footer.

The pU and RNAAB063141_00DMS datasets are not big in size and their base mapping, Tombo and pass rate is extremely worse compared to the other datasets presented in this paper. The RNA198724_stoichometry datasets performed well though having values higher than 88%.

**Figure 18.**
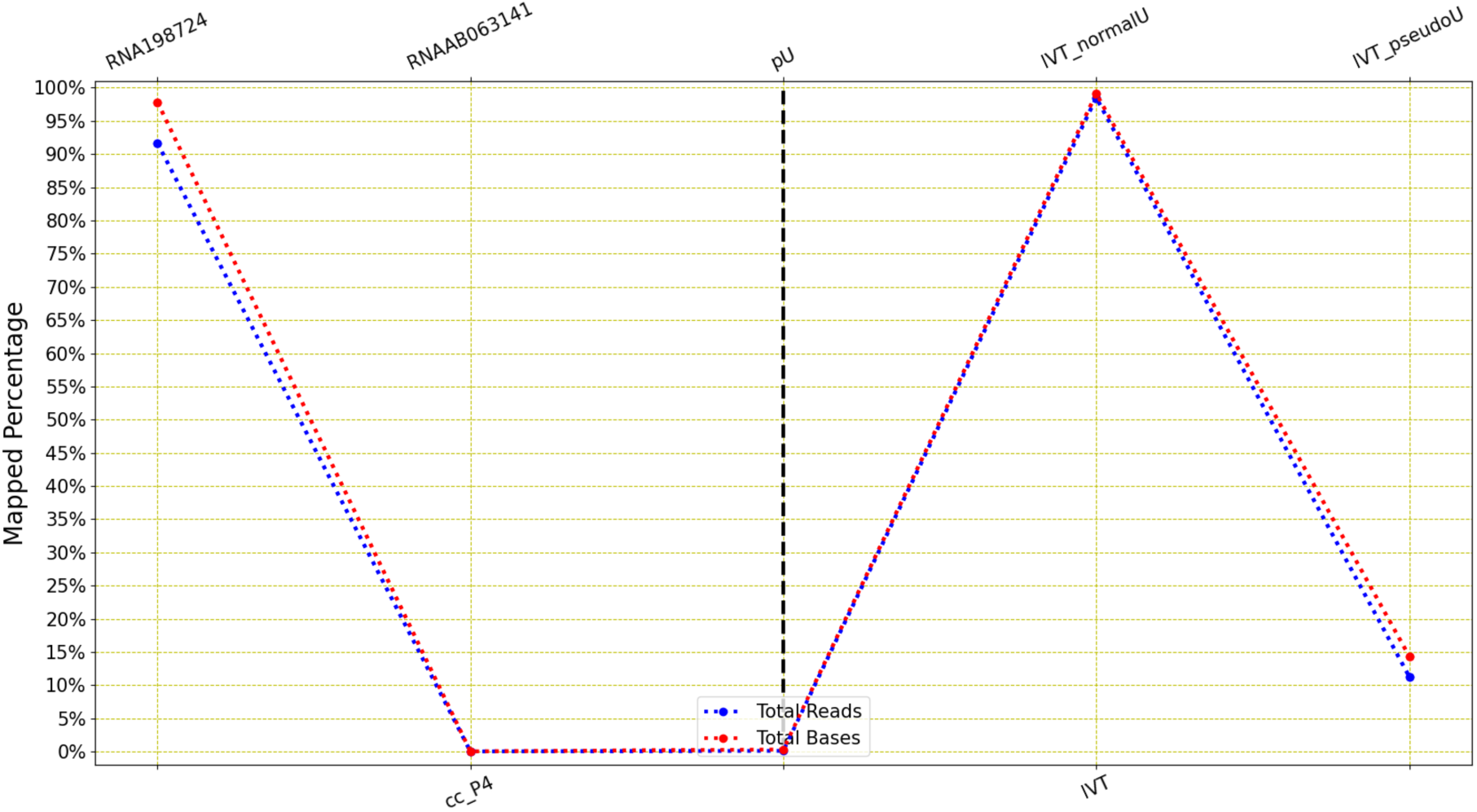
LongReadSum mapping percentage of reads and bases of curlcake and IVT pseudouridine methylation data: This figure shows a huge deviation of mapping success rates for one dataset to the other for both curcake and IVT. Here, one dataset performed really and well and other two have mapping ratio very close to zero for curlcake. Here, one dataset performed really and well and other didn’t.

The above figure shows that ONT datasets for pseudouridine have contrasting performance, where one dataset perform well and other two don’t.

**Figure 19.**
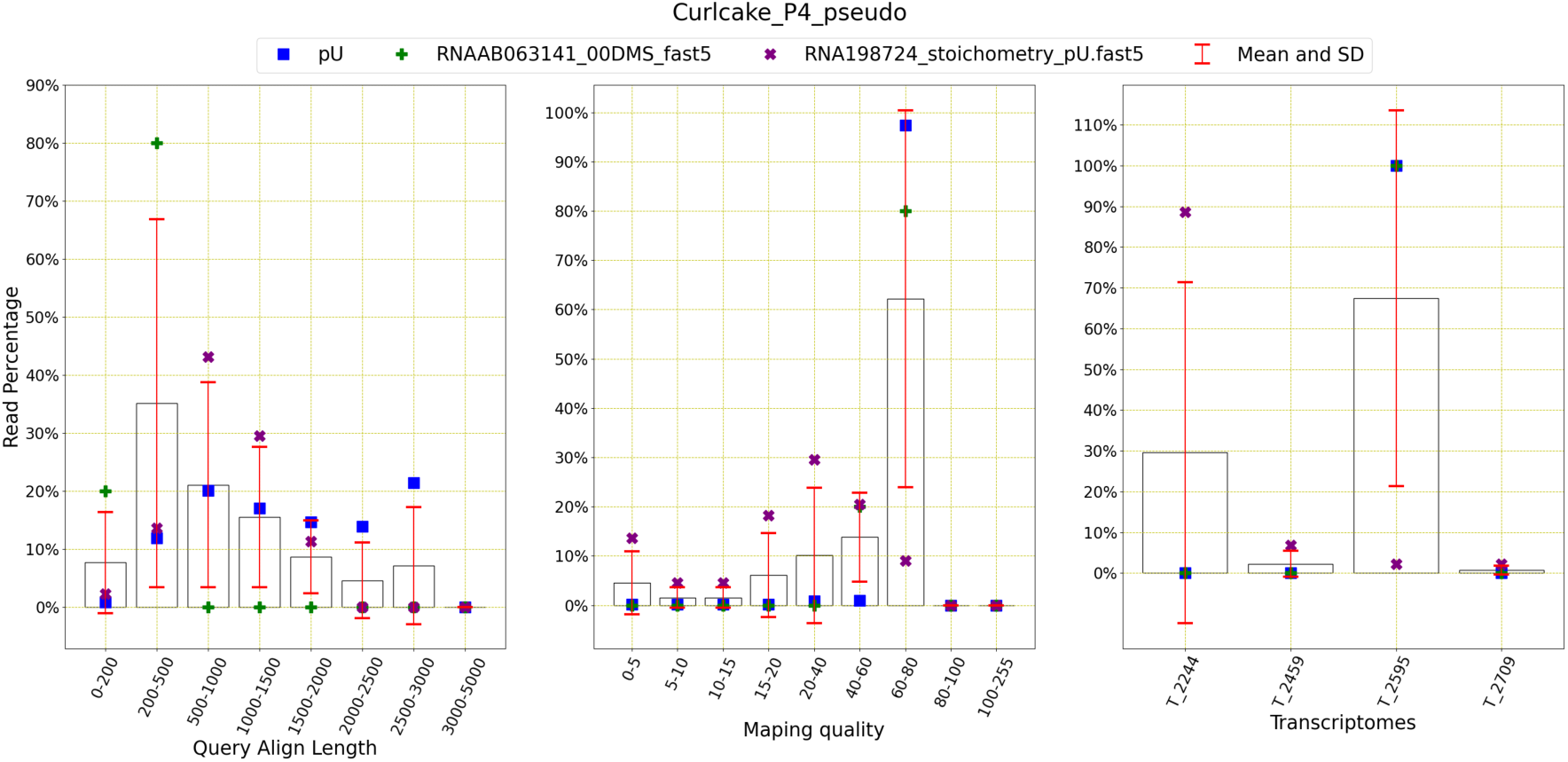
**Read distribution of Curlcake (**[9]**) pseudouridine methylation data:** This figure shows the distribution of the reads for query align length, mapping quality and transcriptomes. The average number of reads with query length under 200 is 1259 and over 200 is 138937. This means 99.10% of the reads have a query length of greater than 200. The average number of reads of mapping quality less than 20 is 1013 and over 20 is 139213. This shows that 99.28% of reads have a mapping quality higher than 20.

The read distribution increases for the higher query lengths and the mapping quality is also higher than 60 for most reads for three datasets. It can also be noticed that only one transcript appear in all the reads for these three datasets.

#### 3.3.2. IVT

This is the same artificially created reference sequence discussed before. For pseudouridine methylation, there are only two datasets presented in Jenjaroenpun et al. [28]. IVT_pseudoU_fast5 is the methylated data and IVT_normalU_fast5 is non-methylated data.

**Table 11.**
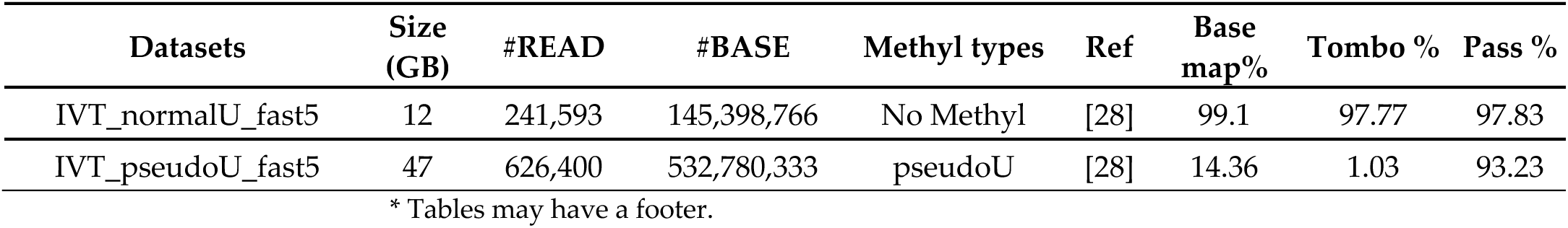
This table explains the datasets for IVT and pseudouridine (Ψ) methylation and shows size (in GB), the number of reads and bases, corresponding references and base mapping, Tombo success rates and pass rates (see Section 2)

The table shows that the methylated dataset has very low base mapping and Tombo rate, but a high pass rate, whereas the non-methylated dataset has a high score in all three aspects.

**Figure 20.**
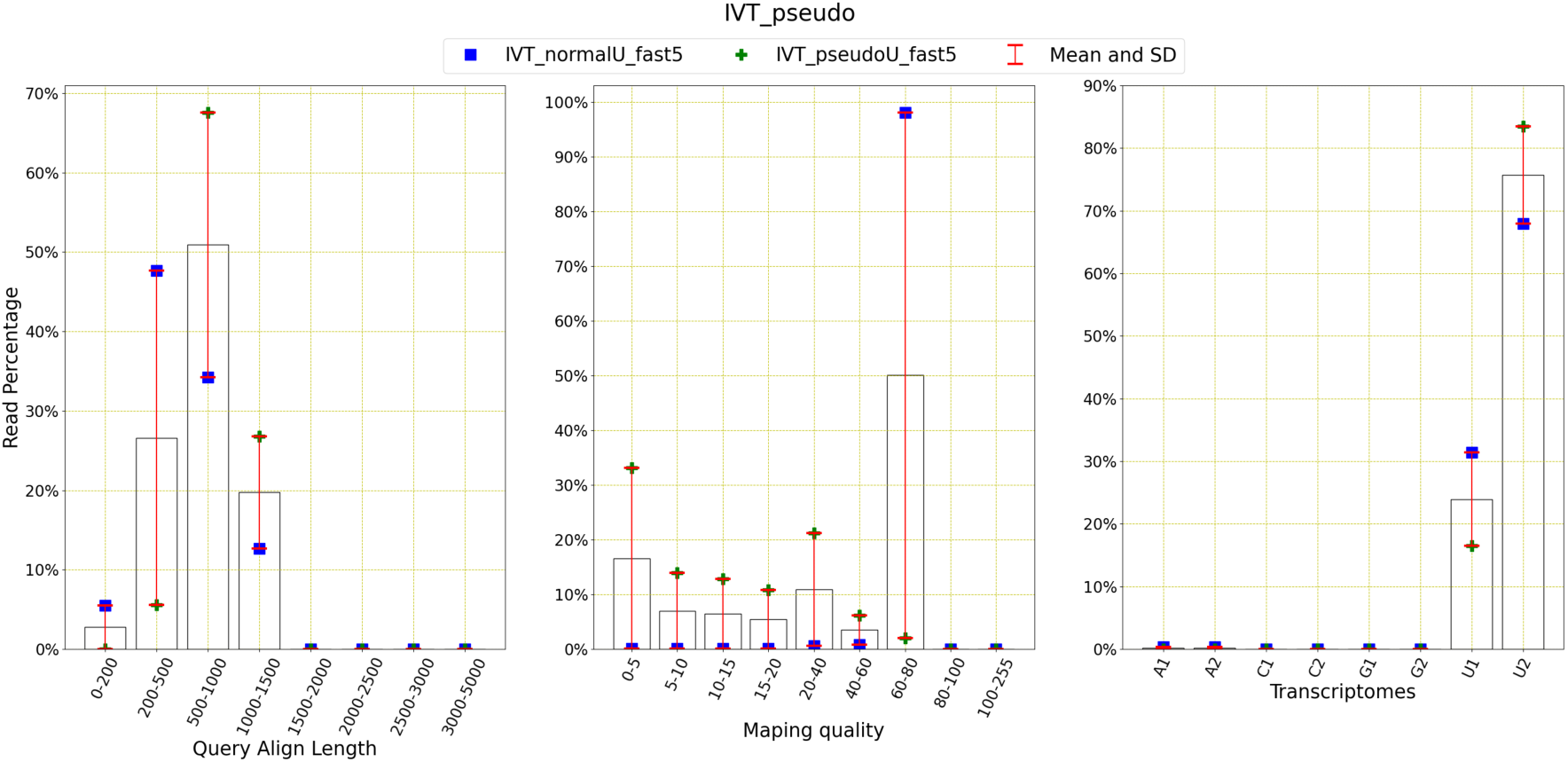
Read distribution of IVT pseudouridine methylation data: This figure shows the distribution of the reads for query align length, mapping quality and transcriptomes. The average number of reads with query length under 200 is 6626 and over 200 is 149262. This means 95.75% of the reads have a query length of greater than 200. The average number of reads of mapping quality less than 20 is 25501 and over 20 is 130387. This shows that 83.64% of reads have a mapping quality higher than 20.

The above figure shows that most of the reads contain a query length of 200 to 1000 for two datasets. The mapping quality higher than 60 is also prevalent in most of the reads. As IVT was laboratory made, only U1 and U2 are present among these two datasets.

#### 3.3.3. Yeast

Unlike the previous two, 23 separate datasets were prepared for pseudounidine methylation detection by Begik et al. [9]. These 23 datasets are grouped into seven different dataset types as shown in the table below.

**Table 12.** This table explains the datasets for yeast and pseudouridine (Ψ) methylation and shows size (in GB), the number of reads and bases, corresponding references and base mapping, Tombo success rates and pass rates (see Section 2)

| Datasets | Groups | Size (GB) | #READ | #BASE | Ref | Base map% | Tombo % | Pass % |
| --- | --- | --- | --- | --- | --- | --- | --- | --- |
| RNA100820 | mRNA | 114 | 1,310,518 | 767,456,221 | [9] | 95.29 | 87.31 | 90.24 |
| RNA140920 | mRNA | 108 | 1,420,228 | 796,995,869 | [9] | 93.98 | 84.19 | 87.59 |
| RNA290920 | mRNA | 64 | 2,529,350 | 1,565,409,944 | [9] | 94.66 | 86.52 | 88.78 |
| RNA598723 | mRNA | 94 | 2,982,805 | 2,187,527,617 | [9] | 96.75 | 92.24 | 93.22 |
| RNA323585 | WT_pU_KOs | 161 | 2,396 | 1,314,507 | [9] | 55.58 | 15.07 | 5.97 |
| RNA323586 | WT_pU_Kos | 443 | 7,410 | 1,168,419 | [9] | 50.61 | 7.50 | 8.91 |
| RNA323587 | WT_pU_Kos | 754 | 12,255 | 2,332,028 | [9] | 56.38 | 9.70 | 10.33 |
| RNA323588 | WT_pU_KOs | 16 | 288 | 81,509 | [9] | 64.93 | 10.07 | 4.17 |
| RNA814001 | WT_pU_KOs | 50 | 441,588 | 361,590,747 | [9] | 92.35 | 55.52 | 51.40 |
| RNA345944 | WT_vs_Xm_KOs | 122 | 1029,199 | 976,954,020 | [9] | 74.43 | 50.83 | 55.50 |
| RNA475631 | WT_vs_Xm_Kos | 26 | 387 | 221,386 | [9] | 21.11 | 7.75 | 3.62 |
| RNA475632 | WT_vs_Xm_Kos | 4.1 | 47,855 | 33,447,975 | [9] | 93.91 | 57.29 | 54.66 |
| RNA475633 | WT_vs_Xm_Kos | 1.5 | 20,922 | 15,344,035 | [9] | 58.89 | 18.02 | 5.31 |
| RNA475634 | WT_vs_Xm_KOs | 3.5 | 36,713 | 29,422,751 | [9] | 95.24 | 59.58 | 55.87 |
| RNA442567 | Translation_Frac | 12 | 114,241 | 97,705,756 | [9] | 88.07 | 57.70 | 57.12 |
| RNA927416 | Translation_Frac | 22 | 242,906 | 138,325,680 | [9] | 38.10 | 22.87 | 51.54 |
| RNA639991 | Oxidative_Stress | 4.6 | 45,434 | 37,916,410 | [9] | 81.01 | 63.15 | 69.51 |
| RNA873248 | Oxidative_Stress | 262 | 2,430,060 | 2,096,752,453 | [9] | 93.85 | 67.20 | 67.60 |
| RNA524356 | Nutrient_Temp | 103 | 1,247,129 | 720,654,683 | [9] | 74.75 | 54.22 | 61.12 |
| RNA564572 | Nutrient_Temp | 78 | 982,504 | 513,304,044 | [9] | 70.26 | 43.95 | 55.07 |
| RNA235628 | Nutrient_Temp | 178 | 2,105,360 | 1,262,485,686 | [9] | 77.79 | 50.15 | 55.51 |
| RNA113124 | Pus4_KO | 189 | 2,142,200 | 1,470,953,365 | [9] | 91.88 | 64.74 | 71.21 |
| RNA325982 | Pus4_KO | 32 | 493,931 | 179,101,531 | [9] | 78.46 | 39.73 | 42.95 |

The above table shows that there exists a huge variance between all these datasets and types. The following figure will explain the differences visually.

**Figure 21.**
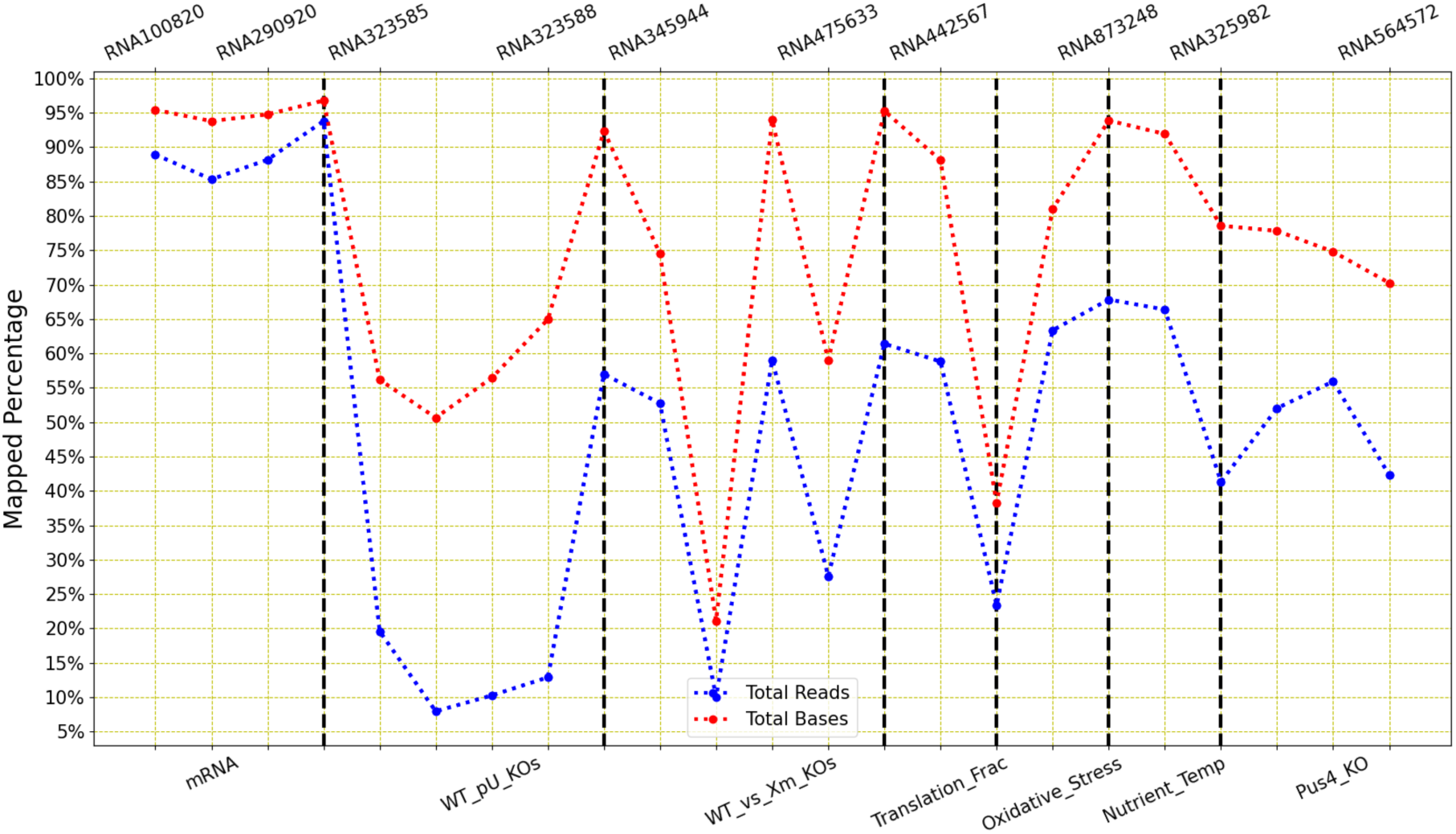
LongReadSum mapping percentage of reads and bases of yeast pseudouridine methylation data: This figure shows a huge deviation of mapping success rates among all these datasets. Here, some datasets types clearly performed better than others.

The figure shows that mRNA types performed better than the others as the mapping success rate is high (over 85%) for both reads and bases. Nutrient_Temp and Pus4_KO also performed decently as the ratio never drops below 40%. However, the other types contain some datasets which performed really poorly as the ratio went down to as low as 10%.

The figure also explains that there is no certain pattern is present among these dataset types. The read distribution for query length is high between 200 and 1000 for mRNA, Oxidative_Stress and Nutrient_Temp. Datasets such as mRNA contain reads with mapping quality higher than 60, however Oxidative_Stress contains reads with reads whose mapping quality is mostly less than 20. The transcriptome distribution is also not equal for all data types. For mRNA, it is fairly equal, however only chrX mostly appears in Oxidative_Stress.

mRNA dataset clearly stands out from the rest of the dataset types here.

### 3.4. m5C and hm5C and FormylC

#### 3.4.1. Curlcake ([9])

This artificially made curlcake reference sequence is prepared by Begik et al. [9], and there are only two datasets which uses this reference sequence for m5C and hm5C methylation.

**Table 13.** This table explains the datasets for curlcake ([9]) and m5C and hm5C and FormylC methylation and shows size (in GB), the number of reads and bases, corresponding references and base mapping, Tombo success rates and pass rates (see Section 2)

| Datasets | Size (GB) | #READ | #BASE | Methyl types | Reference | Base map% | Tombo % | Pass % |
| --- | --- | --- | --- | --- | --- | --- | --- | --- |
| RNA080420191_5hmC | 16 | 111,015 | 113,763,443 | hm5C | [9] | 83.74 | 55.27 | 54.28 |
| RNA010220191_m5C | 30 | 415,453 | 224,786,171 | m5C | [9] | 70.59 | 42.11 | 61.82 |

\* Tables may have a footer.

They vary in their size and number of reads and bases.

**Figure 22.**
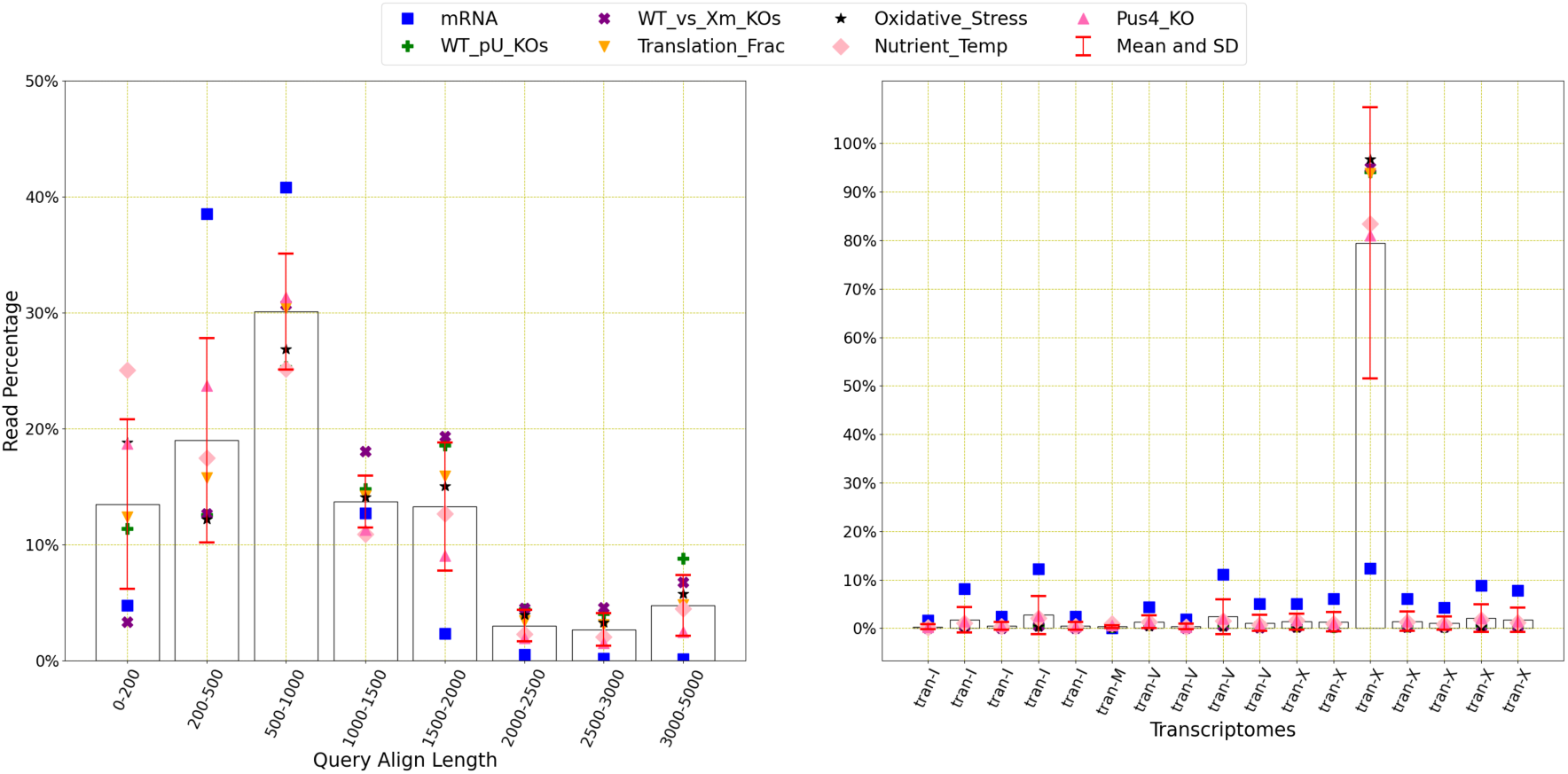
Read distribution of yeast pseudouridine methylation data: This figure shows the distribution of the reads for query align length, mapping quality and transcriptomes. The average number of reads with query length under 200 is 152981 and over 200 is 850804. This means 84.76% of the reads have a query length of greater than 200. The average number of reads of mapping quality less than 20 is 658225 and over 20 is 345886. This shows that 34.45% of reads have a mapping quality higher than 20.

**Figure 23.**
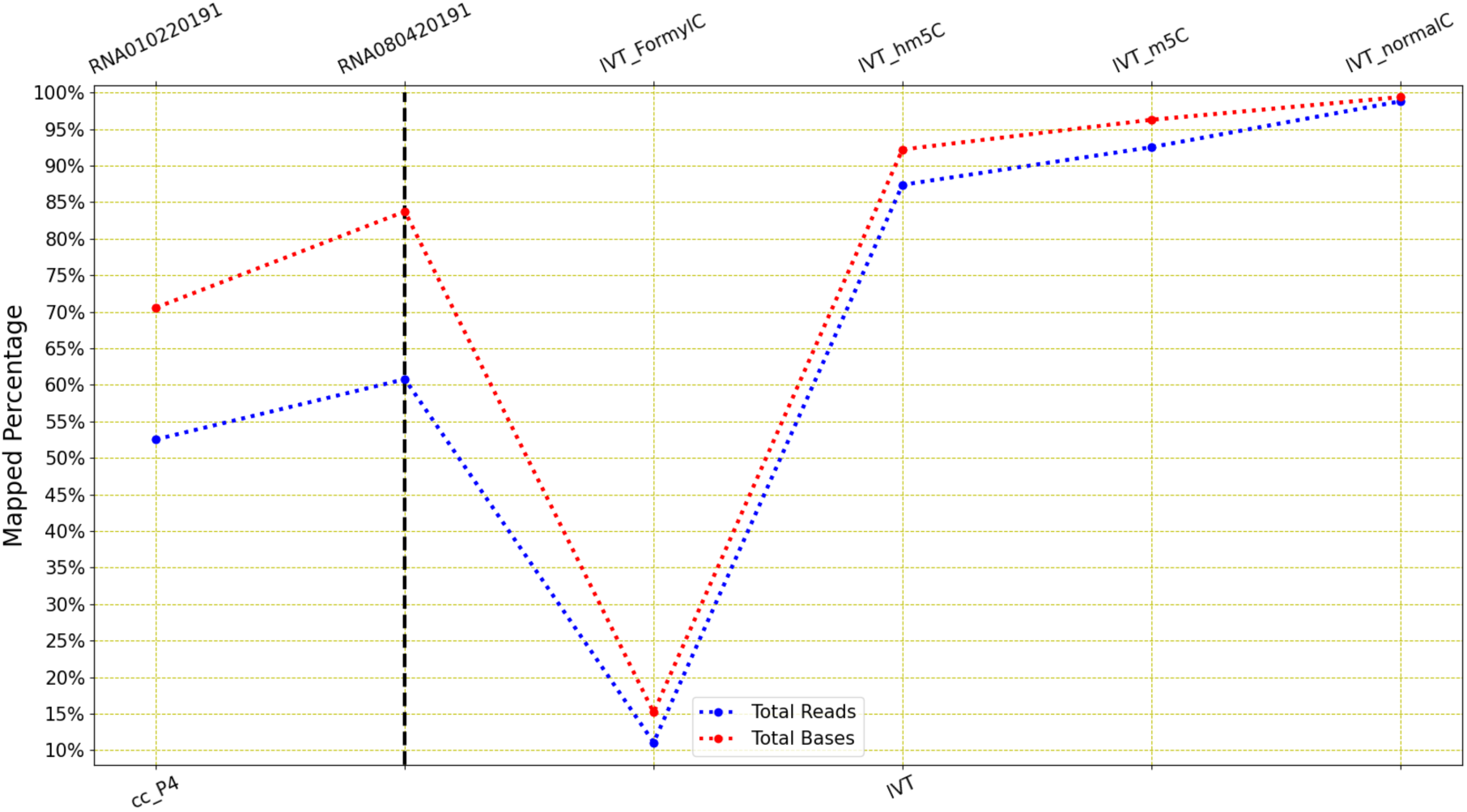
LongReadSum mapping percentage of reads and bases of curlcake. [9] **and IVT m5C and hm5C methylation data:** This figure shows a small variance of mapping success rates among these two datasets. For IVT, all other three datasets have performed well except FormylC.

The mapping success rates for the reads lie between 70 to 80% and for the bases, it lie between 50 to 60%. These statistics are not as high as some datasets discussed previously.

The above figure shows that m5C dataset contains query lengths ranging from 200 to 3000 and they are equally distributed. However, most of the reads in 5hmc dataset has the query length of 500 to 1000. Both datasets have the mapping quality greater than 60. This artificially created reference sequence has four transcripts and all of them are present in the reads for both datasets.

**Figure 24.**
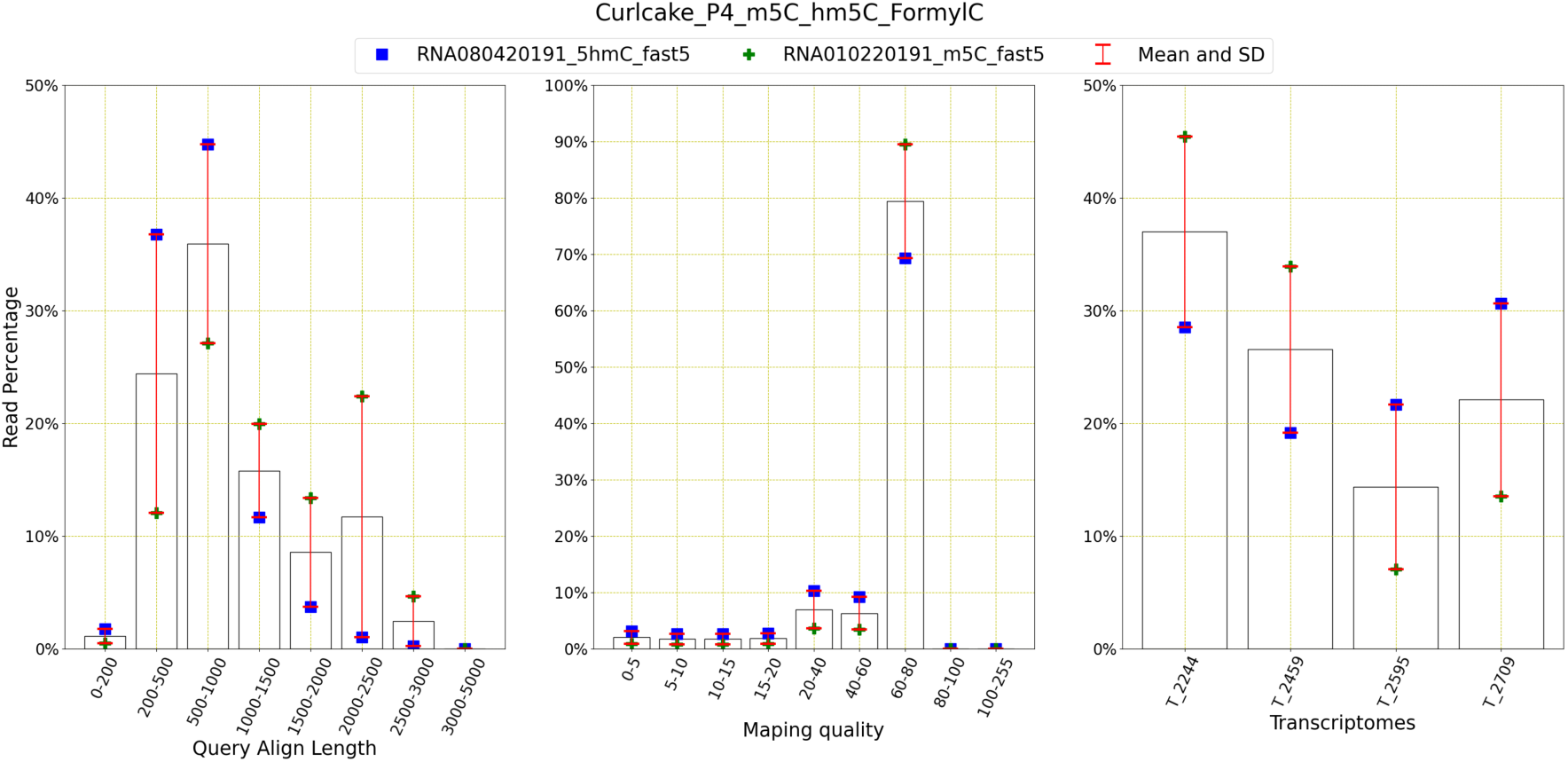
Read distribution of IVT m6A and m5C methylation data: This figure shows the distribution of the reads for query align length, mapping quality and transcriptomes. The average number of reads with query length under 200 is 2107 and over 200 is 140557. This means 98.52% of the reads have a query length of greater than 200. The average number of reads of mapping quality less than 20 is 13481 and over 20 is 129184. This shows that 90.55% of reads have a mapping quality higher than 20.

#### 3.4.2 IVT

The artificially created IVT reference sequence is also developed for m5C and hm5C and FormylC methylations. There are four datasets which are prepared for this methylation on cytosine (C) nucleotide. Three of those datasets are methylated and one is nonmethylated as described in the following table.

**Table 14.** This table explains the datasets for IVT and m5C and hm5C and FormylC methylation and shows size (in GB), the number of reads and bases, corresponding references and base mapping, Tombo success rates and pass rates (see Section 2)

| Datasets | Size | #READ | #BASE | Methyl types | Ref | Base map% | Tombo % | Pass % |
| --- | --- | --- | --- | --- | --- | --- | --- | --- |
| IVT_hm5C_fast5 | 3.7 | 61,304 | 36,232,972 | hm5C | [28] | 92.22 | 74.04 | 90.89 |
| IVT_FormylC_fast5 | 45 | 663,193 | 530,130,955 | FormylC | [28] | 15.19 | 1.59 | 96.99 |
| IVT_m5C_fast5 | 37 | 573,674 | 413,053,134 | m5C | [28] | 96.28 | 79.27 | 93.34 |
| IVT_normalC_fast5 | 29 | 452,806 | 332,024,954 | X | [28] | 99.4 | 97.57 | 96.89 |

\* Tables may have a footer.

The table shows that even though the size and the number of reads vary for these datasets, the base mapping, Tombo and pass success rates are high except IVT_FormylC_fast dataset.

The performances for all four datasets are shown here and as noted previously for the other datasets, the non-methylated datasets perform better than methylated datasets.

**Figure 25.**
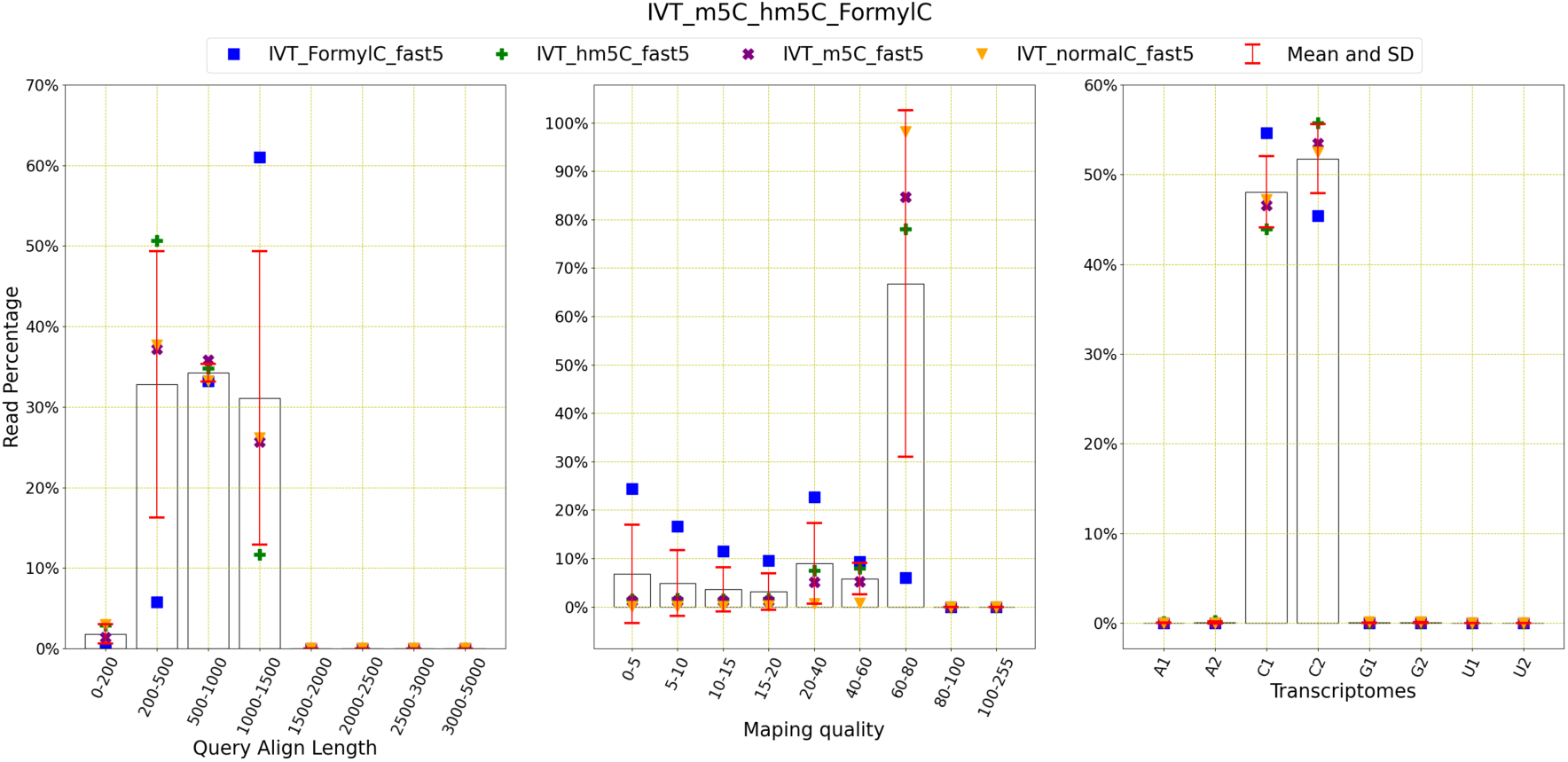
Read distribution of IVT m5C, hm5C and FormylC methylation data: This figure shows the distribution of the reads for query align length, mapping quality and transcriptomes. The average number of reads with query length under 200 is 5556 and over 200 is 271650. This means 98% of the reads have a query length of greater than 200. The average number of reads of mapping quality less than 20 is 19390 and over 20 is 257816. This shows that 93.01% of reads have a mapping quality higher than 20.

**Table 15.** This table explains the datasets for E Coli and Nm and pseudouridine methylation and shows size (in GB), the number of reads and bases, corresponding references and base mapping, Tombo success rates and pass rates (see Section 2)

| Datasets | Size (GB) | #READ | #BASE | Reference | Base map% | Tombo % | Pass % |
| --- | --- | --- | --- | --- | --- | --- | --- |
| SRX8386931 | 165 | 1,517,105 | 1,365,411,124 | [29] | 96.13 | 68.42 | 77.08 |
| SRX8386932 | 157 | 1,416,308 | 1,466,427,832 | [29] | 77.29 | 87.40 | 89.98 |

The above figure shows that all four datasets have reads with query length ranging from 200 to 3000 and most of the reads have mapping quality greater than 60. IVT is the sequence which contain transcripts such as C1 and C2 specifically made to detect C methylations, thus reads contain only those two transcripts.

### 3.5. Nm and pseudouridine

#### 3.5.1. E Coli

ONT datasets for Nm and pseudouridine methylations using E Coli reference is found in Stephenson et al. [29].

The above table shows that datasets doesn’t vary a lot in their size and number of reads, however the base mapping, Tombo and pass success rates do vary.

The above figure shows a large variance between these two datasets. One performed better than the other in terms of both reads and bases with a larger variance and the other has both read and base mapping rate of around 77%.

**Figure 26.**
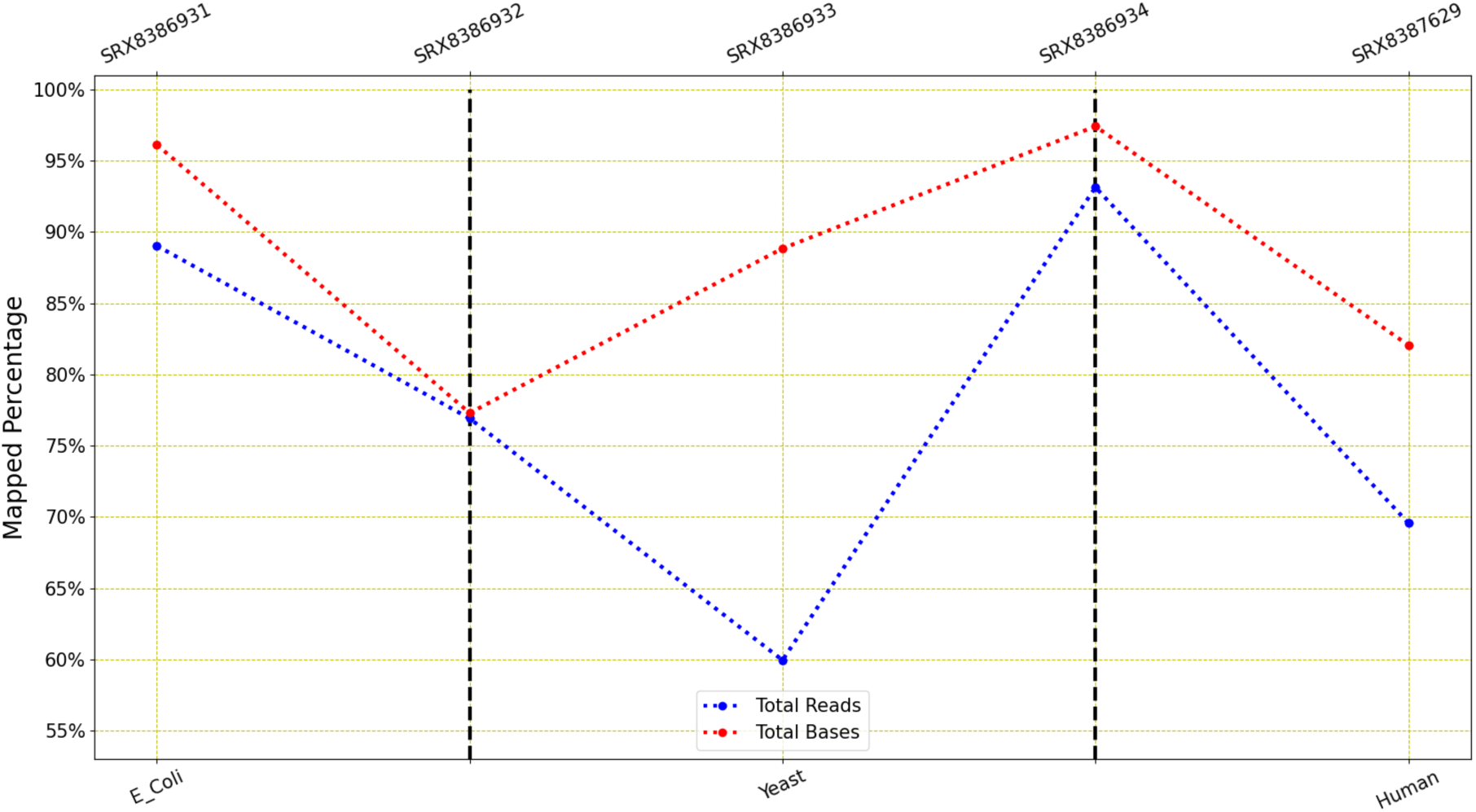
LongReadSum mapping percentage of reads and bases of E Coli and yeast Nm and pseudouridine methylation data: This figure shows that the mapping success rate for these two datasets vary a lot for both methylations.

Both datasets contain reads whose query length mostly lie between 1000 and 2000. The mapping quality for both is less than 20 and there is only one transcript present in E Coli sequence and all reads are mapped to that transcript only.

#### 3.5.2. Yeast

ONT datasets for Nm and pseudouridine methylations using yeast reference is found in Stephenson et al. [29] as well.

The base mapping and pass success rates are high for both datasets, however the Tombo rates are not so high.

**Figure 27.**
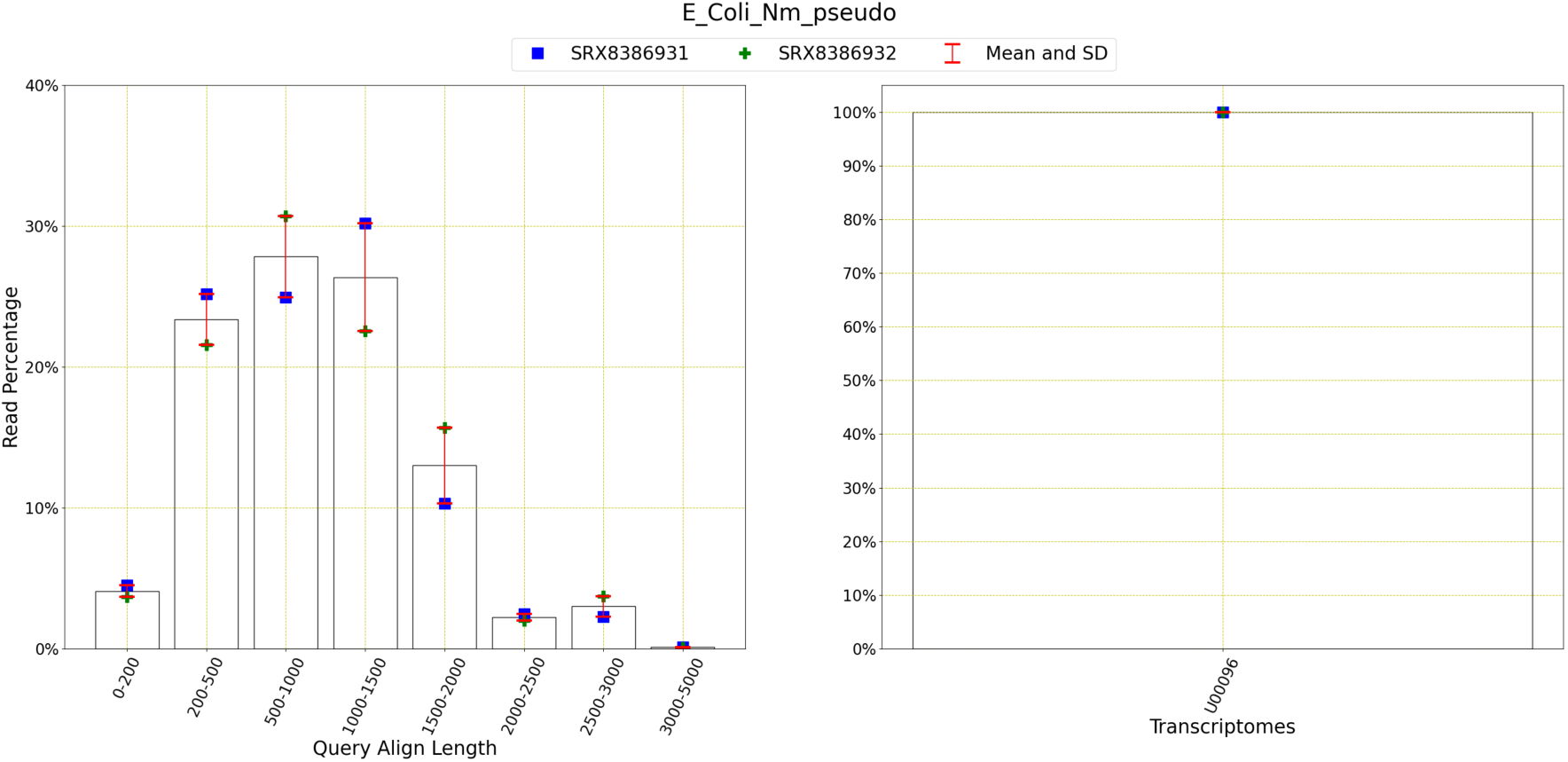
Read distribution of IVT m6A and m5C methylation data: This figure shows the distribution of the reads for query align length, mapping quality and transcriptomes. The average number of reads with query length under 200 is 286258 and over 200 is 6644460. This means 95.87% of the reads have a query length of greater than 200. The average number of reads of mapping quality less than 20 is 6742505 and over 20 is 195506. This shows that 2.82% of reads have a mapping quality higher than 20.

The above figure shows that base mapping ratios for both datasets are high, but the read mapping ratio is high for SRX8386934, but low for SRX8386933.

The above figure shows that both datasets contain reads with query lengths ranging from 0 to 3000. Most of the reads in SRX8386933 have the query lengths ranging from 0 to 1000 and most of the reads in SRX8386934 have the query length of 1000-2000. The mapping quality of reads in both datasets is less than 20. Yeast contain 17 transcripts and only chrXII is present abundantly among the reads.

#### 3.5.3. Human

There is only one ONT dataset for Nm and pseudouridine methylations

**Table 16.** This table explains the datasets for yeast and Nm and pseudouridine methylation and shows size (in GB), the number of reads and bases, corresponding references and base mapping, Tombo success rates and pass rates (see Section 2)

| Datasets | Size (GB) | #READ | #BASE | Reference transcript | Base map% | Tombo % | Pass % |
| --- | --- | --- | --- | --- | --- | --- | --- |
| SRX8386933 | 132 | 1,514,181 | 880,493,782 | [29] | 88.85 | 75.89 | 95.82 |
| SRX8386934 | 174 | 1,529,837 | 1,645,169,328 | [29] | 97.42 | 59.55 | 93.07 |

using human reference in Stephenson et al. [29].

The above table shows a high value for all three success scores.

**Figure 28.**
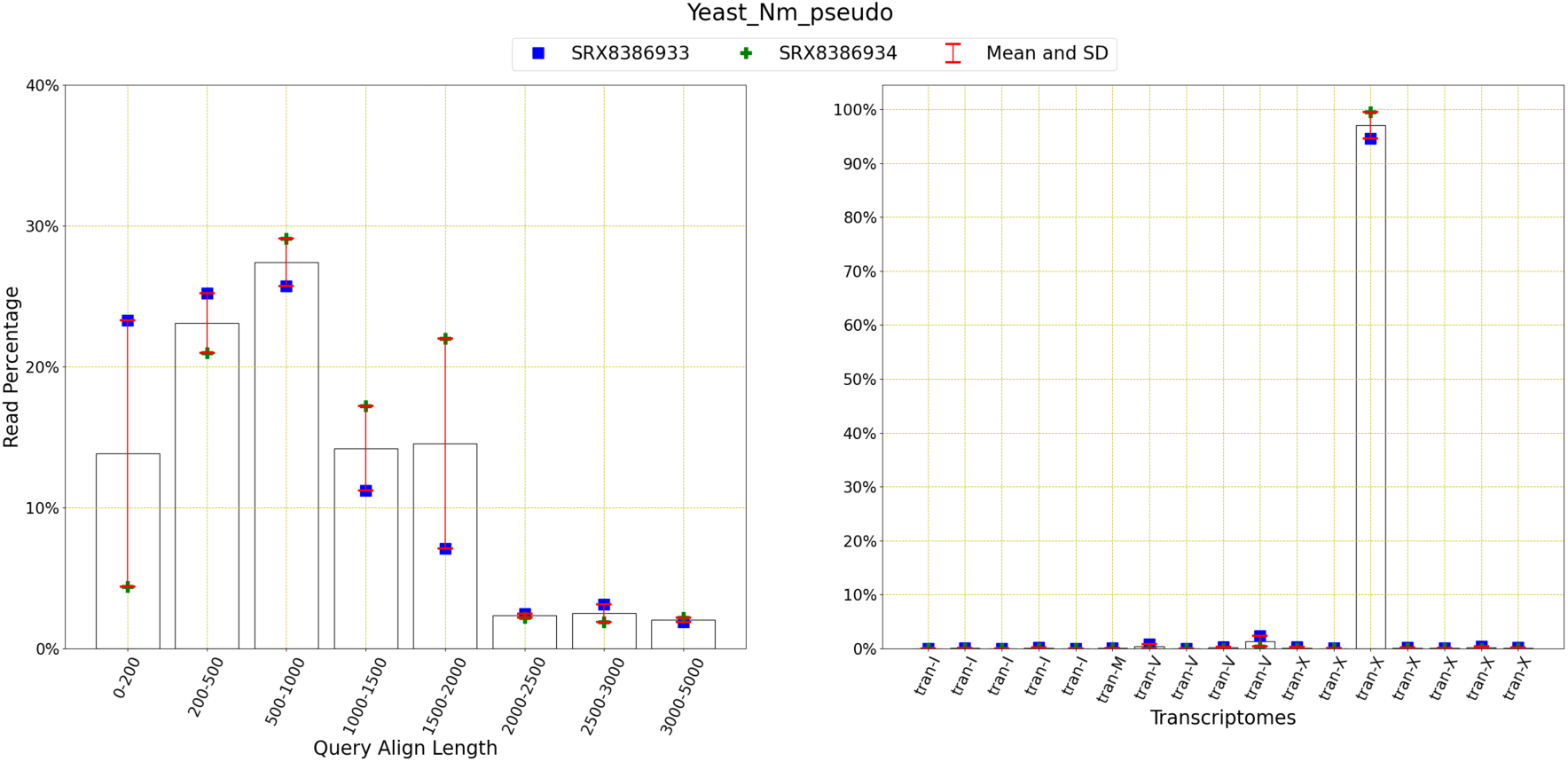
Read distribution of IVT m6A and m5C methylation data: This figure shows the distribution of the reads for query align length, mapping quality and transcriptomes. The average number of reads with query length under 200 is 282498 and over 200 is 2036074. This means 87.82% of the reads have a query length of greater than 200. The average number of reads of mapping quality less than 20 is 2296297 and over 20 is 71480. This shows that 3.02% of reads have a mapping quality higher than 20.

The figure shows that most of the reads in the dataset have the query length of 200 to 500 and their mapping quality is less than 20. Only three among all the other transcripts in human transcriptomes are present in the reads for this dataset.

**Figure 29.**
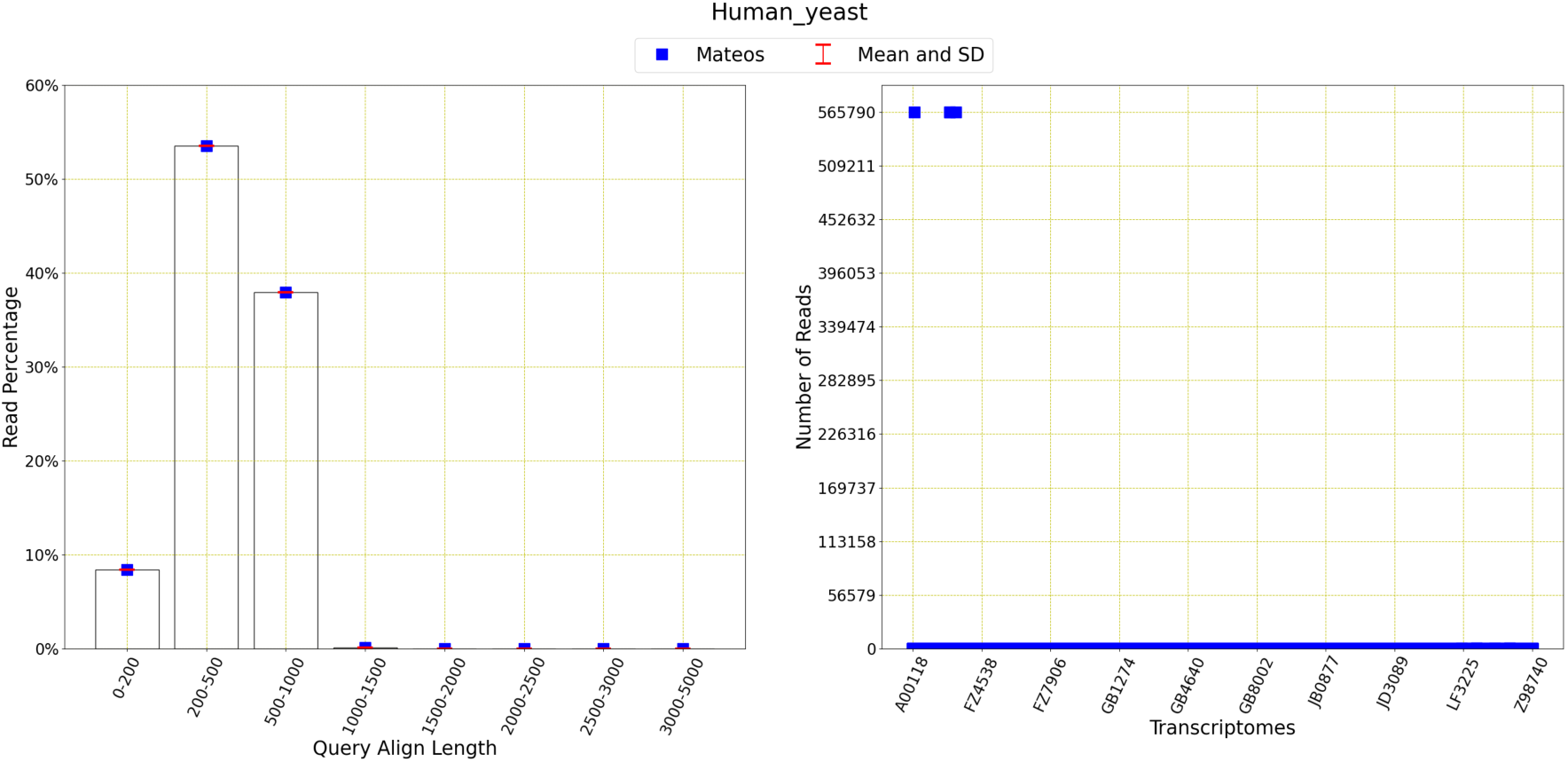
Read distribution of IVT m6A and m5C methylation data: This figure shows the distribution of the reads for query align length, mapping quality and transcriptomes. The average number of reads with query length under 200 is 190761 and over 200 is 2076358. This means 91.59% of the reads have a query length of greater than 200. The average number of reads of mapping quality less than 20 is 2266940 and over 20 is 179. This shows that 0.01% of reads have a mapping quality higher than 20.

### 3.6. Inosine, m1A and m7G

#### 3.6.1. IVT

The ONT datasets for Inosine, m1A and m7G methylations using artificially made IVT sequence is also found in Jenjaroenpun et al. [28]. There are five datasets and there of them are methylated and two are non-methylated.

**Table 17.** This table explains the datasets for human and Nm and pseudouridine methylation and shows size (in GB), the number of reads and bases, corresponding references and base mapping, Tombo success rates and pass rates (see Section 2)

| Datasets | Size (GB) | #READ | #BASE | Reference | Base map% | Tombo % | Pass % |
| --- | --- | --- | --- | --- | --- | --- | --- |
| SRX8387629 | 54 | 806,550 | 358,366,721 | [29] | 82.05 | 92.23 | 60.24 |

The table shows a strickiing difference in success rates between these datasets. All three success rates such as base mapping, Tombo and pass rates are high for the nonmethylated datasets and they are exteremly low for IVT_Inosine_fast5 and IVT_m7G_fast5.

**Figure 30.**
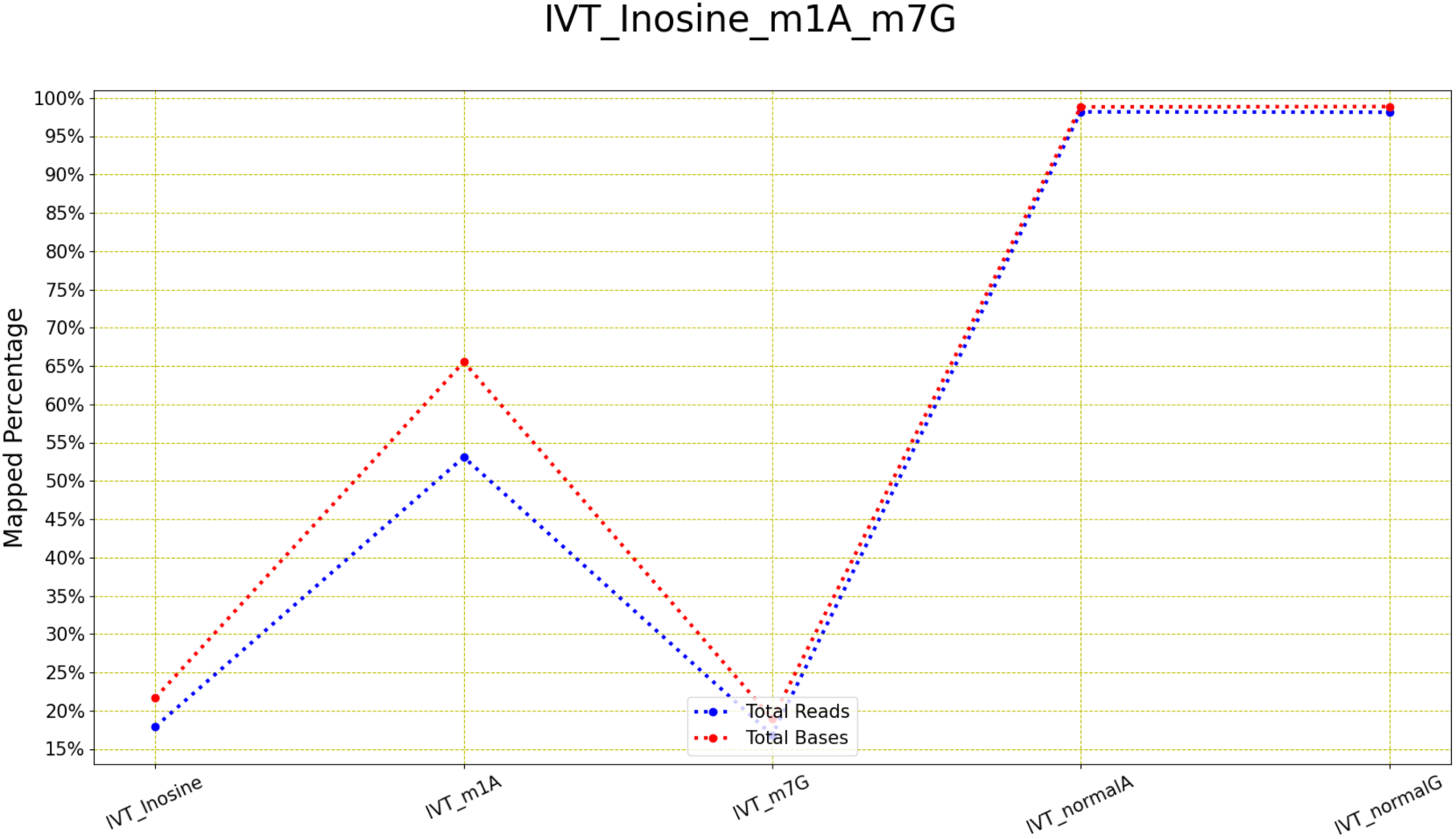
LongReadSum mapping percentage of reads and bases of IVT Inosine, m1A and m7G methylation data: This figure shows that the mapping success rate for all these five datasets vary a lot.

**Table 18.** This table explains the datasets for IVT and Inosine, m1A and m7G methylation and shows size (in GB), the number of reads and bases, corresponding references and base mapping, Tombo success rates and pass rates (see Section 2)

| Datasets | Size (GB) | #READ | #BASE | Methyl types | Ref | Base map% | Tombo % | Pass % |
| --- | --- | --- | --- | --- | --- | --- | --- | --- |
| IVT_Inosine_fast5 | 38 | 811,953 | 685,427,011 | Inosine | [28] | 21.68 | 8.1 | 53.31 |
| IVT_m1A_fast5 | 8.8 | 187,199 | 93,501,694 | m1A | [28] | 65.53 | 27.18 | 77.95 |
| IVT_m7G_fast5 | 0.4 | 6,413 | 4,624,868 | m7G | [28] | 18.93 | 25.78 | 2.84 |
| IVT_normalA_fast5 | 19 | 383,209 | 238,341,386 | No Methyl | [28] | 98.82 | 96.63 | 96.95 |
| IVT_normalG_fast5 | 41 | 703,754 | 492,403,961 | No Methyl | [28] | 98.87 | 96.21 | 95.68 |

\* Tables may have a footer.

The similar trend is also found for the base mapping ratios as well. The nonmethylated datasets performed better than the methylated datasets. The difference between the reads and bases are higher for IVT_m1A_fast5, which performed the best among all the other methylated datasets, than all the other datasets.

**Figure 31.**
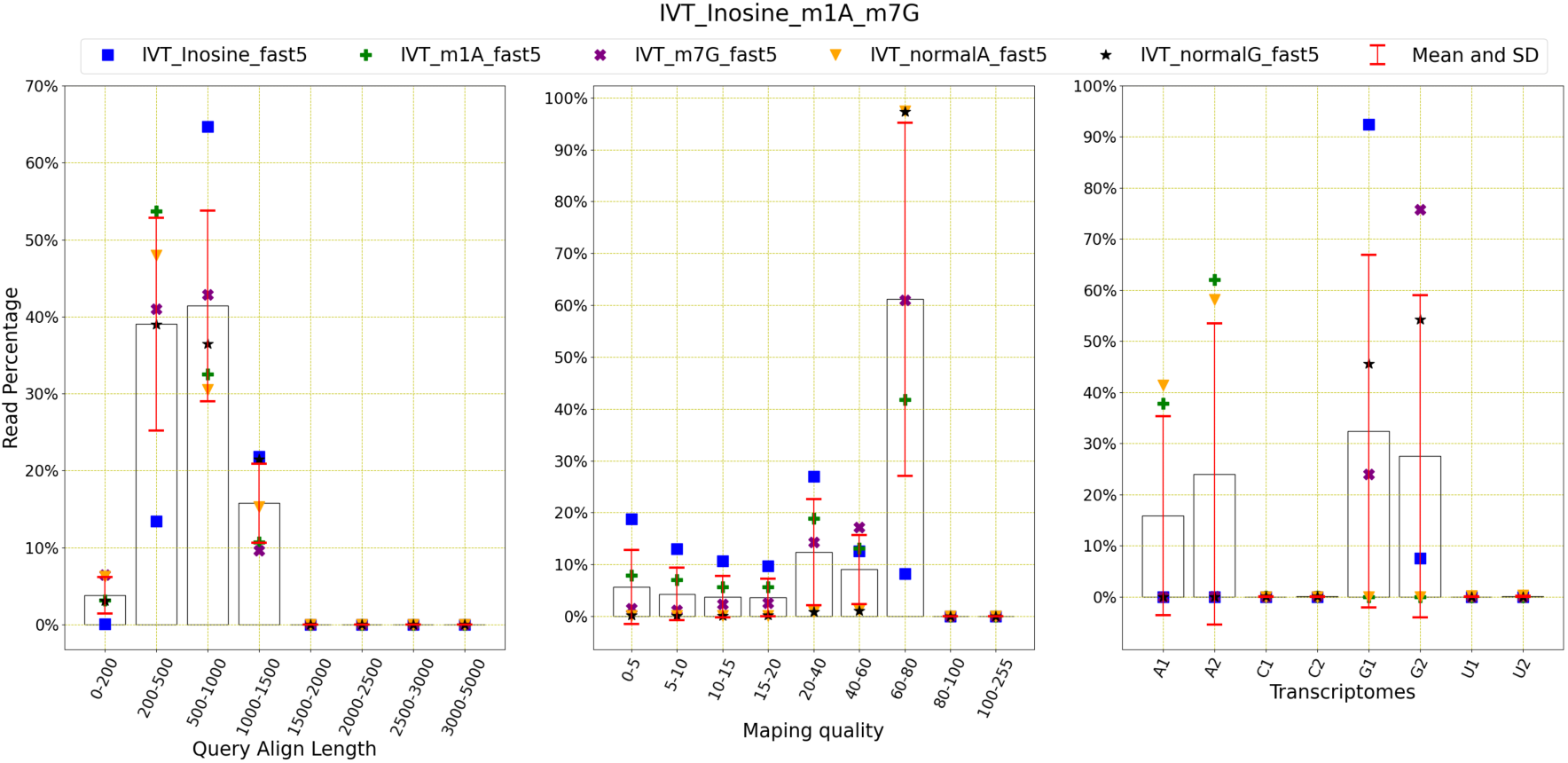
Read distribution of IVT m6A and m5C methylation data: This figure shows the distribution of the reads for query align length, mapping quality and transcriptomes. The average number of reads with query length under 200 is 9648 and over 200 is 254730. This means 96.35% of the reads have a query length of greater than 200. The average number of reads of mapping quality less than 20 is 21861 and over 20 is 242516. This shows that 91.73% of reads have a mapping quality higher than 20.

The above figure shows that non-methylated datasets have reads whose query align length is mainly 200 to 500. The other datasets have reads with query lengths varying from 0 to 2000. The mapping quality has been greater than 60 for the non-methylated data and less than 40 for the methylated data. All transcripts except C1, C2, U1 and U2 appear in the reads for these five datasets.

**Figure 32.**
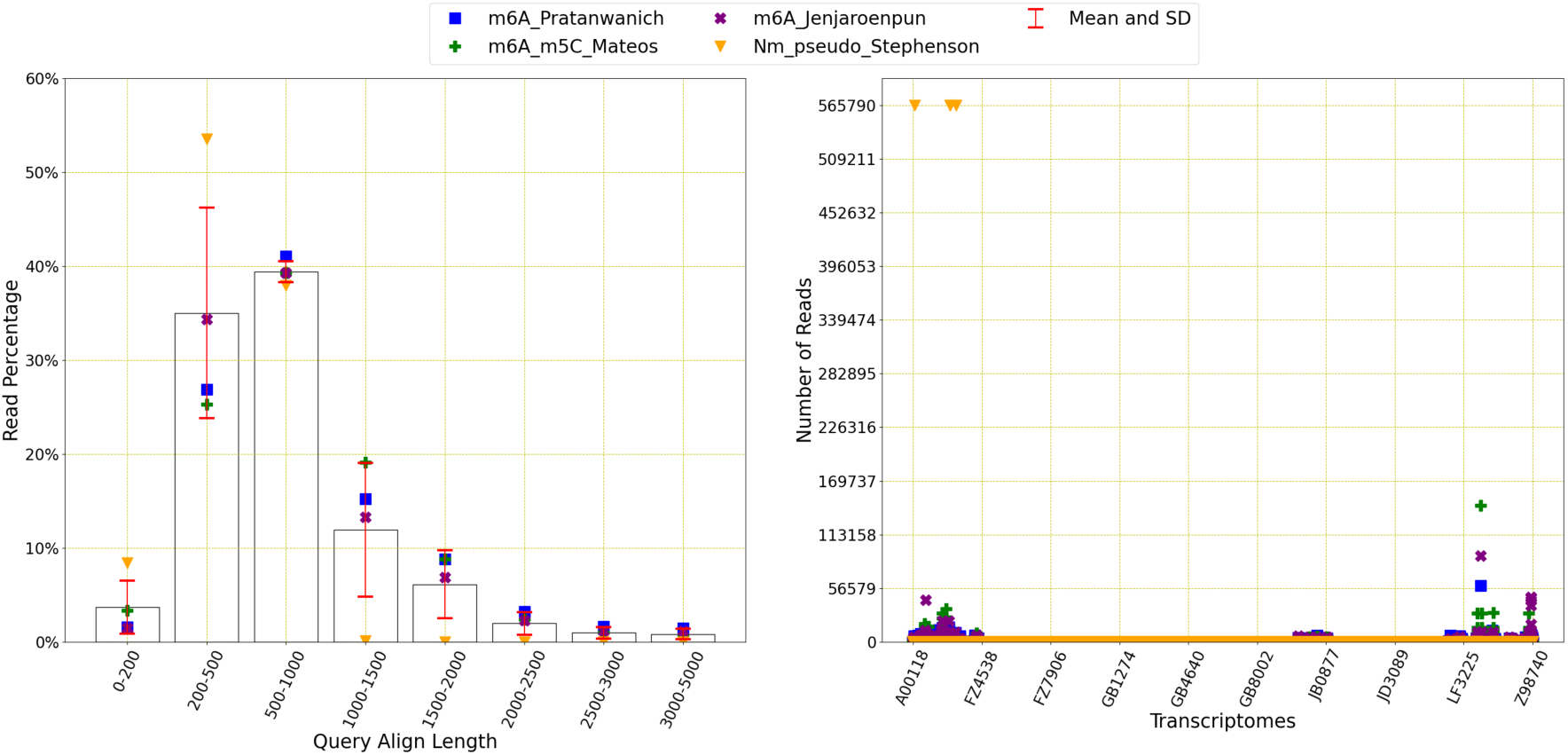
Read distribution of human data: This figure shows the distribution of the reads for query align length, mapping quality and transcriptomes. The average number of reads with query length under 200 is 132125 and over 200 is 4824960. This means 97.33% of the reads have a query length of greater than 200. The average number of reads of mapping quality less than 20 is 4909752 and over 20 is 104682. This shows that 2.09% of reads have a mapping quality higher than 20.

## 4. Datasets from various reference transcriptomes

We have also organised the datasets according to each reference transcriptome. They are discussed below.

### 4.1. Human

Human transcriptome data has been found in four sources which are Pratanwanich et al. [20], Mateos et al. [27], Jenjaroenpun et al. [28] and Stephenson et al. [29].

The above figure shows that query lengths do not follow a certain pattern, but most of the reads have the mapping quality of less than 20. Not all transcripts are present in the reads.

### 4.2. Mouse

ONT datasets appear from three different sources ([27], [28], [10]) and two methyaltion types (m6A and m5C).

The figure shows that most of the reads contain the query length between 200 and 1000, with mapping quality less than 5. Among all 586,852 transcripts, only a few appear among the reads.

**Figure 33.**
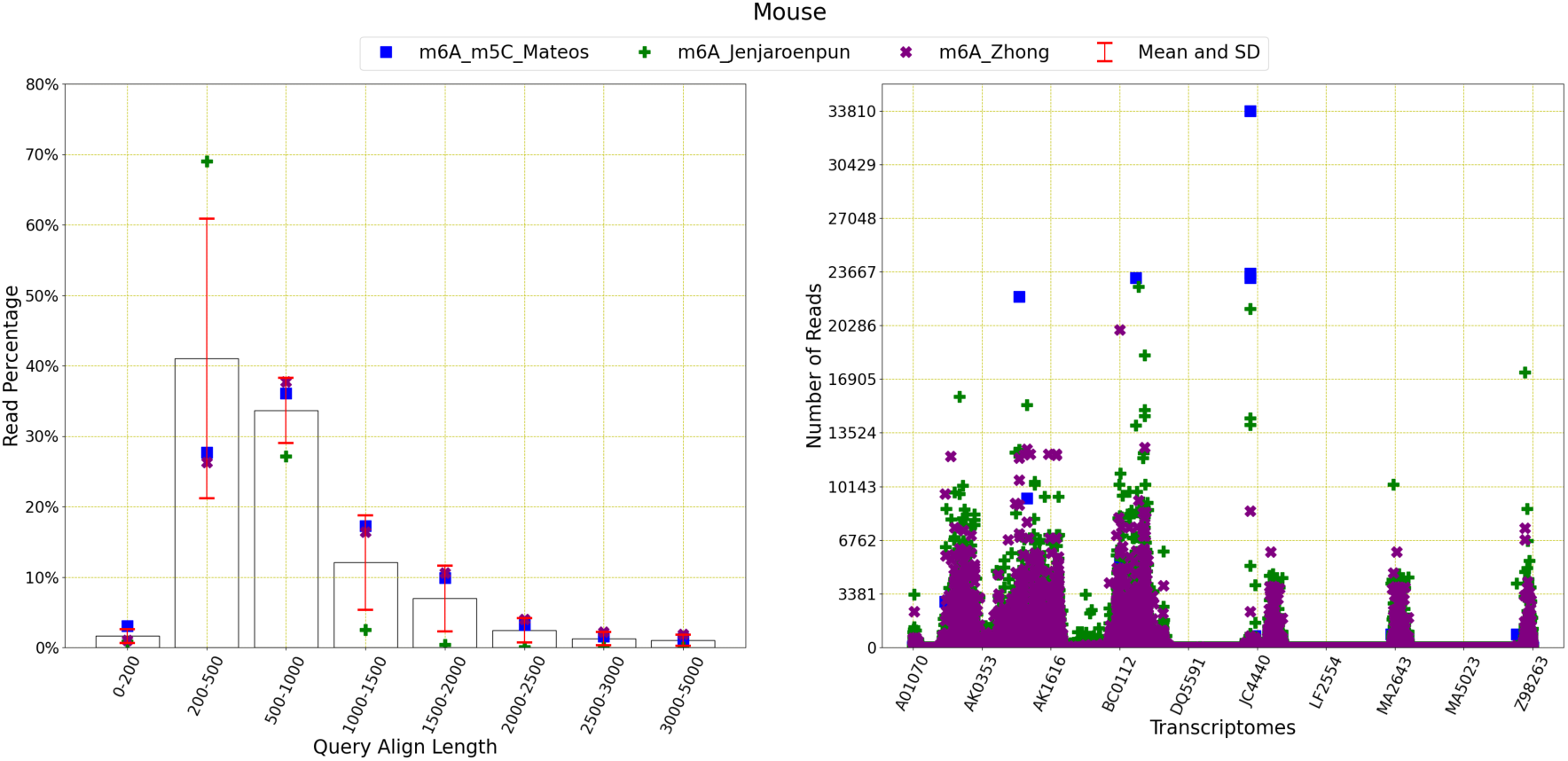
Read distribution of mouse data: This figure shows the distribution of the reads for query align length, mapping quality and transcriptomes. The average number of reads with query length under 200 is 66334 and over 200 is 6212872. This means 98.94% of the reads have a query length of greater than 200. The average number of reads of mapping quality less than 20 is 6244513 and over 20 is 86478. This shows that 1.37% of reads have a mapping quality higher than 20.

### 4.3. Curlcake ([24])

This is already been explained in Section 3.1.5 as there is only one dataset containing this reference sequence.

### 4.4. Curlcake ([9])

This artificially made curlcake reference sequence is developed by Begik et al. [9].

There are three types of methylations and the read distribution is shown below.

**Figure 34.**
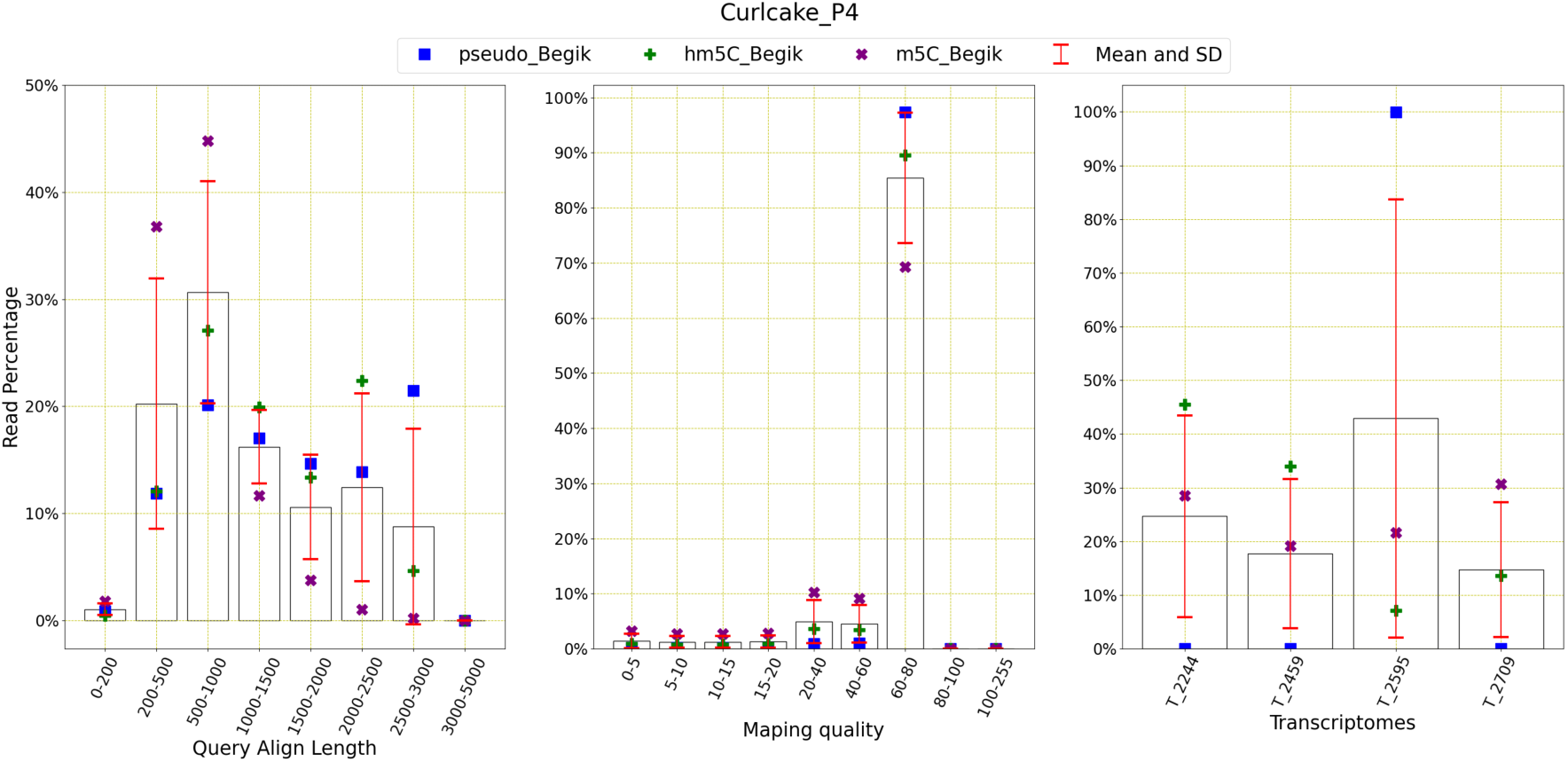
**Read distribution of Curlcake (**[9]**) data:** This figure shows the distribution of the reads for query align length, mapping quality and transcriptomes. The average number of reads with query length under 200 is 1824 and over 200 is 140017. This means 98.71% of the reads have a query length of greater than 200. The average number of reads of mapping quality less than 20 is 9325 and over 20 is 132527. This shows that 93.43% of reads have a mapping quality higher than 20.

Here, most of the reads have the query lengths vary between 200 to 3000 and the mapping quality is higher than 60. All four transcripts appear almost equally among the reads from different methylation types.

### 4.5. IVT

This artificially created IVT reference sequence is found only in Jenjaroenpun et al. [28]. There are multiple methylation types are present as shown in the following figure.

**Figure 35.**
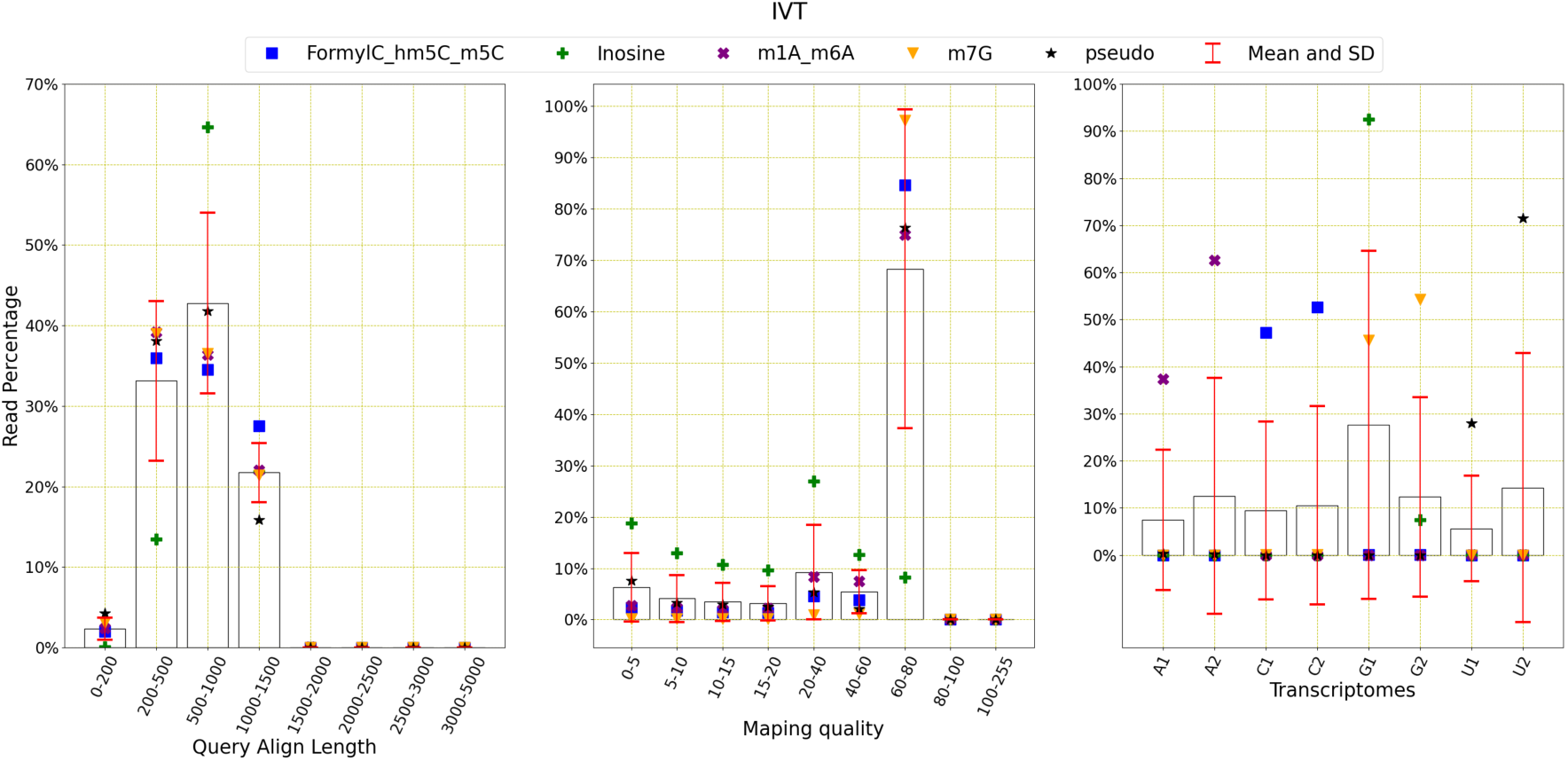
Read distribution of IVT data: This figure shows the distribution of the reads for query align length, mapping quality and transcriptomes. The average number of reads with query length under 200 is 7322 and over 200 is 292942. This means 97.56% of the reads have a query length of greater than 200. The average number of reads of mapping quality less than 20 is 35302 and over 20 is 264961. This shows that 88.24% of reads have a mapping quality higher than 20.

The figure shows that most reads contain query align length of 0 to 1500 and the mapping quality is mostly higher than 60. All eight transcriptomes appear almost equally among all reads from five different types of methylations.

### 4.6. Yeast

There are three methylations (m6A, pseudouridine and Nm) from four sources ([24], [9], [28], [29]) for yeast.

**Figure 36.**
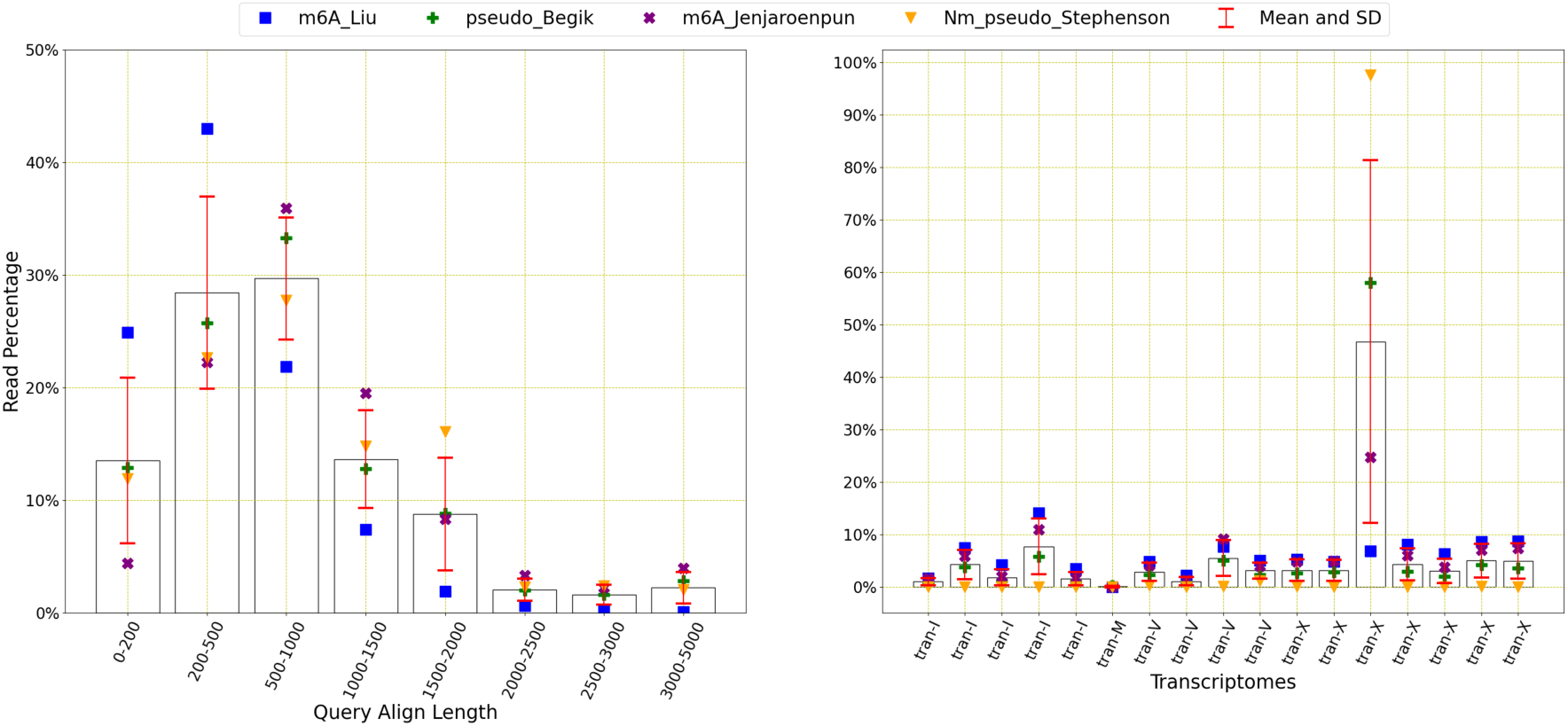
Read distribution of yeast data: This figure shows the distribution of the reads for query align length, mapping quality and transcriptomes. The average number of reads with query length under 200 is 147,053 and over 200 is 1,177,333. This means 88.90% of the reads have a query length of greater than 200. The average number of reads of mapping quality less than 20 is 851937 and over 20 is 519716. This shows that 37.89% of reads have a mapping quality higher than 20.

The figure shows that most of the reads from all four datasets have the query lengths between 0 and 2000 with variations, on average. Mapping quality is found either between 0 and 5 or 60 and 80, showing the presence of both good and bad reads. It is to be noted that Liu et al. dataset [24] contain mostly good reads of mapping quality greater than 60. Among all 17 transcriptomes, chrXII has appeared the most in all reads.

### 4.7. E Coli

The reference sequence e coli has been used for two methylation types, such as m6A, Nm and pseudouridine coming from two different sources.

**Figure 37.**
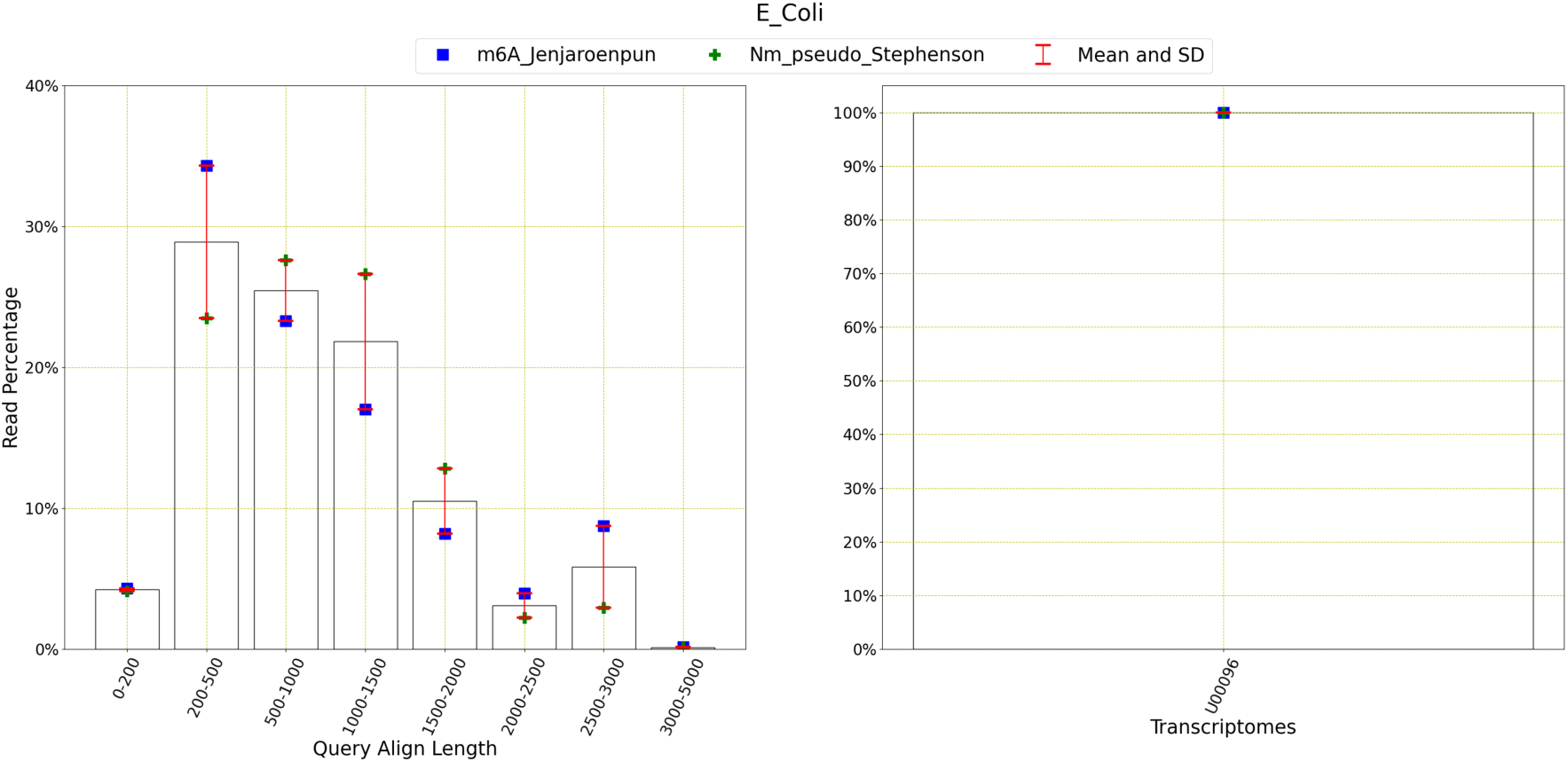
Read distribution of E Coli data: This figure shows the distribution of the reads for query align length, mapping quality and transcriptomes. The average number of reads with query length under 200 is 173962 and over 200 is 4003333. This means 95.84% of the reads have a query length of greater than 200. The average number of reads of mapping quality less than 20 is 4058735 and over 20 is 123296. This shows that 2.95% of reads have a mapping quality higher than 20.

On average, reads from both sources have the query align length of 200 to 1500. Majority of the mapping qualities lie in between 0 and 5, although some contain a higher mapping quality of 60 to 80. There is only one transcriptome.

### 4.8. Arabidopsis Thaliana

This is already been explained in Section 3.1.7 as there is only one dataset containing this reference sequence.

## 5. Datasets from various sources

We have used seven different sources to gather our data. The analysis based on those sources are described here.

### 5.1. Pratanwanich et al

As Pratanwanich et al. [20] contains only the human reference on the m6A, the following figure is very similar to the previous **Figure 3**.

I need to double check the values mentioned in the flowing figures.

The above tables shows all the datasets mentioned in Pratanwanich et al. follow the same trend found in **Figure 3**.

**Figure 38:**
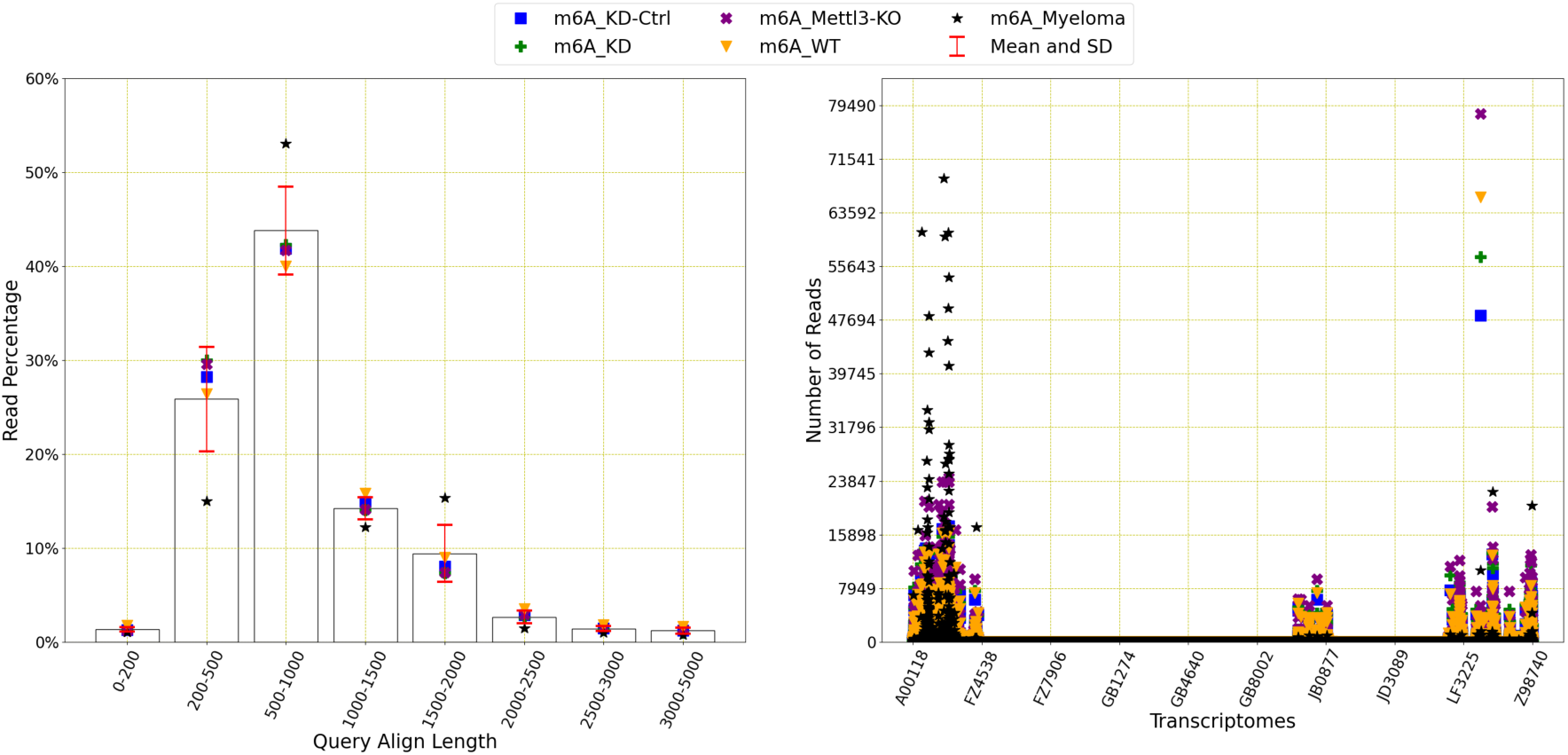
R**e**ad **distribution of Pratanwanich et al. data:** This figure shows the distribution of the reads for query align length, and transcriptomes. The average number of reads with query length under 200 is 85181 and over 200 is 6028257. This means 98.61% of the reads have a query length of greater than 200. The average number of reads of mapping quality less than 20 is 6088874 and over 20 is 110630. This shows that 1.78% of reads have a mapping quality higher than 20.

### 5.2. Mateos et al

Mateos et al. [27] contains only two reference sequences, human and mouse for m6A methylation. The following figure explains the query align length and mapping quality.

**Figure 39.**
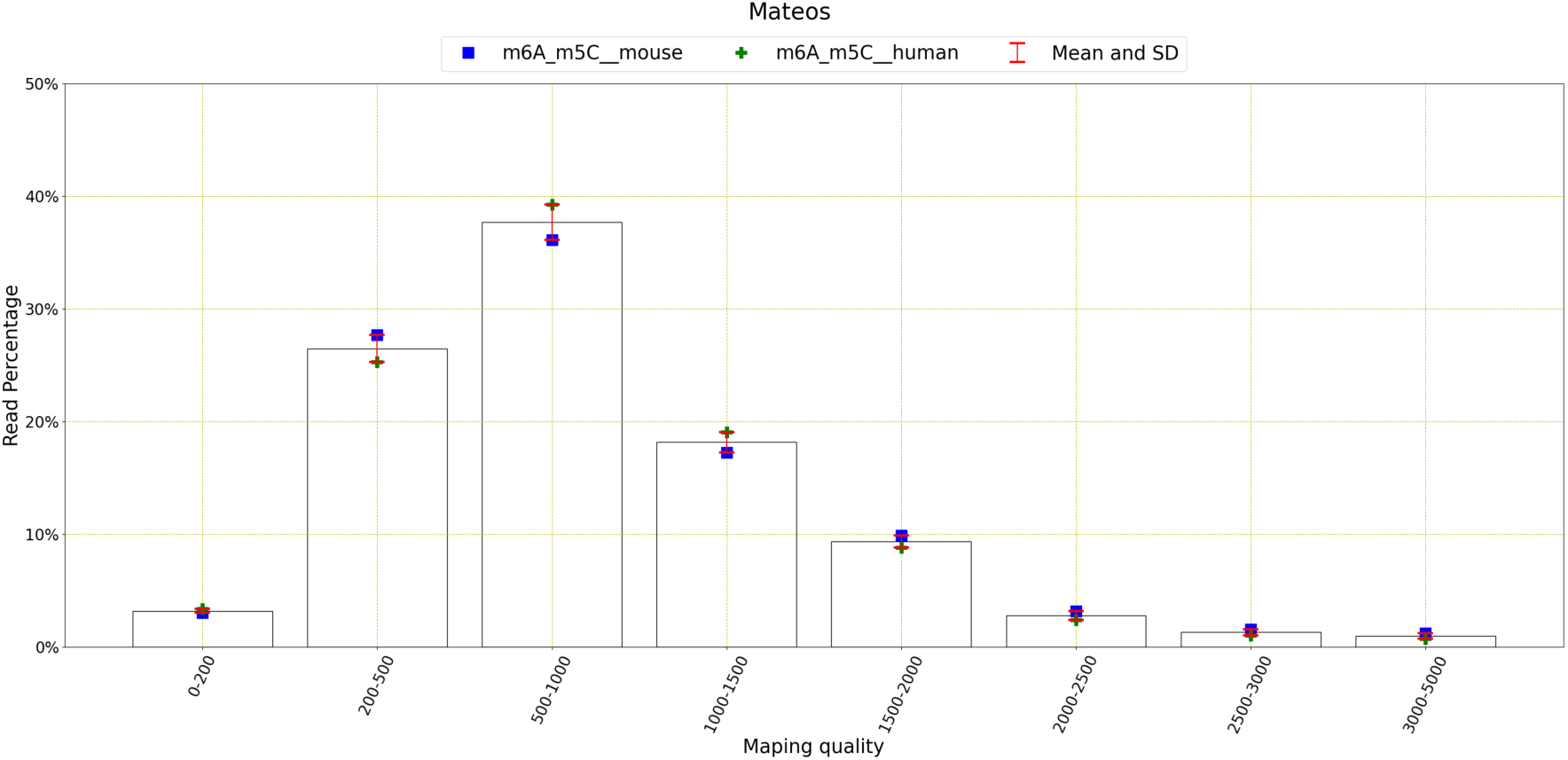
Read distribution of Mateos et al. data: This figure shows the distribution of the reads for query align length, mapping quality and transcriptomes. The average number of reads with query length under 200 is 77956 and over 200 is 2253985. This means 96.66% of the reads have a query length of greater than 200. The average number of reads of mapping quality less than 20 is 2308568 and over 20 is 72577. This shows that 3.05% of reads have a mapping quality higher than 20.

The figure shows that most of the query lengths lie in between 500 and 1000. Query lengths for both mouse and human reference sequence do not vary a lot from each other. Also, most of the mapping qualities are in between 0 to 5 for both datasets.

### 5.3. Liu et al

In Liu et al. [24], there are two reference sequences, both used for m6A methylation. The following figure shows that most of the query lengths lie between 200 and 500 for yeast, and 500 and 1000 for curlcake ([24]) sequence. The query lengths vary a lot between these two datasets.

The mapping quality lie between 60 and 80 for most reads for both datasets.

**Figure 40.**
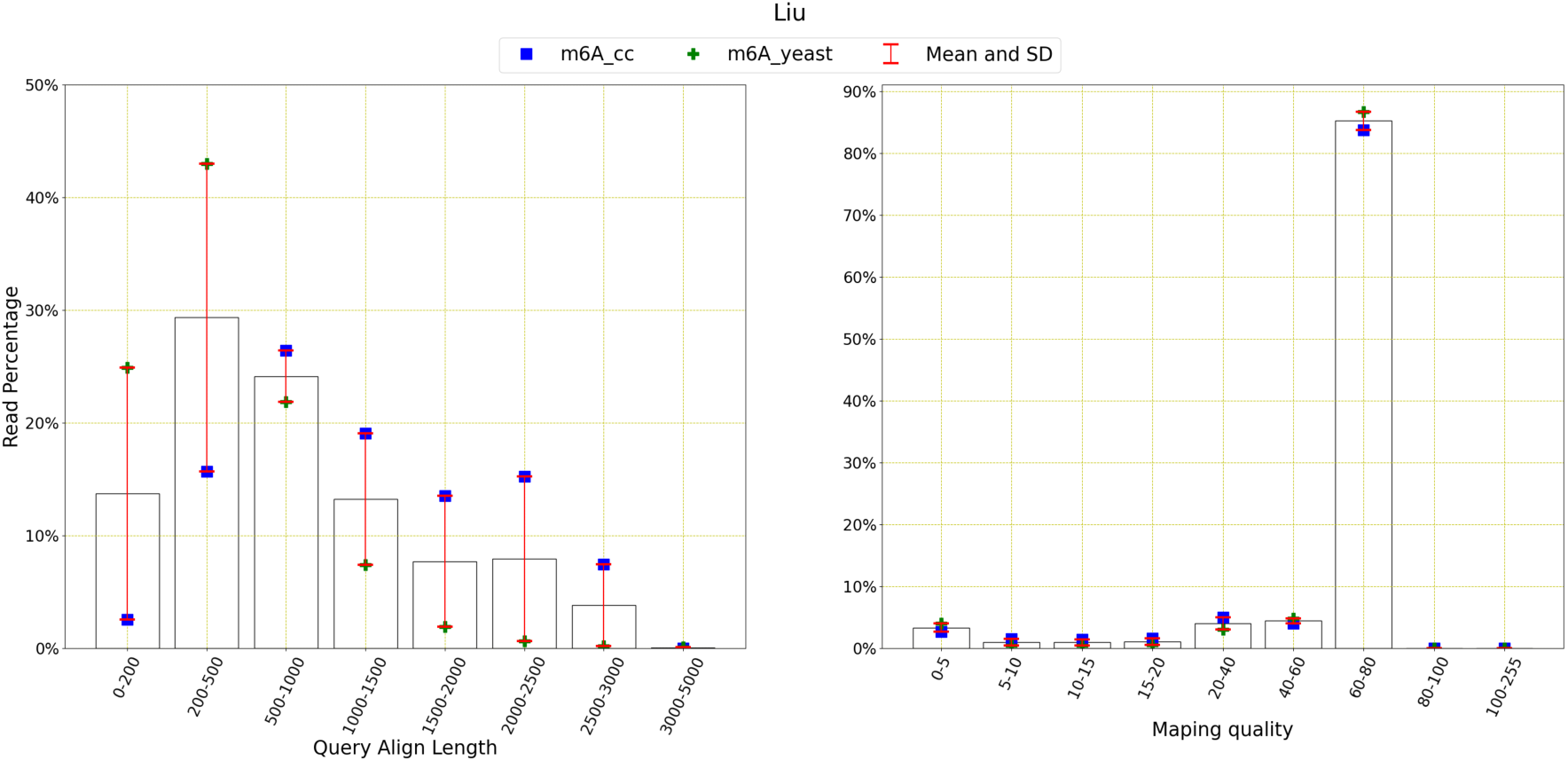
Read distribution of Liu et al. data: This figure shows the distribution of the reads for query align length, mapping quality and transcriptomes. The average number of reads with query length under 200 is 60995 and over 200 is 291443. This means 82.69% of the reads have a query length of greater than 200. The average number of reads of mapping quality less than 20 is 22096 and over 20 is 340903. This shows that 93.91% of reads have a mapping quality higher than 20.

### 5.4. Begik et al

Begik et al. [9] contains two reference sequences, such as curlcake ([9]) and yeast and three different methylations such as pseudo, 5hmC and m5C. They are explained in the figure below.

The above figure shows that four different datasets have query lengths varying from 0 to 3000 and most of the reads have the query length of 500 to 1000 on average. Even though some reads do contain mapping quality between 0 and 5, most of the reads have a higher mapping quality of 60 to 80.

### 5.5. Jenjaroenpun et al

Jenjaroenpun et al. [28] is dedicated to m6A methylation only, but contains E Coli, human, IVT and mouse reference sequences, as shown in the flowing figure.

The figure shows that most of the reads on average have the query length of 200 to 500, however it varies a lot among different reference sequences. The reads mainly fall under the mapping quality of 0 to 5, however a significant portion of reads also contain the mapping quality of 60 to 80.

### 5.6. Zhong et al

A similar trend is also noticed in Zhong et al. [10] as it contains m6A methylation only for two reference sequences such as mouse and Arabidopsis. The read distribution is shown below.

**Figure 41.**
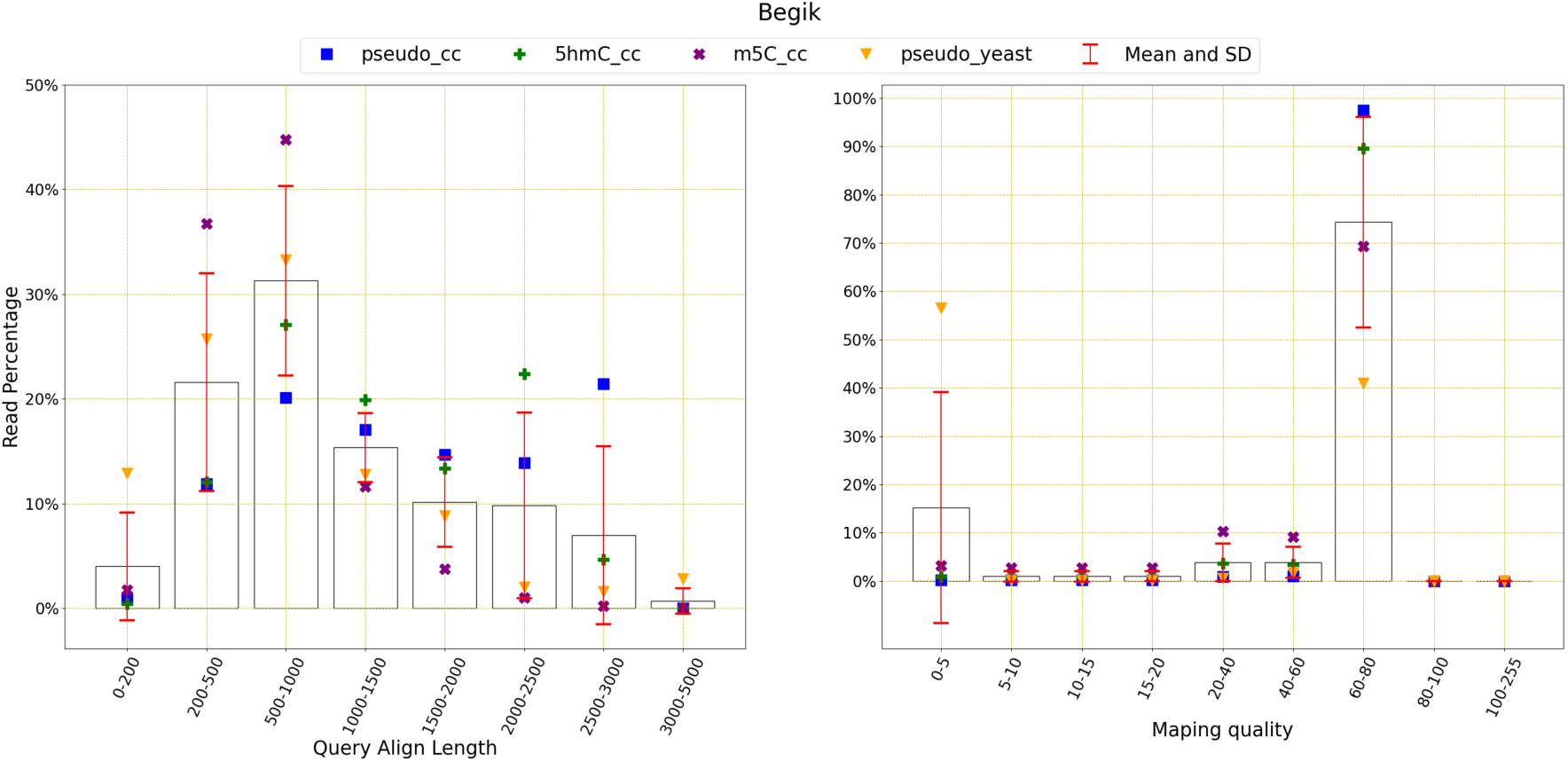
Read distribution of Begik et al. data: This figure shows the distribution of the reads for query align length, mapping quality and transcriptomes. The average number of reads with query length under 200 is 30077 and over 200 is 280388. This means 90.31% of the reads have a query length of greater than 200. The average number of reads of mapping quality less than 20 is 133442 and over 20 is 195339. This shows that 59.41% of reads have a mapping quality higher than 20.

**Figure 42.**
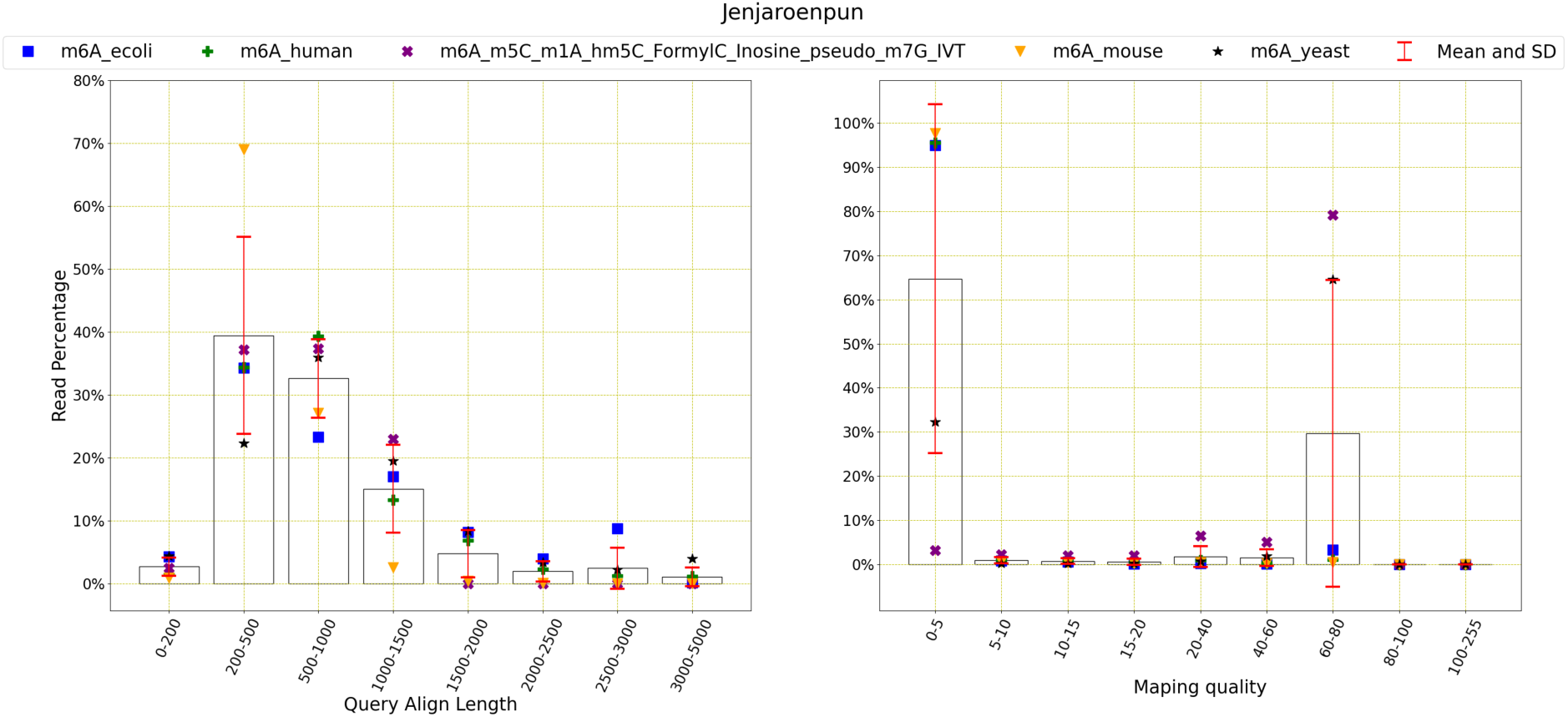
Read distribution of Jenjaroenpun et al. data: This figure shows the distribution of the reads for query align length, mapping quality and transcriptomes. The average number of reads with query length under 200 is 70252 and over 200 is 4230206. This means 98.37% of the reads have a query length of greater than 200. The average number of reads of mapping quality less than 20 is 4026253 and over 20 is 368068. This shows that 8.38% of reads have a mapping quality higher than 20.

The figure shows that most reads have the query length of 500 to 1000 for both datasets. The mapping quality also lie in two extremes, such as 0 to 5 and 60 to 80.

### 5.7. Stephenson et al

Stephenson et al. [29] contains only Nm and pseudo types of methylation for e coli, yeast and human reference sequences. Their read distribution is shown below.

**Figure 43.**
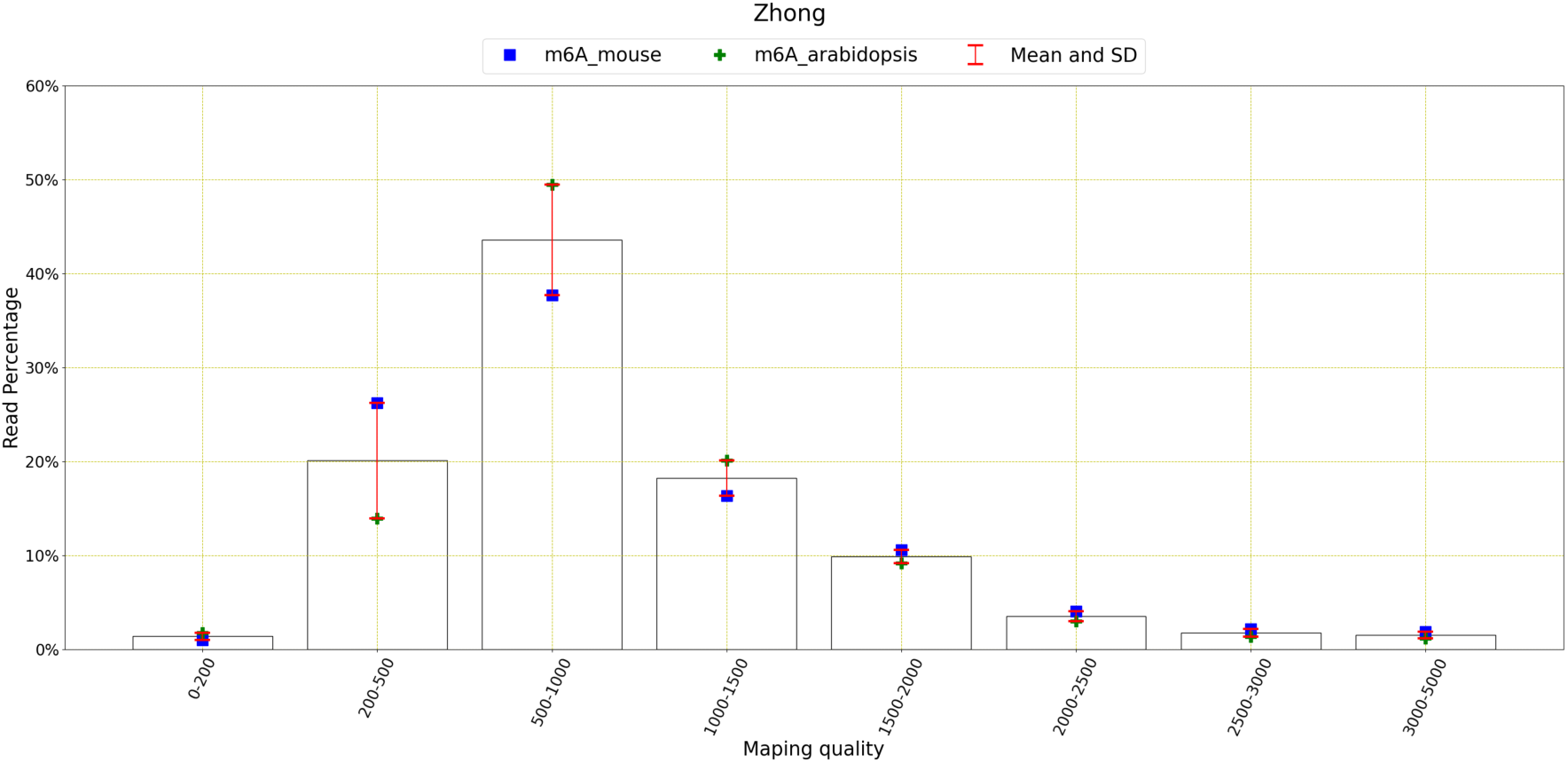
Read distribution of Zhong et al. data: This figure shows the distribution of the reads for query align length, mapping quality and transcriptomes. The average number of reads with query length under 200 is 42923 and over 200 is 3756274. This means 98.87% of the reads have a query length of greater than 200. The average number of reads of mapping quality less than 20 is 3445787 and over 20 is 510415. This shows that 12.90% of reads have a mapping quality higher than 20.

**Figure 44.**
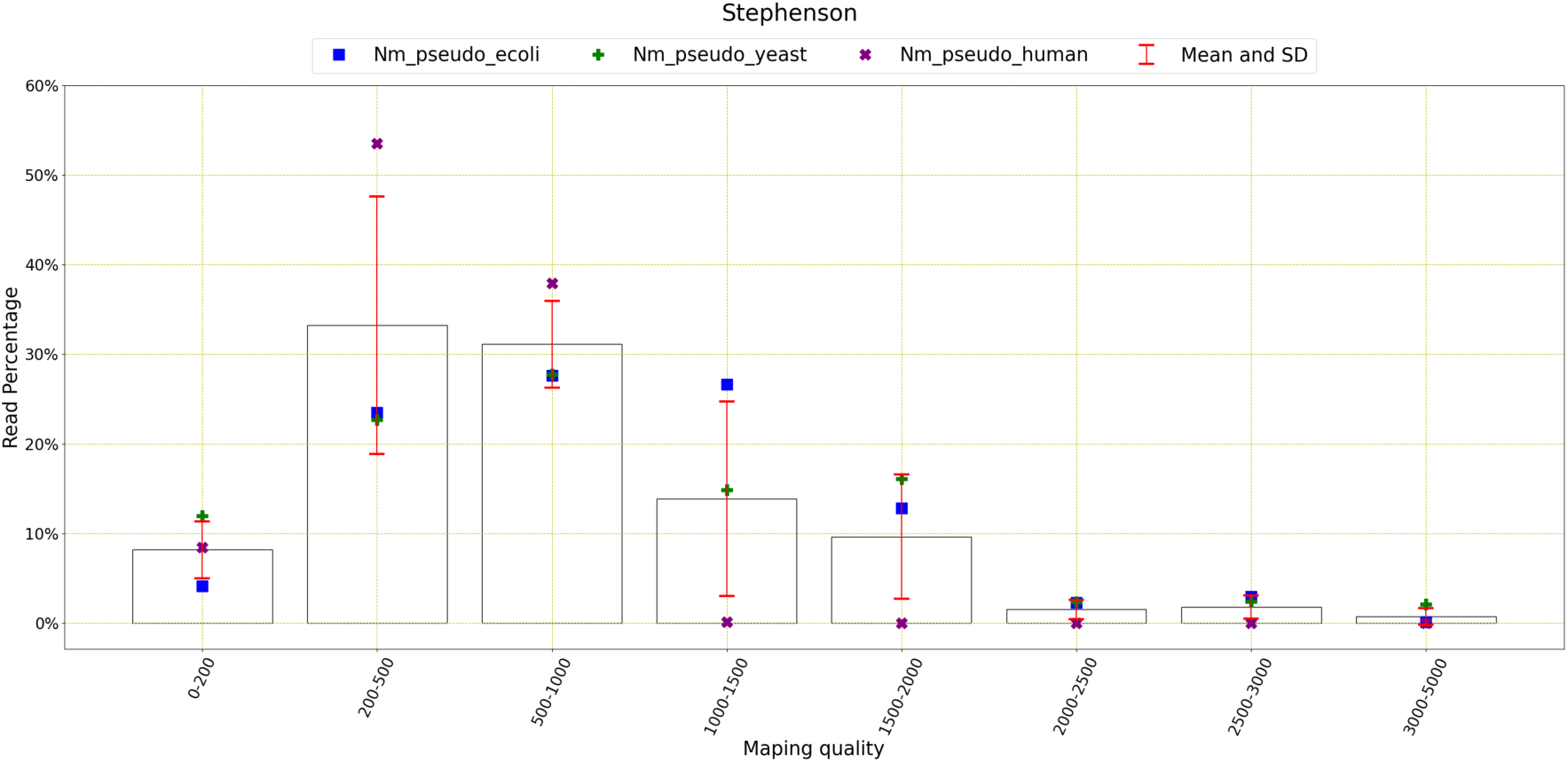
Read distribution of Stephenson et al. data: This figure shows the distribution of the reads for query align length, mapping quality and transcriptomes. The average number of reads with query length under 200 is 253,172 and over 200 is 3,498,470. This means 93.25% of the reads have a query length of greater than 200. The average number of reads of mapping quality less than 20 is 3,768,581 and over 20 is 89,055. This shows that 2.31% of reads have a mapping quality higher than 20.

The above figure shows that most of the reads have a query align length of 200 to 1000 on average, specially the human dataset. Almost all reads have the mapping quality of 0 to 5.

## 6. Conclusions

In this study, we have considered Oxford Nanopore sequencing data from seven different sources, and they include eight different reference transcriptomes such as human, mouse, curlcakes, In vitro test (IVT), yeast, E Coli and Arabidopsis Thaliana. The datasets are large, several gigabytes of size. We have organized the data according to various types of methylation, transcriptomes and sources. Nine different types of methylations are considered, and they are m6A, m5C, Nm, pseudouridine, hm5C, FormylC, Inosine, m1A and m7G. We find that methylated data has lower base-mapping success rate, Tombo success rates and number of pass-reads. Most the datasets represented m6A methylations as this is the most popular type of methylations and had the highest success rate among the other methylation types.

In our future work, we will generate our own ONT data and compare it with the existing ones. We have developed an automatic CNN-based m6A methylation detection system and later we will extend the application to the other types of methylations mentioned in this paper.

